# scRefine: hierarchical and contrastive learning for biologically faithful single-cell atlas integration and annotation

**DOI:** 10.64898/2026.11.25.769099

**Authors:** Sven Hauns, Costerwell Khyriem, Ankita Singh, Yousef A. Elkenawi, Rasheed Mohammad, Aditya Sharma, Imran Khan, Nelson C. Soares, Babacar Cisse, Fahad Ali, Ahmad Abou Tayoun, Mohammed Uddin, Rolf Backofen, Omer S. Alkhnbashi

## Abstract

Single-cell RNA sequencing (scRNA-seq) studies increasingly encompass diverse tissues, donors, disease contexts, and sequencing platforms, resulting in large-scale datasets characterized by substantial technical heterogeneity and inconsistent cell-type annotations. These datasets offer unprecedented opportunities to resolve fine-grained cellular diversity and identify tissue-specific, developmental, and disease-associated cell states. However, existing integration and annotation approaches often struggle to preserve rare or transcriptionally subtle subpopulations, depend heavily on extensively labelled reference datasets or predefined cell-type taxonomies, and can generate latent representations that obscure biologically meaningful lineage relationships and fine-scale cellular structure.

We present scRefine, an annotation-aware contrastive learning framework for integrating heterogeneous single-cell RNA-seq datasets while refining cell identities from partially labelled data. scRefine learns a biologically structured latent space, enabling improved resolution of rare and transcriptionally similar populations. In particular, scRefine better separates rare or biologically distinct populations that exhibit convergent or overlapping translational states than other integration methods. For example, scRefine cleanly separates SPP1+ melanocytic tumor cells from SPP1+ tumor-associated macrophages (TAMs), indicating that it preserves lineage-specific biological identity despite shared tumor-associated programs. Beyond cell-state resolution, scRefine also preserves continuous biological structure, as demonstrated by its ability to recapitulate the developmental trajectory of the frontal brain in a manner consistent with the patterns reported in the original study. scRefine also calculates a confidence score by estimating cluster membership using nearest neighbour computations for each cell, which aids in defining cells with ambiguous or mixed transcriptional signatures.

Across multi-tissue immune atlases spanning 92 fine-grained classes, scRefine consistently outperforms state-of-the-art methods in accuracy, rare population recovery, and generalization to unseen datasets. scRefine also maintains high annotation fidelity even when trained with only 1–5% labelled cells, greatly reducing the training effort and increasing the generalizability. scRefine is available as an open-source package and offers a robust foundation for constructing coherent, high-resolution cell-type reference atlases across studies and technologies.

## Introduction

Single-cell RNA sequencing (scRNA-seq) has become a central approach in biological research, providing insight into cellular diversity across tissues and disease states. In contrast to bulk transcriptomics, which averages gene expression signals across heterogeneous cell populations within a biological sample, scRNA-seq captures transcriptomic profiles at single-cell resolution across diverse sample sources, including primary tissues, biopsies, blood, organoids, and cultured cell populations [1–4]. This capability enables the identification of rare cell types, transient cellular states, and dynamic lineage transitions [1–4]. Such resolution is particularly important in immunology, oncology, and developmental biology, where the cellular microenvironment shapes pathophysiological processes such as immune evasion, tumor progression, and organogenesis [6–9]. However, the analysis of scRNA-seq data presents substantial challenges, particularly in data integration, cell-type annotation, and the harmonization of independently generated datasets [10–14]. The rapid expansion of scRNA-seq datasets across diverse platforms, biological conditions, and laboratories introduces substantial technical variability and inconsistencies in cell-type annotation, hindering robust cross-study comparisons and the construction of unified reference atlases. Accurate and consistent resolution of cell identities across heterogeneous datasets is therefore essential for deriving biologically meaningful, reproducible, and broadly comparable insights from large-scale single-cell studies [15–16].

To tackle these challenges, a range of computational approaches have been developed to address cell-type annotation and cross dataset integration. For automated annotation, CellTypist uses reference-trained machine-learning models to provide rapid and scalable cell-type prediction, with particular strength in resolving immune populations across tissues [17]. However, predictions are constrained by the cell identities represented in the training model, making previously unrepresented populations or differences in annotation granularity more difficult to resolve. CellHint addresses a related problem by harmonizing independently generated annotations using predictive clustering trees and transcriptomic relationships between cell types. This enables reconciliation of differing annotation resolutions and identification of shared, dataset-specific, or potentially novel populations across studies [18]. Its performance, however, necessarily depends on the availability and quality of the initial annotations used to establish these relationships.

In parallel, several methods focus primarily on removing technical variation during dataset integration. Harmony iteratively adjusts low-dimensional representations to achieve efficient integration whereas Scanorama identifies shared cellular populations between datasets and aligns them with a common representation [19,20]. Both approaches are computationally efficient and broadly applicable, but neither explicitly incorporates prior knowledge of cellular identity during representation learning. scVI instead uses a probabilistic deep generative model to jointly represent biological and technical variation, providing a scalable latent representation that supports integration and several downstream analyses [21]. However, the standard scVI framework is unsupervised with respect to known cell identities and marker programs.

More recent methods aim to improve preservation of biological structure during integration. CellANOVA explicitly models unwanted technical variation using a pool-of-controls experimental design, allowing biological signals that may otherwise be removed during batch correction to be recovered [22], although its application depends on an appropriate experimental design and suitable control samples. Contrastive-learning approaches provide an alternative strategy. scCM uses self-supervised contrastive learning to organize transcriptionally related cells while separating dissimilar populations and has demonstrated effective integration of large-scale CNS datasets [23]. CONCORD similarly employs self-supervised contrastive learning with dataset-aware and hard-negative sampling to mitigate technical variation while preserving the underlying cell-state landscape [24]. These approaches learn cellular structure directly from transcriptional similarity but do not explicitly use established marker-gene knowledge to define cellular identity before integration.

To address this gap, we developed scRefine, which couples biologically informed cell classification with contrastive representation learning. Rather than integrating cells solely according to their complete transcriptional similarity, scRefine also focuses on established marker-gene programs to infer cellular identities and organize cells within a hierarchical biological framework. These inferred identities subsequently guide integration, encouraging transcriptionally and biologically related cells to converge while preserving distinctions between lineages and fine-grained cellular states. By linking marker-informed embedding, annotation-aware integration, and cell-level confidence estimation within a unified framework, scRefine is designed to harmonize heterogeneous datasets while retaining biologically meaningful cellular heterogeneity. We benchmarked its performance using a cancer liver-metastasis atlas, a metastatic breast cancer dataset, a global immune atlas, and a frontal brian development dataset, demonstrating improved annotation accuracy, effective batch correction, and biologically meaningful representations. Across these datasets, scRefine demonstrated consistently strong performance in cell-type annotation, dataset integration, and preservation of biological structure compared with CellTypist [17], scVI [25], CellANOVA [26], Harmony [27], scANVI [28], scCM [29], Scanorama [30], and CONCORD [31]. Overall, these results demonstrate the ability of scRefine to integrate diverse single-cell datasets while retaining biologically meaningful cellular variation.

## Results

### scRefine framework

Previous analyses of single-cell data indicate that gene expression across cell types is significantly constrained by cell-specific gene programs, often characterised by the covariation of specific marker genes [32–34]. Regularisation [33,34] enables the establishment of lower-dimensional embeddings that explain and influence the high-dimensional gene-cell matrix. These embeddings can be informed by established expert insights into the significance of particular gene programs.

To capture structures that reflect known variations in gene programs, scRefine integrates unsupervised embedding generation with marker-gene-guided conditioning. The unsupervised embedding is formed by adhering to the structure of multi-dataset integration and by segregating priors into hierarchically dependent biological and technological factors [35,36]. This approach enables scRefine to model independent batch-specific effects while accounting for their relationship with biological priors. To estimate the quality of the reconstruction a Zero-Inflated Negative Binomial (ZINB) loss function is used [37,38].

To further enhance the embeddings, scRefine implements two variants to condition them with marker genes: (i) the extent of marker gene expression is binned in a data-specific manner and then predicted using classifiers trained on the biological embedding; and (ii) the latent space is directly conditioned via contrastive learning based on the cosine similarity of marker gene based samples. This requires scRefine to learn to cluster independently of dataset-specific similarities.

To further refine cluster analysis, scRefine utilises confidence scores derived from radius-based nearest-neighbour predictions. Cells with lower confidence are systematically removed according to a predefined cutoff until a clearer delineation of high-density clusters is achieved.

### scRefine integrates heterogeneous single-cell atlases while preserving biological topology

Single-cell gene-expression profiles are not randomly distributed. Instead, they are constrained by coordinated regulatory programs that generate an organised transcriptional landscape in low-dimensional space. Within this landscape, compact clusters generally correspond to relatively stable cell identities, intermediate-density regions may represent transient or mixed states, and continuous or branching geometries can reflect differentiation, activation, or pathology-associated transitions. Recovering these structures from large-scale single-cell atlases requires not only effective batch correction but also preservation of the underlying geometry and topology of the gene-expression space.

In large multi-study atlases, however, biologically meaningful structures are frequently obscured by dataset-specific technical variation arising from differences in experimental protocols, sequencing chemistry, tissue-dissociation procedures, sample handling, human error, and other study-specific biases [39–43]. Consequently, low-dimensional embeddings may separate cells predominantly according to their dataset or study of origin rather than their underlying biological identity or state. Inadequate correction therefore leaves technical effects embedded within the representation, whereas excessive correction can remove genuine biological variation by forcing transcriptionally distinct populations to mix. We designed scRefine to address this challenge by integrating highly heterogeneous single-cell, multi-tissue atlases while preserving the intrinsic organisation of the transcriptional landscape and preventing dataset-specific batch effects from dominating the learned representation (Figure 1).

**Figure 1:**
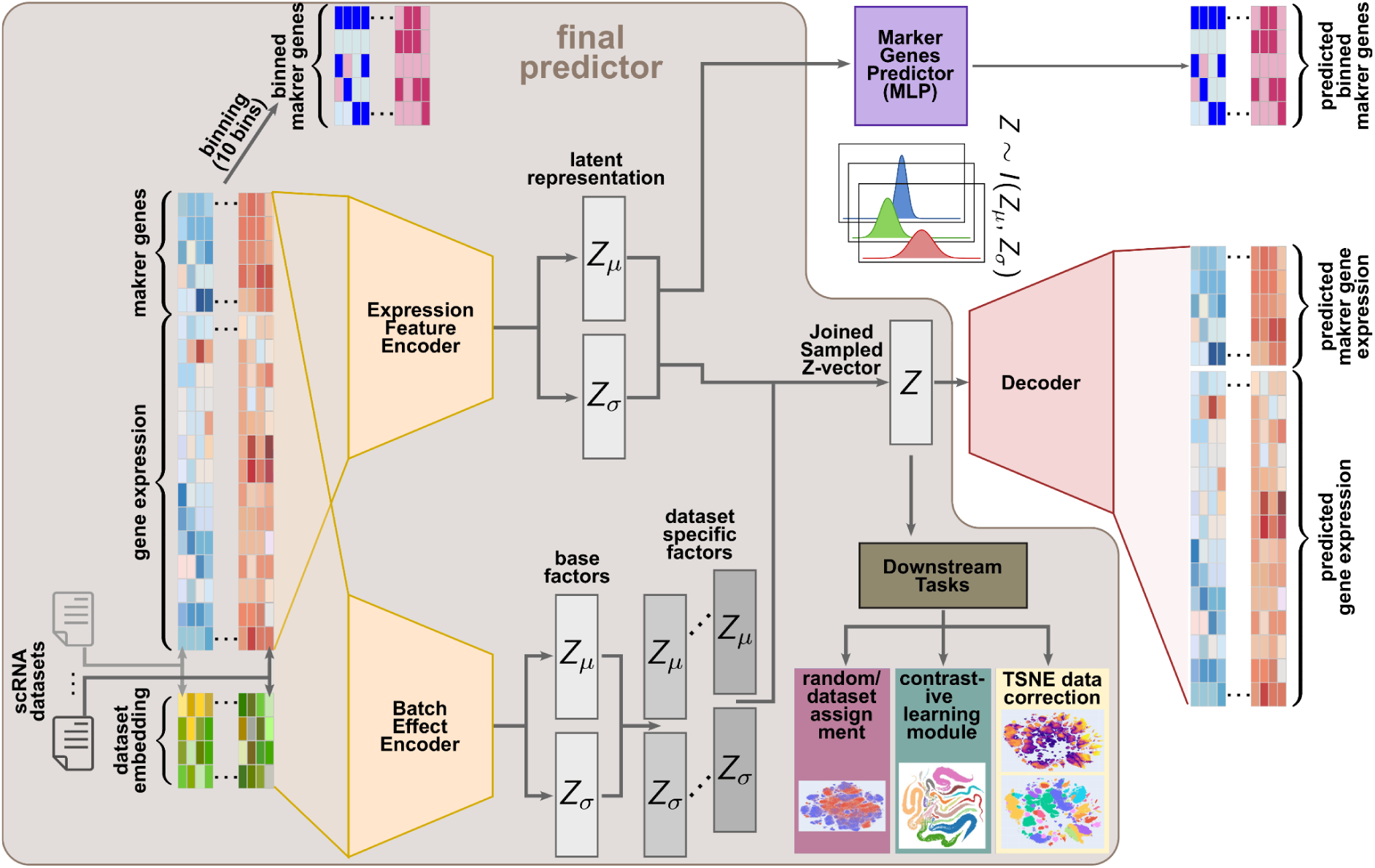
Overview of the scRefine framework for biologically informed integration of scRNA datasets. The scRefine architecture integrates heterogeneous scRNA datasets using a hierarchical variational autoencoder (HVAE) augmented with biological supervision and dataset-aware latent factors. Gene expression matrices from multiple datasets are first processed together with dataset embeddings that encode the origin of each cell. Marker genes associated with key biological programs are additionally incorporated and discretized into expression bins to guide the model toward biologically meaningful representations. Two complementary encoders capture different sources of variation. The expression feature encoder learns a latent representation of biological variation from gene expression profiles, producing mean and variance parameters (*Z*μ, *Z*σ) that describe the biological latent distribution. In parallel, a batch-effect encoder learns dataset-specific latent variables that model technical variation and library-size effects across input datasets. These latent variables are combined to generate a joint sampled latent representation (*Z*), which serves as the unified embedding space for downstream analysis. The shared latent representation is used in two ways. First, it is passed through a decoder network that reconstructs gene expression using a zero-inflated negative binomial model, enabling the model to capture dropout and over-dispersion typical of single-cell data. Second, the latent embedding is connected to a marker-gene prediction head, implemented as a multilayer perceptron (MLP), which predicts binned expression levels of selected marker genes. This auxiliary prediction task constrains the latent space to preserve biologically meaningful transcriptional programs. Alternatively the model makes use of a contrastive module using the cosine similarity of samples to further condition the latent space. The model is trained using a joint objective combining reconstruction loss, Kullback–Leibler divergence regularization, and marker-gene prediction loss. Once trained, the learned embedding supports multiple downstream analyses, including dataset harmonization through random dataset assignment, contrastive representation learning for improved cell-type discrimination, and visualization or data correction methods such as t-SNE-based embedding refinement. Together, these components enable scRefine to disentangle dataset-specific technical effects from shared biological variation, producing a unified latent representation suitable for atlas-scale single-cell integration and high-resolution cell-state analysis.

To evaluate this capability, we assembled a multi-cancer liver-metastasis single-cell atlas, hereafter referred to as the LiverMet atlas. The atlas comprised primary and metastatic tumour biopsies from uveal melanoma, colorectal, pancreatic, neuroendocrine, melanoma, thyroid, and breast cancer cohorts, together with normal liver samples incorporated from an independent liver atlas (Supplementary Figure S1; Table S1). This collection represents a stringent integration challenge because it combines samples generated across multiple studies, tissues, cancer types, and experimental protocols. It also contains substantial variation in cellular composition, ambient RNA contamination, tumour state, and overlapping but non-identical biological programs across cohorts [44–47].

The inclusion of an independent normal liver reference further enabled us to investigate whether scRefine could harmonise datasets while retaining biologically meaningful hepatocyte heterogeneity, including stress-associated hepatocyte populations enriched in metastatic samples [48–50]. To determine which information was captured directly by the model, scRefine was trained from scratch on the LiverMet atlas. The resulting latent representation was subsequently projected into two dimensions using t-SNE for visualisation.

The scRefine embedding displayed a well-organised transcriptional landscape in which cells formed compact and interpretable neighbourhoods corresponding to major immune, epithelial, stromal, endothelial, melanocytic, neuroendocrine, and hepatocyte compartments (Figure 2A; Supplementary Figure S2). Related cell populations occupied adjacent regions, whereas transcriptionally distinct lineages remained spatially separated. Importantly, the representation did not collapse into a uniformly mixed or amorphous cloud. Instead, scRefine preserved regional organisation and local neighbourhood structure while integrating cells across studies.

**Figure 2:**
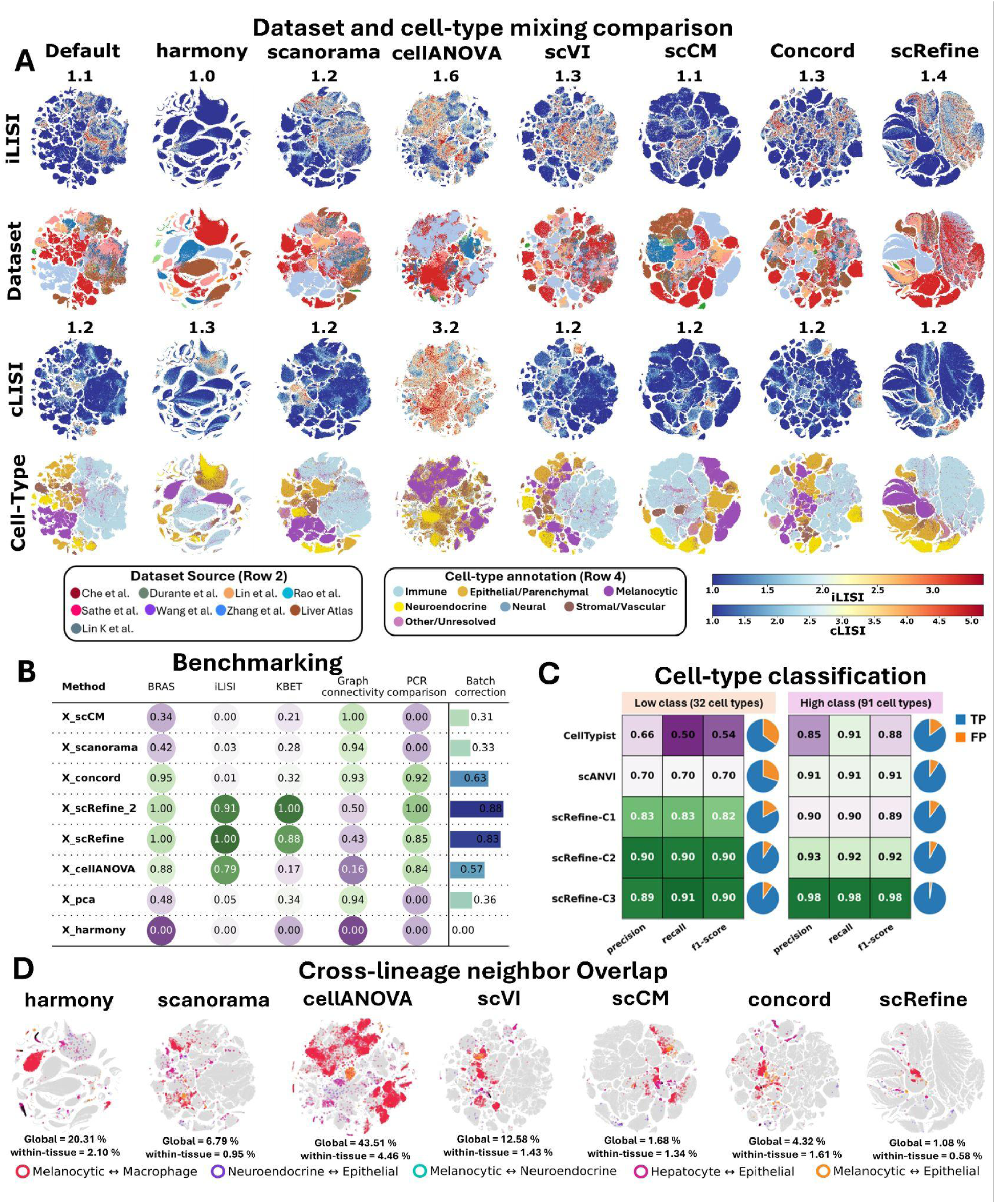
Comparative evaluation of tissue integration, cell-type preservation, primary–metastatic alignment, and cell-type classification. (**A)** Method-specific t-SNE representations of the integrated atlas. Rows show standardized integration iLISI, tissue identity, standardized tissue iLISI calculated within cell types, and cell-type annotation (pan-human azimuth annotation). Higher tissue iLISI indicates stronger mixing across data sets, whereas within-cell-type tissue iLISI assesses whether such mixing is retained after controlling for cellular identity. Cells are colored by the iLISI score per cell across integration methods. Notably, melanocytic tumor cells localize predominantly to regions occupied by primary and metastatic uveal melanoma samples (UM and mUM, see row 2), consistent with their expected tissue-of-origin specificity. This correspondence remains visible after integration, indicating that alignment of UM with mUM does not eliminate the melanocytic lineage signal (row 4). The clearer co-localization in scRefine suggests improved primary–metastatic alignment within this lineage while retaining its separation from unrelated epithelial and stromal populations. **(B)** Quantitative comparison of integration performance using BRAS, iLISI, kBET, graph connectivity, PCR comparison, and the aggregate batch-correction score. **(C)** Cell-type classification performance in low- and high-class-number settings. Precision, recall, and F1 score are reported together with the proportions of true-positive and false-positive assignments. C1–C3 denote the evaluated scRefine classifier configurations. Results for all 92 cell types in the high-resolution category are shown in Supplementary Figure S4. **(D)** Cross-lineage neighborhood overlap reveals differences in the preservation of cellular identity across integration methods. Comparative visualization of cross-lineage nearest-neighbor overlap across seven single-cell transcriptomic integration approaches: Harmony, Scanorama, cellANOVA, scVI, scCM, Concord, and scRefine. Each panel displays the corresponding two-dimensional embedding of the integrated cancer atlas, with the overall cellular landscape shown in light gray and cells exhibiting cross-lineage neighborhood overlap highlighted according to the participating cell-type pairs. Five biologically distinct lineage combinations are examined: melanocytic–macrophage (red), neuroendocrine–epithelial (purple), melanocytic–neuroendocrine (turquoise), hepatocyte–epithelial (magenta), and melanocytic–epithelial (orange). These comparisons assess the extent to which transcriptionally distinct cellular lineages occupy overlapping neighborhoods in the integrated representation, providing an additional measure of cellular identity preservation and potential overintegration. Two complementary overlap measurements are reported beneath each embedding: global cross-lineage neighborhood overlap, calculated across the integrated atlas, and within-tissue overlap, evaluating cross-lineage neighborhood relationships within individual tissue contexts. Substantial differences in cross-lineage overlap are observed among the integration methods. cellANOVA exhibits the highest global overlap (43.51%) and within-tissue overlap (4.46%), with widespread highlighted populations distributed across multiple regions of the embedding, suggesting extensive mixing between distinct cellular lineages. Harmony also demonstrates considerable global overlap (20.31%), with a within-tissue overlap of 2.10%, followed by scVI (12.58% and 1.43%) and Scanorama (6.79% and 0.95%). Concord and scCM show substantially reduced global overlap, reaching 4.32% and 1.68%, respectively, although their within-tissue overlap remains at 1.61% and 1.34%. Notably, **scRefine achieves the lowest global (1.08%) and within-tissue (0.58%) cross-lineage overlap**, with highlighted cells confined to relatively restricted regions of the embedding. These findings suggest that scRefine more effectively maintains separation between the evaluated major cellular lineages while limiting inappropriate cross-lineage neighborhood mixing. The consistently lower within-tissue overlap percentages compared with global values further emphasize the importance of accounting for tissue origin when evaluating cell-type segregation across heterogeneous cancer datasets. Although reduced cross-lineage overlap is consistent with improved preservation of distinct cellular identities, some overlap may reflect genuine biological similarities, transitional cell states, or shared transcriptional programs rather than integration artifacts. Together, these analyses demonstrate pronounced method-dependent differences in cross-lineage neighborhood organization and support the ability of scRefine to preserve major lineage boundaries within the integrated primary and metastatic cancer atlas.

We next compared scRefine with representative integration methods, including cellANOVA, CONCORD, Harmony, Scanorama, scCM, scVI, and a no-integration baseline. Examination of tissue origin and source dataset revealed pronounced tissue- and study-associated segregation in the uncorrected embedding, demonstrating that technical variation strongly influenced the original expression space (Figure 2A, rows 1–2; Supplementary Figure S2 and S3). This segregation persisted to different degrees after integration with Harmony, Scanorama, scCM, and CONCORD. Harmony substantially altered the global organisation of the embedding, generating prominent ring-like or striped structures, while producing comparatively limited improvement in dataset-level mixing. Scanorama, scCM, CONCORD, and scVI reduced dataset separation to varying extents but retained residual study-associated structures or fragmentation of biologically related populations.

scVI showed modestly greater cross-dataset overlap than several of the conventional integration approaches, whereas cellANOVA and scRefine produced the strongest overall dataset and tissue mixing. Quantitative assessment using the integrative LISI ( iLISI), a score that uses the inverse Local Simpson’s Index (LISI) to assess the mixing of different datasets in a local neighborhood, supported this observation: scRefine achieved the second highest dataset-level mixing (mean iLISI = 1.4), following cellANOVA (1.6) (Supplementary Figure S2), while both methods achieved the highest LISI score for Tissue and Tumor/Normal level mixing (Figure 2A). However, the embeddings generated by these methods differed markedly in their biological organisation. In scRefine, cells from different datasets were integrated within coherent lineage-specific compartments, and transcriptionally related populations occupied neighbouring regions. By comparison, the strong mixing achieved by cellANOVA was accompanied by poorer preservation of clearly delineated cell-type structure (Figure 2A). These observations highlight an important distinction between maximising batch mixing and achieving biologically faithful integration (Figure 2A).

Analysis of normal and tumour-derived cells provided an additional measure of biological preservation (Supplementary Figure S2). Despite achieving strong dataset and tissue mixing, scRefine showed relatively limited mixing between normal and tumour-derived cells (mean tumour/normal iLISI = 0.33). In comparison, cellANOVA exhibited greater tumour–normal mixing (0.43), indicating that its stronger global harmonisation was accompanied by increased mixing across this biological axis. In contrast, scRefine aligned cells across datasets within shared cellular compartments while retaining localised regions enriched for normal or tumour-derived populations. Normal liver cells therefore remained spatially coherent within an otherwise predominantly tumour-associated landscape. Together, these patterns indicate that scRefine reduces dataset- and tissue-associated variation while retaining major transcriptional differences associated with disease state.

To determine whether the improved harmonisation was achieved without compromising cell identity, cell-type annotations were generated independently using Pan-human azimuth [51] and projected onto each embedding (Figure 2A, row 4). Because these annotations were generated externally, they provided a reference that reduced potential circularity arising from the marker-gene information used during scRefine contrastive learning.

In the scRefine embedding, independently annotated cell populations formed coherent compartments, with related cell types occupying neighbouring regions while distinct lineages remained separated. Dataset mixing was subsequently quantified within these independently defined cell-type groups (Figure 2A, row 5). scRefine achieved the second highest dataset-mixing index of 1.4, following cellANOVA at 1.6 while scVI and concord follow suit with both at 1.3. Importantly, the similar global mixing achieved by scRefine and cellANOVA did not translate into equivalent biological organisation. High mixing observed for cellANOVA was accompanied by reduced coherence of the azimuth-defined populations, whereas scRefine combined strong within-cell-type cross-dataset mixing with clearer preservation of cell-type-associated neighbourhoods. Thus, dataset mixing alone was insufficient to determine integration quality: an effective representation must integrate equivalent populations across datasets while maintaining the boundaries and relationships that distinguish biologically different cell states. It should be noted that cellANOVA was developed specifically for comparative analyses of diseased and normal tissues. In our integration framework, the normal liver atlas was explicitly used as the normal reference, which may have contributed to stronger correction of disease-associated variation and, consequently, to the apparent over-correction observed for cellANOVA.

To further assess whether scRefine preserves pan-human azimuth cellular identities while promoting biologically appropriate integration across tissue sources, we developed a tissue-aware neighborhood benchmarking framework comprising of five complementary metrics: cell-type purity (CT-Purity), which measures the preservation of cell-type identity within local neighborhoods; tissue similarity (CT-TissueSim), which evaluates the agreement between observed and expected tissue-source distributions; tissue mixing (CT-TMix), which combines cell-type purity and tissue similarity; adjusted tissue coverage (CT-TCov), which quantifies the representation of expected tissue sources while accounting for cell-type purity; and raw tissue coverage (CT-TCovRaw), which measures tissue representation independently of cell-type purity. Using these metrics, scRefine maintained high cell-type purity while achieving the highest median tissue similarity (∼0.76), tissue mixing (∼0.71), adjusted tissue coverage (∼0.44), and raw tissue coverage (∼0.50) among the eight evaluated methods (Supplementary Figure S3). In contrast, although most baseline methods preserved relatively high cell-type purity, they exhibited lower tissue-aware mixing and coverage, whereas cellANOVA showed substantially reduced cell-type purity and tissue mixing. These findings suggest that scRefine achieves a favorable balance between preserving distinct cellular identities and integrating cells across biologically relevant tissue sources, without indiscriminately mixing unrelated cell types.

We next assessed whether transcriptionally related cells formed coherent local neighbourhoods by visualising canonical lineage-marker densities across embeddings using sjanpy [52] (Supplementary Figure S4). In scRefine, PTPRC formed a broad but consolidated immune compartment with internal structure, while several alternative methods showed more fragmented PTPRC-positive regions; scVI showed a similarly coherent pattern. ALB, EPCAM, MLANA, and SCGB2A2 likewise formed distinct lineage-associated domains in scRefine. Notably, ALB-positive hepatocytes occupied several separate but highly pure neighbourhoods that showed improved integration between normal-liver and metastatic cells while retaining state-specific differences, including metastatic stress-associated populations (Supplementary Figure S10). These results indicate that scRefine brings transcriptionally related cells together without collapsing biologically meaningful within-lineage heterogeneity, consistent with its auxiliary prediction task preserving informative transcriptional variation during integration.

To complement the qualitative comparisons, we evaluated integration using quantitative metrics designed to capture both batch mixing and biological conservation (Figure 2B). Local dataset mixing was assessed using the integration local inverse Simpson’s index (iLISI), which quantifies the diversity of datasets represented within the neighbourhood of each cell. Higher iLISI values indicate that cells from different datasets are well mixed locally, whereas lower values suggest residual dataset-specific segregation. We additionally calculated the k-nearest-neighbour batch-effect test (kBET) acceptance rate, which evaluates whether the dataset composition of local cellular neighbourhoods is consistent with the global dataset distribution. A high kBET acceptance rate therefore indicates effective removal of batch-associated structure, while a low acceptance rate reflects significant local deviations caused by residual batch effects. Finally, principal-component regression (PCR)-based batch-effect estimates were used to determine the extent to which variation in the integrated representation remained attributable to dataset identity. In this analysis, lower residual batch-associated variance indicates more effective removal of technical variation. Together, these metrics provide complementary assessments of whether integration successfully mixes cells across datasets at both local and global levels, while avoiding reliance on any single definition of batch correction.

Perfect dataset mixing can be achieved trivially by placing all cells into a common undifferentiated representation, but such a solution eliminates cell identity and is therefore biologically uninformative. Conversely, strict separation of annotated populations may preserve apparent cell-type boundaries while retaining substantial technical variation within each population. Integration quality must consequently be evaluated as a balance between these competing objectives.

Across these complementary measurements, scRefine achieved strong dataset-mixing scores while maintaining robust cell-type and graph-structure preservation. Several competing methods performed well for either batch removal or biological conservation but showed weaker performance for the complementary objective. scRefine more consistently balanced these properties, supporting the qualitative observation that it removes study-associated variation without destroying the local structure of biologically distinct populations.

### scRefine improves cell-type annotation accuracy

We compared the annotation performance of scRefine, CellTypist, and scANVI across both broad and fine-grained cell-type hierarchies. At lower resolution, all methods successfully identified the primary cell types; however, scRefine demonstrated superior overall precision and F1 score, with fewer false-positive classifications (Figure 2C). At higher resolution (91 cell types), the performance advantage of scRefine became more pronounced, particularly for closely related immune and progenitor populations that are inherently difficult to distinguish. Under these conditions, the contrastive classifiers based on scRefine maintained greater classification accuracy than those of CellTypist and scANVI, indicating an enhanced capability to discriminate among transcriptionally similar subtypes.

Cell-type-level comparisons further corroborated this observation: the accuracy advantage for transcriptionally similar cell types was most pronounced in the high-resolution setting. Supplementary Figures S5 shows that scRefine, particularly the contrastive k-means implementation, accurately classified a higher proportion of cells across a diverse array of immune and progenitor populations. By contrast, CellTypist and scANVI showed variable performance across fine-grained classes, with accuracy diminishing for several closely related cellular states like DC, DC1, DC2 or Type 1 Helper T-cell and Type 17 Helper T-cell. These findings indicate that scRefine offers a more consistent annotation framework, particularly as label granularity increases and subtype boundaries become less pronounced. However, performance remained contingent on cell type; although the results indicate an overall advantage for scRefine, no single method was universally superior across all populations.

### scRefine preserves tissue-conditioned separation of distinct cellular lineages

We quantified whether integration distorted local transcriptional neighborhoods by measuring contacts between five prespecified, biologically distinct lineage pairs—melanocytic–macrophage, melanocytic–neuroendocrine, melanocytic–epithelial, neuroendocrine–epithelial and hepatocyte–epithelial cells—across 704,188 cells. For each method, we constructed a 30-nearest-neighbor graph in the corresponding latent representation and classified a cell as cross-lineage contact-positive when at least three neighbors belonged to the paired opposite lineage. We evaluated both global contacts and tissue-conditioned contacts, the latter requiring the opposite-lineage neighbor to share the focal cell’s tissue annotation. This design distinguishes broad embedding overlap from local mixing among cells originating from comparable biological contexts.

scRefine produced the lowest prevalence of cross-lineage contact-positive cells across the evaluated methods (Figure 2D; Supplementary Figures S6). Only 1.08% of lineage-eligible cells were contact-positive in the global analysis, compared with 20.31% for Harmony, 43.51% for cellANOVA, 12.58% for scVI, 6.79% for Scanorama, 4.32% for Concord and 1.68% for scCM. This separation was retained after tissue conditioning: 0.58% of lineage-eligible cells were contact-positive with scRefine, versus 2.10%, 4.46%, 1.43%, 0.95%, 1.61% and 1.34% for Harmony, cellANOVA, scVI, Scanorama, Concord and scCM, respectively. Thus, the reduction in inappropriate cross-lineage neighborhood overlap was not explained simply by contacts between cells from different tissue contexts.

At the level of individual lineage comparisons, scRefine retained particularly strong separation between melanocytic and epithelial cells (0.06% tissue-conditioned contact-positive cells), neuroendocrine and epithelial cells (0.04%), and hepatocyte and epithelial cells (0.28%). Melanocytic–macrophage contacts constituted the largest residual cross-lineage signal (0.96%), consistent with the greater potential for shared inflammatory, stress-associated or tumor-microenvironmental transcriptional programs between these populations. Importantly, highlighted cells were sparse and localized in the scRefine embedding, whereas several alternative integrations showed more extensive regions of cross-lineage overlap. These patterns indicate that scRefine limits broad transcriptional blending while retaining the local structure needed to resolve biologically distinct compartments.

Together, these results show that scRefine achieves integration without eroding major lineage boundaries. The consistently low global and tissue-conditioned contact frequencies support preservation of biologically meaningful cellular organization, including within matched tissue contexts where confounding by tissue composition is reduced. Because nearest-neighbor overlap reflects transcriptional proximity rather than physical cell–cell proximity, this analysis should be interpreted as evidence for preservation of lineage-resolved embedding structure rather than spatial co-localization. Nonetheless, the results provide a stringent complementary test that scRefine avoids the overcorrection-associated mixing of distinct cell states observed for several competing methods.

### High-resolution annotation reveals cellular heterogeneity within the metastatic liver microenvironment

We next investigated whether scRefine could resolve metastatic-associated remodeling of the liver microenvironment while preserving subtle hepatocyte-state differences along the portal-to-central axis. Leiden clustering of the scRefine embedding at a resolution of 1.2 identified 66 transcriptionally distinct clusters (0–65), ranging in size from 164 to 37,203 cells (Figure 3A; Supplementary Figure S7). Examination of cluster composition revealed heterogeneous contributions from multiple studies and tissue sources, with several clusters containing cells from diverse datasets, while others exhibited pronounced tissue- or study-specific enrichment. Similarly, the proportions of normal- and tumor-associated cells varied considerably across clusters, with some populations represented in both conditions and others predominantly associated with either normal or tumor tissues. These patterns suggest that scRefine captures shared transcriptional populations across datasets while retaining tissue- and disease-associated cellular heterogeneity, providing a foundation for investigating hepatocyte-state diversity and alterations associated with the metastatic liver microenvironment.

**Figure 3.**
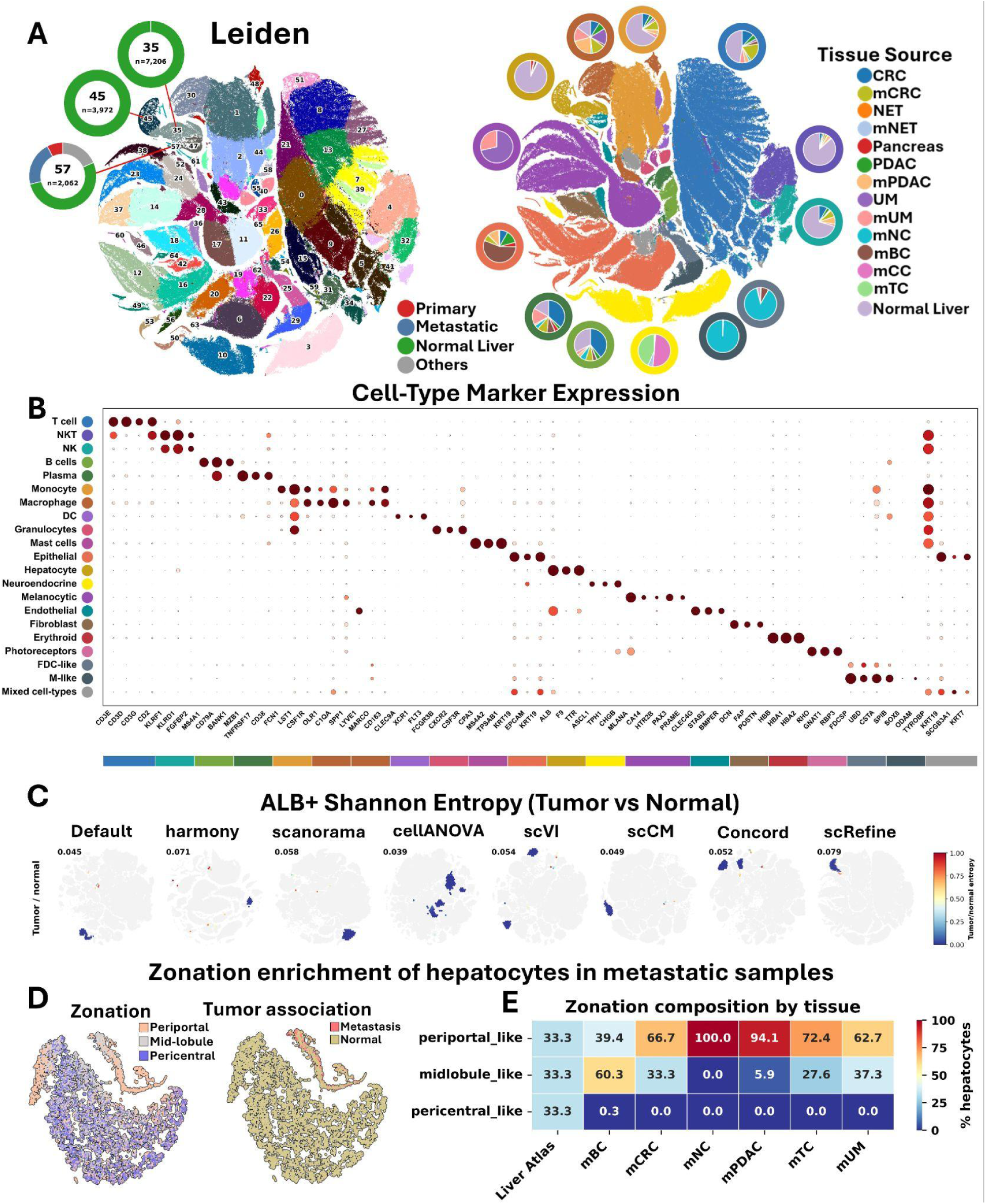
Cellular heterogeneity, lineage-specific marker expression, hepatocyte identity preservation, and metabolic zonation across an integrated primary and metastatic cancer atlas. **(A) Leiden clustering reveals transcriptionally distinct cellular populations and tissue-specific contributions across the integrated cancer atlas**. Two-dimensional visualization of the integrated single-cell transcriptomic atlas illustrating cellular heterogeneity, unsupervised Leiden clustering, and the distribution of individual tissue sources. The left projection displays cells colored according to their Leiden cluster assignments, with numerical labels identifying the individual transcriptionally defined clusters. The atlas comprises cellular populations derived from primary tumors, metastatic lesions, and normal liver tissues, representing diverse malignant and non-malignant cell populations. Selected clusters are accompanied by enlarged donut charts depicting their composition according to four broad sample-origin categories: primary tumors (red), metastatic tumors (blue), normal liver (green), and other sources (gray). Clusters 35 (n = 7,206) and 45 (n = 3,972) are dominated by normal liver-derived cells, whereas cluster 57 (n = 2,062) contains a more heterogeneous distribution of sample-origin categories, including substantial normal liver and metastatic contributions. These differences illustrate the coexistence of tissue-enriched transcriptional populations and populations represented across multiple biological contexts. The right projection displays the same integrated cellular landscape, with cells colored according to their annotated cell-type identities using the same color palette as the cell-type annotations in Panel B. This representation enables visualization of the spatial organization, transcriptional relationships, and distribution of distinct cellular populations across the integrated atlas. The surrounding donut charts provide an additional layer of information by simultaneously illustrating cellular identity and tissue-of-origin composition within selected regions of the embedding. The outer ring represents the annotated cell type, whereas the inner pie chart depicts the proportional contribution of the different tissue sources to the corresponding cellular population. Tissue sources include colorectal cancer (CRC), metastatic colorectal cancer (mCRC), neuroendocrine tumors (NET), metastatic neuroendocrine tumors (mNET), pancreas-derived samples, pancreatic ductal adenocarcinoma (PDAC), metastatic pancreatic ductal adenocarcinoma (mPDAC), uveal melanoma (UM), metastatic uveal melanoma (mUM), additional metastatic cancer categories (mNC, mBC, mCC, and mTC), and normal liver reference tissue. The inner pie-chart segments are colored according to the tissue-source legend, allowing the relative contributions of individual primary, metastatic, and normal tissue categories to be assessed within each annotated cellular population. This visualization reveals that certain cell types, like melanocytic cells, are predominantly associated with specific tissue sources, whereas others are represented across multiple cancer types and tissue contexts, highlighting both tissue-enriched cellular populations and shared cellular identities across the atlas. Together, the cell-type-colored projection and the nested composition charts provide a complementary view of cellular heterogeneity, tissue-of-origin contributions, and the organization of shared and tissue-associated cellular populations within the integrated cancer atlas. **(B)** Canonical lineage-specific marker expression supports the identification of major immune, stromal, epithelial, and specialized cellular populations. Dot-plot visualization of selected marker genes across 21 annotated cell populations, including T cells, natural killer T (NKT) cells, natural killer (NK) cells, B cells, plasma cells, monocytes, macrophages, dendritic cells (DCs), granulocytes, mast cells, epithelial cells, hepatocytes, neuroendocrine cells, melanocytic cells, endothelial cells, fibroblasts, erythroid cells, photoreceptors, follicular dendritic cell-like (FDC-like) cells, M-like cells, and mixed cell-type populations. Dot size represents the proportion of cells expressing each marker within the respective annotated population, while color intensity represents relative expression abundance. Colored symbols adjacent to the cell-type labels and the annotation strip below the plot distinguish the corresponding cellular categories and marker groups. Immune populations exhibit characteristic lineage-associated transcriptional signatures. T cells express the canonical T-cell receptor-associated markers CD3E, CD3D, CD3G, and CD2, whereas NK and NKT populations exhibit expression of cytotoxicity-associated markers, including KLRF1, KLRD1, and FGFBP2. B cells are characterized by expression of MS4A1, CD79A, and BANK1, while plasma cells exhibit markers associated with antibody-secreting differentiation, including MZB1, TNFRSF17, and CD38. Myeloid populations display distinct but partially overlapping transcriptional programs, with monocytes enriched for FCN1, LST1, and CSF1R and macrophages expressing markers such as OLR1, C1QA, SPP1, LYVE1, MARCO, and CD163. Dendritic cell populations exhibit expression of CLEC9A, XCR1, and FLT3, while granulocytes are characterized by FCGR3B, CXCR2, and CSF3R. Mast cells display characteristic expressions of CPA3, MS4A2, and TPSAB1. Non-immune compartments demonstrate similarly distinctive molecular signatures. Epithelial populations express KRT19, EPCAM, and KRT8, whereas hepatocytes exhibit the hepatic functional markers ALB, F9, and TTR. Neuroendocrine populations are distinguished by expression of ASCL1, TPH1, and CHGB. Melanocytic populations show expression of lineage-associated genes including MLANA, CA14, HTR2B, PAX3, and PRAME. Vascular endothelial populations express CLEC4G, STAB2, and BMPER, while fibroblasts exhibit extracellular matrix and stromal remodeling markers such as DCN, FAP, and POSTN. Erythroid populations show prominent expression of hemoglobin-associated genes including HBB, HBA1, and HBA2. A distinct photoreceptor-associated population expresses RHO, GNAT1, and RBP3, consistent with a retinal photoreceptor-like transcriptional identity. Additional FDC-like and M-like populations display distinct marker-expression patterns, while the mixed cell-type category exhibits expression of markers associated with multiple cellular lineages. The overall diagonal enrichment pattern of lineage-associated marker genes supports the transcriptional distinction of the annotated cell populations, while off-diagonal expression illustrates marker sharing, lineage relationships, or potentially heterogeneous cellular assignments requiring further investigation. **(C)** Comparison of tumor-versus-normal transcriptional mixing among albumin-positive cells across eight integration methods. Evaluation of local tumor-versus-normal label heterogeneity within albumin-positive (ALB+) cellular populations using Shannon entropy across eight transcriptomic representations: Default, Harmony, Scanorama, cellANOVA, scVI, scCM, Concord, and scRefine. Each panel displays the corresponding two-dimensional cellular projection, with the complete atlas shown in light gray and the evaluated ALB+ populations highlighted according to their tumor-versus-normal entropy values. The blue-to-red gradient represents entropy values ranging from 0 to 1, with blue indicating low entropy and red indicating high entropy. Lower entropy values represent neighborhoods dominated by a single tumor/normal category, whereas higher values indicate a more heterogeneous distribution of these categories within the evaluated neighborhoods. The numerical summary values displayed above the respective embeddings are 0.045 for the Default representation, 0.071 for Harmony, 0.058 for Scanorama, 0.039 for cellANOVA, 0.054 for scVI, 0.049 for scCM, 0.052 for Concord, and 0.079 for scRefine. Across all approaches, the reported values remain relatively low, indicating limited tumor-versus-normal label heterogeneity under the applied entropy-based evaluation. Nevertheless, substantial method-dependent differences are observed in the spatial arrangement of the highlighted ALB+ populations, suggesting that individual integration strategies differentially organize hepatocyte-associated transcriptional neighborhoods. scRefine achieves the highest displayed entropy summary value (0.079), followed by Harmony (0.071), whereas cellANOVA exhibits the lowest (0.039). These differences indicate variation in the extent of tumor-versus-normal label intermixing within the evaluated populations. Importantly, greater entropy should not be interpreted independently as improved biological preservation, because mixing may reflect successful alignment of transcriptionally equivalent cells or excessive correction of disease-associated differences. This analysis therefore provides a complementary measure of local sample-origin heterogeneity rather than a standalone measure of integration accuracy or preservation of hepatocyte identity. **(D)** Zonation states of hepatocytes in metastatic and normal-liver samples, projected onto the scRefine embedding. Zonation signatures were derived from the human hepatocyte zonation profiles reported by Yakubovsky *et al.* [54], in which Zone 1 corresponds to the central/pericentral region and Zone 8 to the portal/periportal region. Genes exhibiting significant and sufficiently strong monotonic expression changes across Zones 1–8 were selected after excluding stress-, inflammation-, contamination-, and broadly expressed hepatocyte genes. Pericentral and periportal signature scores were calculated separately, and their difference was used as a continuous zonation score. Cut-offs corresponding to the lower and upper tertiles of the normal-liver reference distribution classified hepatocytes as periportal-like, mid-lobule-like, or pericentral-like. The zonation map shows that the hepatocyte compartment is dominated by periportal-like cells, while pericentral-like cells occupy more restricted regions. The adjacent map indicates that metastatic hepatocytes are concentrated in discrete regions of the embedding rather than forming a uniformly distributed population. (E) Relative zonation-state composition of hepatocytes from the normal Liver Atlas reference and individual metastatic tissue sources. Because the classification thresholds were defined using tertiles of the normal-liver reference distribution, reference hepatocytes are divided equally among periportal-like, mid-lobule-like, and pericentral-like states by design. The heatmap on the right summarizes the relative abundance of each zonation-associated hepatocyte population across seven tissue categories: Liver Atlas, mBC, mCRC, mNC, mPDAC, mTC, and mUM. Columns represent the different tissue sources and rows correspond to periportal-like, midlobule-like, and pericentral-like hepatocyte classifications. Values indicate the percentage of hepatocytes assigned to each zonation category within the corresponding tissue source, with colors ranging from blue (low proportion) to red (high proportion). The Liver Atlas reference displays an approximately equal representation of the three zonation categories, with 33.3% periportal-like, 33.3% midlobule-like, and 33.3% pericentral-like hepatocytes. In contrast, metastatic tissue-associated hepatocyte populations exhibit a pronounced shift toward periportal-like and midlobule-like transcriptional identities, with markedly reduced representation of pericentral-like hepatocytes. Among metastatic breast cancer-associated samples (mBC), midlobule-like hepatocytes constitute the largest fraction (60.3%), followed by periportal-like hepatocytes (39.4%), while pericentral-like hepatocytes account for only 0.3%. Metastatic colorectal cancer samples (mCRC) show enrichment of periportal-like hepatocytes (66.7%), with the remaining 33.3% classified as midlobule-like. The strongest periportal-like enrichment occurs in mNC samples, where 100.0% of classified hepatocytes exhibit a periportal-like transcriptional identity. Similarly, mPDAC samples display a predominantly periportal-like composition (94.1%), with 5.9% midlobule-like hepatocytes. In mTC samples, periportal-like hepatocytes comprise 72.4% of the population and midlobule-like hepatocytes account for 27.6%. Metastatic uveal melanoma samples (mUM) demonstrate a comparable pattern, with 62.7% periportal-like and 37.3% midlobule-like hepatocytes. No pericentral-like hepatocytes are represented in the displayed mCRC, mNC, mPDAC, mTC, or mUM compositions. Collectively, these data identify a consistent enrichment of periportal-like transcriptional programs across several metastatic tissue contexts, accompanied by a substantial reduction in the representation of pericentral-like hepatocyte signatures. Given the established metabolic specialization of hepatocytes along the hepatic lobular axis, these differences are compatible with alterations in the representation of hepatocyte metabolic states within tumor-associated liver tissue. Periportal hepatocytes are generally associated with pathways such as gluconeogenesis, ureagenesis, and oxidative metabolism, whereas pericentral hepatocytes are associated with distinct xenobiotic metabolism, glycolytic, and glutamine-synthesis programs. Accordingly, the observed zonation distribution raises the possibility that metastatic tissue environments are associated with changes in the composition or transcriptional state of surrounding hepatocytes. However, transcriptomically inferred zonation does not directly establish anatomical localization, metabolic flux, or tumor-induced cellular reprogramming. Further investigation incorporating hepatocyte-specific differential gene expression, zonation signature scoring, spatial transcriptomics, and metabolic pathway analyses would be required to distinguish genuine hepatocyte reprogramming from differences in tissue sampling, cellular composition, and selective recovery of particular hepatocyte subpopulations.

We achieved high-resolution cell-type annotation by integrating cluster-specific differential gene-expression profiles, the tissue and dataset composition of each Leiden cluster, and evidence from an internally curated cell-annotation database. This combined framework enabled the assignment of cells to both major lineages and more refined cellular states and subtypes (Figure 3A,B, Supplementary Figure 8). Annotation was supported by the spatially restricted expression of canonical marker programs, including CD3D/CD3E and CD4/CD8A for T-cell populations, MS4A1 and CD79A for B cells, S100A8/S100A9 and FCN1 for classical monocytes, FCGR3A and LST1 for non-classical monocytes, and C1QA, C1QB, TREM2, SPP1, MSR1, and OLR1 for distinct macrophage states. Additional marker combinations resolved NK cells, dendritic cells, mast cells, hepatocytes, epithelial populations, endothelial cells, melanocytic tumour cells, and neuroendocrine-like cells (Supplementary Figure S8).

The resulting annotations resolved coherent immune, epithelial, stromal, endothelial, hepatocyte, melanocytic, and neuroendocrine compartments, with lineage-appropriate marker expression supporting most assignments; ambiguous clusters were retained as low-confidence or unassigned (Supplementary Figure S8). Across integration methods, canonical marker populations remained detectable but differed in compactness and continuity, with scRefine maintaining coherent lineage-specific neighbourhoods while preserving transcriptional heterogeneity (Supplementary Figure S9). Lineage-context enrichment, calculated from observed-to-expected contacts among 30 nearest neighbours after accounting for sample composition, further showed that scRefine combined strong lineage self-enrichment with comparatively high within-lineage dataset mixing (Supplementary Figure S10). Concord and scVI retained moderate lineage organisation, whereas cellANOVA approached the compositional null, consistent with possible over-integration in our setting. Notably, scRefine preserved limited mixing of tumour-restricted melanocytic, hepatocyte and neuroendocrine populations, indicating that technical dataset effects were reduced without artificially merging biologically distinct tumour-specific identities. In contrast, broadly shared immune and stromal populations contained contributions from multiple datasets and tissue sources, consistent with successful cross-study alignment of biologically equivalent cells. These findings collectively demonstrate that scRefine supports detailed cell-state annotation and cross-dataset integration while preserving the tissue specificity, lineage identity, and specialised transcriptional programs represented within the LiverMet atlas. Hepatocytes segregate into three transcriptionally distinct clusters, characterized by elevated expression of *CPS1* and *CYP* family genes (cluster 35), *F9* (cluster 45), and *TTR* and *RBP4* (cluster 57) (Figure 3A). Examination of their tissue-of-origin composition (Figure 3A, pie charts), together with Shannon entropy-based analysis (Figure 3C), revealed differences in their association with normal and metastatic tissues. While clusters 35 and 45 were predominantly composed of hepatocytes derived from normal liver samples, cluster 57 contained a substantially higher proportion of hepatocytes originating from metastatic tissues. These findings suggest that hepatocytes within the metastatic liver microenvironment exhibit transcriptional states distinct from those predominantly represented in normal liver, potentially reflecting tumor-associated alterations in hepatocyte identity and function.

To determine whether this hepatocyte organisation was specific to scRefine or similarly retained by other integration approaches, we performed a latent-space neighbourhood analysis of ALB-high, marker-supported liver hepatocytes across all embeddings. Normalised Shannon entropy was used to address two complementary aspects of integration: cell-type entropy measured the extent to which hepatocyte neighbourhoods were contaminated by other lineages, whereas normal-liver/metastasis entropy measured the local integration of hepatocytes from the two disease conditions. Hepatocyte-neighbourhood purity was additionally calculated as the mean fraction of hepatocytes among each cell’s nearest neighbours. These quantities were calculated from latent-space nearest neighbours and should therefore not be interpreted as physical spatial colocalization.

Among the evaluated methods, scRefine produced the lowest mean cell-type entropy (0.002) and the highest mean hepatocyte-neighbourhood purity (0.99), indicating that ALB-high cells were most consistently retained within hepatocyte-dominated neighbourhoods (Supplementary Figure S11A,C). At the same time, scRefine achieved the highest normal-liver/metastasis entropy (0.079), demonstrating improved local integration between hepatocytes derived from normal and metastatic liver samples (Supplementary Figure S11B). Importantly, this increased cross-condition mixing did not obscure disease-associated differences in hepatocyte composition. Clusters 35 (*CPS1* and *CYP* family genes) and 45 (*F9*) remained predominantly composed of hepatocytes derived from normal liver, whereas cluster 57 (*TTR* and *RBP4*) contained a substantially higher proportion of hepatocytes originating from metastatic tissues. Together, these findings suggest that scRefine achieves a favorable balance between hepatocyte lineage preservation and cross-condition integration while retaining distinct hepatocyte transcriptional states associated with normal and metastatic liver microenvironments.

To further investigate hepatocyte heterogeneity, we isolated hepatocytes and re-embedded them for a focused analysis of their zonation transcriptional landscape using zonation markers established in Yakubovsky et al. [54] (Supplementary Figure S12 and 13) [53]. This hepatocyte-specific embedding revealed multiple Leiden subclusters arranged along gradients consistent with hepatic zonation patterns.

The analysis of the lobular zonation distribution of hepatocytes relative to metastatic associations revealed a notable spatial bias: hepatocytes associated with metastatic samples were significantly enriched in the periportal region of the hepatocyte manifold (Figure 3D). Zonation analysis confirmed that these metastasis-associated hepatocytes predominantly localised to zone 1 (periportal) regions, with an estimated 80% to 90% of these cells occupying periportal states, in stark contrast to fewer than 10% observed in pericentral regions (Figure 3D,E; Supplementary Figure S12 and S13). Although these conclusions are based on a relatively limited subset of metastasis-associated hepatocytes, the findings highlight a potentially underappreciated role for SAA1⁺ periportal-like hepatocyte programs in early metastatic niche formation identified SAA1 as a periportal-enriched immune-associated hepatocyte gene in the healthy human liver, while Lee et al. [56] demonstrated that hepatocyte IL-6–STAT3-dependent SAA1/2 production promotes formation of the hepatic pro-metastatic niche [53–56]. Together, these observations provide a plausible mechanistic link between the periportal-like state observed here and metastatic niche priming, although larger cohorts, spatial transcriptomics, and longitudinal sampling will be required to establish its generality and causal role in metastatic progression.

### Recovery of macrophage activation programs and lineage structure across datasets

Robust single-cell integration should effectively delineate major cell lineages while maintaining important neighbourhood structures within closely related immune populations. To assess whether scRefine preserves biologically interpretable lineage organisation, we compared scRefine with CONCORD, scCM, and scVI using melanocyte and macrophage populations from primary and metastatic uveal melanoma samples (Figure 4). The analysis examined whether these methods retained both global lineage separation and the representation of subtle SPP1⁺ tumor-associated macrophage (TAM) states [54,57,58] while distinguishing them from SPP1⁺ malignant melanocytic populations, which have been associated with invasive and metastatic melanoma phenotypes [59–62]. The evaluation considered both major-lineage preservation and the more challenging distinction between SPP1-high melanocytic tumour cells and SPP1-high tumour-associated macrophages.

**Figure 4:**
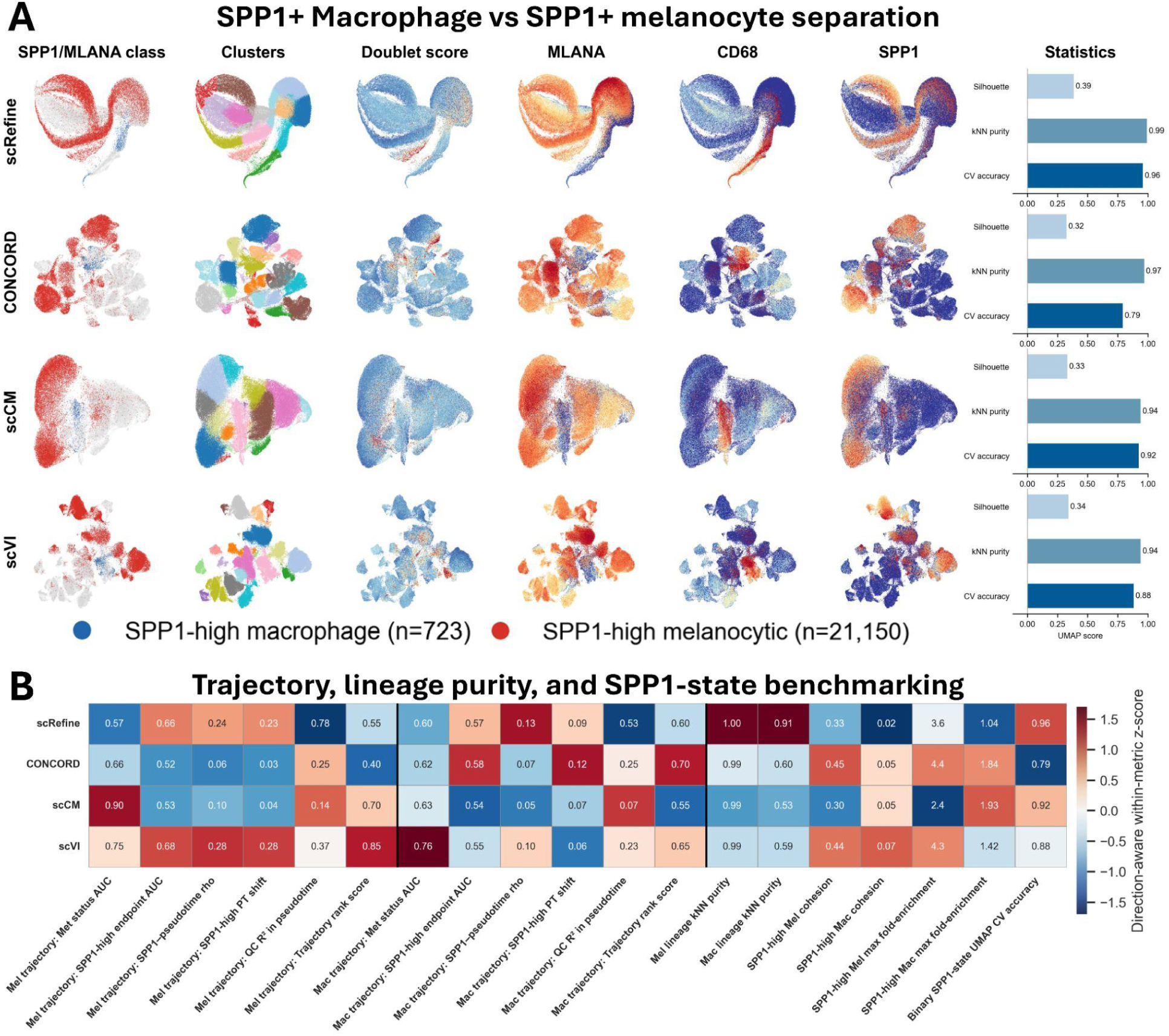
scRefine preserves melanoma–macrophage lineage structure and resolves SPP1-associated cellular states. **(A)** The first column indicates the four cell classes defined according to SPP1, CD68, and MLANA expression. Cells with SPP1 expression ≥ 2.25 were classified as SPP1-high and separated into SPP1+ macrophages (CD68+SPP1+) and SPP1+ melanocytes (MLANA+SPP1+). Among SPP1-low cells (SPP1 < 2.25), cells with MLANA expression ≥ 2.99 were classified as MLANA+ cells, whereas cells not meeting these criteria were assigned to the *Other* category. Thus, the four classes comprised SPP1+ macrophages, SPP1+ melanocytes, MLANA+ cells, and *Other* cells. The final column summarizes four-class UMAP silhouette score, balanced 15-nearest-neighbor purity, and five-fold cross-validated balanced classification accuracy. scRefine achieved the best silhouette score (-0.108) — although it remained slightly negative, and the highest balanced classification accuracy (0.49). scVI and CONCORD produced higher local neighborhood purity (0.056 and 0.55). Across all cells, median doublet scores were 0.120 for SPP1-high macrophages, 0.093 for Other cells, 0.074 for MLANA-high cells, and 0.061 for SPP1-high melanocytes (Kruskal–Wallis *P* < 10⁻³⁰⁰). (**B)** Direction-aware comparison of primary-to-metastatic molecular trajectories, lineage preservation, and SPP1-state organization across embeddings. Melanocyte and macrophage trajectories were constructed independently to avoid implying trans-lineage conversion. Primary, SPP1-low cells were used as trajectory roots, and graph pseudotime was calculated in each latent embedding. Cells were evaluated for metastatic-status ordering, SPP1-high endpoint recovery, correlation between SPP1 expression and pseudotime, SPP1-high pseudotime displacement, QC-associated pseudotime variance, and an aggregate trajectory rank. Additional metrics quantify melanocyte and macrophage kNN purity, adjusted SPP1-high neighborhood cohesion, maximum cluster fold-enrichment, and four-class UMAP accuracy. Values are shown directly; colors represent direction-aware within-metric z-scores, with red indicating better performance. QC R² is direction-reversed so lower technical dependence is favorable. Method-specific UMAP representations generated from scRefine, CONCORD, scCM, and scVI embeddings. Cells are classified as SPP1-high macrophages, SPP1-high melanocytes, MLANA-high cells, or Other using expression-mixture lured by lineage-specific pseudotime are provided in Supplementary Figure S17.

All methods recovered the major biological axis separating melanocytes from macrophages (Figure 4A). Melanocytes generally formed a distinct manifold from macrophages, confirming that the principal lineage distinction was retained across integration approaches. However, the topology and internal organisation of these populations varied substantially. scRefine generated comparatively continuous lineage manifolds with clear internal substructure, whereas CONCORD and scVI divided the data into multiple disconnected islands and scCM produced broader regions of overlap and transition. These differences were reflected by lineage-specific neighbourhood metrics. Melanocyte k-nearest-neighbour purity was consistently high across all embeddings, ranging from approximately 0.98 to 1.00. Macrophage purity was more variable, with scRefine achieving the highest value ( 0.91), compared with CONCORD ( 0.60), scCM ( 0.53), and scVI ( 0.59; Figure 4B).

Marker-expression overlays confirmed the biological identity of the major manifolds. MLANA expression was enriched within the melanocytic compartment, whereas CD68 marked macrophage-enriched regions. SPP1 was detected in subsets of both lineages rather than acting as a macrophage-specific marker. We therefore classified cells as SPP1-high macrophages, SPP1-high melanocytes, MLANA-high cells, or Other using expression-mixture thresholds. This identified 723 SPP1-high macrophages, 21,150 SPP1-high melanocytes, 43,178 MLANA-high cells, and 34,932 Other cells. SPP1-high macrophages displayed the highest median doublet score (0.120), compared with Other cells (0.093), MLANA-high cells (0.074), and SPP1-high melanocytes (0.061; Kruskal–Wallis *P* < 10⁻³⁰⁰). This indicates that SPP1-high macrophages require cautious interpretation because of possible doublet enrichment, whereas the much larger SPP1-high melanocyte population was not associated with elevated doublet scores.

The four molecular classes remained partially overlapping in every UMAP, consistent with their representation as transcriptional states rather than completely discrete cell types. scRefine achieved the highest silhouette score (0.39) and five-fold cross-validated classification accuracy (0.96). CONCORD, scCM, and scVI produced silhouette scores of 0.32, 0.33, and 0.34, respectively, with corresponding cross-validated accuracies of 0.79, 0.92, and 0.88. Neighbourhood purity was highest for scRefine (0.99) followed by concord (0.97) with scCM and scVI both at 0.94. In the complementary UMAP interpretability benchmark, scRefine achieved the highest aggregate score (0.615), compared with scVI (0.538), CONCORD (0.471), and scCM (0.339; Supplementary Figure S16). Its principal advantage was stronger UMAP-level separation between SPP1-high melanoma and macrophage populations, accompanied by greater centroid separation and a substantially lower cross-lineage neighbourhood-mixing risk. The scaled UMAP centroid separation was highest for scRefine (2.7), compared with scCM (2.3), scVI (2.1), and CONCORD (1.6). In addition, scRefine exhibited the highest UMAP-level SPP1-high macrophage kNN cohesion (0.092), although the differences relative to CONCORD (0.090), scVI (0.086), and scCM (0.079) were modest. Notably, this result differed from the latent-space SPP1-high macrophage cohesion metric, for which scRefine showed a lower value (0.02) than CONCORD and scCM (0.05) and scVI (0.07). However, we do not consider the incomplete clustering of SPP1-high melanoma cells as a failed biological resolution. sSingle-cell profiling has shown that melanocytic, invasive, MITF-high, and AXL-high programs coexist within individual melanomas and vary across continuous or partially overlapping cell states [63]. Additional work identified multiple coexisting and reversible melanoma states during treatment adaptation, including melanocytic, invasive, neural crest-like, and drug-tolerant programs [64]. Thus, scRefine’s combination of recoverable molecular-state information and continuous SPP1 organisation may be more biologically appropriate than forcing every expression programme into an isolated cluster.

Primary-to-metastatic graph trajectories were next calculated separately within the melanocyte and macrophage lineages, avoiding an unsupported implication of trans-lineage conversion (Figure 4B). For melanoma cells, scVI achieved the highest direction-aware trajectory rank score (0.85), followed by scCM (0.70), scRefine (0.55), and CONCORD (0.40). scRefine nevertheless recovered an SPP1-associated ordering, with an SPP1-high endpoint AUC of 0.66, an SPP1–pseudotime correlation (Spearman’s ρ) of 0.24, and an SPP1-high median pseudotime displacement of 0.23. However, scVI achieved slightly higher values for all three measures (AUC = 0.68, ρ = 0.28, and displacement = 0.28), indicating stronger SPP1-associated trajectory ordering under these criteria. Furthermore, scRefine exhibited the highest QC-associated pseudotime R² (0.78), compared with CONCORD (0.25), scCM (0.14), and scVI (0.37), suggesting that its inferred trajectory may be more strongly associated with technical variation. Although scCM and scVI more strongly separated primary from metastatic melanoma cells, scRefine retained an interpretable SPP1-associated transcriptional gradient, albeit without outperforming scVI on the trajectory metrics.

For macrophages, scVI showed the strongest metastatic-status ordering (AUC = 0.76), whereas scRefine exhibited the highest SPP1–pseudotime correlation (ρ = 0.13), compared with scVI (0.10), CONCORD (0.07), and scCM (0.05). CONCORD achieved the highest macrophage trajectory rank score (0.70), followed by scVI (0.65), scRefine (0.60), and scCM (0.55). scRefine nevertheless retained the highest macrophage lineage purity (0.91), although its latent-space SPP1-high macrophage cohesion remained lower than that of the alternative methods. The scRefine melanoma trajectory is therefore compatible with published evidence linking SPP1-positive uveal melanoma cells to metastatic risk and invasive behaviour, but the stronger trajectory performance of scVI and the substantial QC-associated pseudotime variation observed with scRefine warrant caution when interpreting the inferred progression. However, these graph-derived pseudotimes represent molecular-state ordering rather than direct temporal or clonal evidence. The lineage-specific UMAPs in Supplementary Figure S17 should consequently be interpreted as visualisations of trajectories calculated in the full latent embeddings, not as proof that individual primary cells directly generated the observed metastatic cells.

Together, these analyses indicate that scRefine provides particularly strong preservation of major-lineage identity and UMAP-level separation of SPP1-high melanocytic and macrophage populations, achieving the highest silhouette score, cross-validated classification accuracy, and composite UMAP interpretability score among the evaluated approaches. Its main advantage is not universal maximisation of every clustering statistic, but the preservation of distinct melanocytic and macrophage transcriptional identities while maintaining a continuous organisation of SPP1-associated cellular states. The complementary UMAP analysis supports strong separation between the cellular sources of SPP1, although this advantage does not extend uniformly to latent-space SPP1-high macrophage cohesion or trajectory reconstruction. In particular, scVI outperformed scRefine in melanoma trajectory ordering and several SPP1-associated pseudotime metrics, while CONCORD achieved the highest macrophage trajectory rank score. Furthermore, the relatively high QC-associated pseudotime R² for scRefine indicates that technical covariation remains an important consideration when interpreting its inferred trajectories. These limitations should be acknowledged, but the combined lineage-purity and UMAP-level separation results support scRefine as a useful representation for distinguishing melanocytic and macrophage populations while retaining interpretable SPP1-associated transcriptional heterogeneity. On this basis, scRefine was selected as the primary representation for subsequent analyses, while the alternative embeddings were retained to assess method-dependent effects.

### scRefine preserves developmental trajectories and rare cellular states across integration

Recovering biologically meaningful insights from large single-cell atlases requires integration methods that preserve both overall lineage structures and detailed developmental trajectories. To assess whether scRefine preserves these features, we benchmarked its embedding against three established methods; namely Concord, Harmony, and scVI, using the developmental brain atlas from Herring et al. as a reference (Figure 5) [56]. The reference embedding provides a baseline layout that captures the main neuronal and non-neuronal populations, enabling a systematic comparison of how each method retains lineage relationships and developmental progress. The analysis showed that scRefine produced a structured and interpretable manifold in which major cell classes, including inhibitory neurons (IN), projection neurons (PN), and non-neuronal populations, remained clearly separated while maintaining overall organisation (Figure 5, Supplementary Figure S16). Compared to the reference layout, which had large, isolated islands with limited spatial continuity between clusters, scRefine generated a more compact representation that preserved the biologically meaningful differentiation of major cell types. Concord produced a similarly compact manifold but exhibited broader mixing of developmental stages within clusters. Harmony, despite substantial batch mixing, fragmented the manifold into many small islands, reducing the overall coherence of the lineage structure. scVI preserved the broad-scale arrangement of neuronal populations but showed wider spatial mixing of developmental stages across different regions within the embedding. The differences in preserving age-related structures were further clarified by visualising cells based on developmental stage (Figure 5, Supplementary Figure S16). In the reference layout, the stage labels appeared patchy and discontinuous across islands, suggesting a lack of explicit organisation along a smooth developmental axis. Concord showed partial improvement in stage continuity but still exhibited considerable mixing. Harmony caused substantial mixing across numerous small islands, complicating the interpretation of developmental progression. scVI maintained some enrichment of stages within wider regions but lacked clear stratification by developmental stage. Conversely, scRefine demonstrated spatially coherent stage organisation, with larger regions enriched in specific stages and smoother transitions between fetal and adult populations, indicating better preservation of biologically relevant temporal structures. To quantitatively evaluate developmental ordering, we visualised developmental age gradients across the embeddings with a continuous age scale derived from harmonised prenatal and postnatal annotations (Figure 5C). The reference embedding concentrated age signals in limited regions, reflecting localised structures rather than a comprehensive global developmental gradient. While Concord improved age continuity, it still showed mixed ages within certain lineages. Harmony displayed scattered age patterns across fragmented islands, indicating weaker preservation of trajectory geometry. scVI maintained broad developmental gradients but often showed diffuse age distributions across large parts of the embedding. In contrast, scRefine produced the most consistent age gradients, especially within lineage-specific trajectories, where cellular arrangements aligned along smooth temporal progressions. This effect was especially noticeable when examining curated developmental trajectories for oligodendrocyte precursor cells (OPC), astrocytes, and layer 5/6 excitatory neurons marked by TLE4 expression. In the scRefine embeddings, the cells along these trajectories formed compact and continuous regions, clearly demonstrating within-lineage progression of developmental age, while background cells remained distinctly separate. Other integration methods often disrupted these trajectories across multiple regions or created mixed-age neighbourhoods that blurred the temporal order.

**Figure 5.**
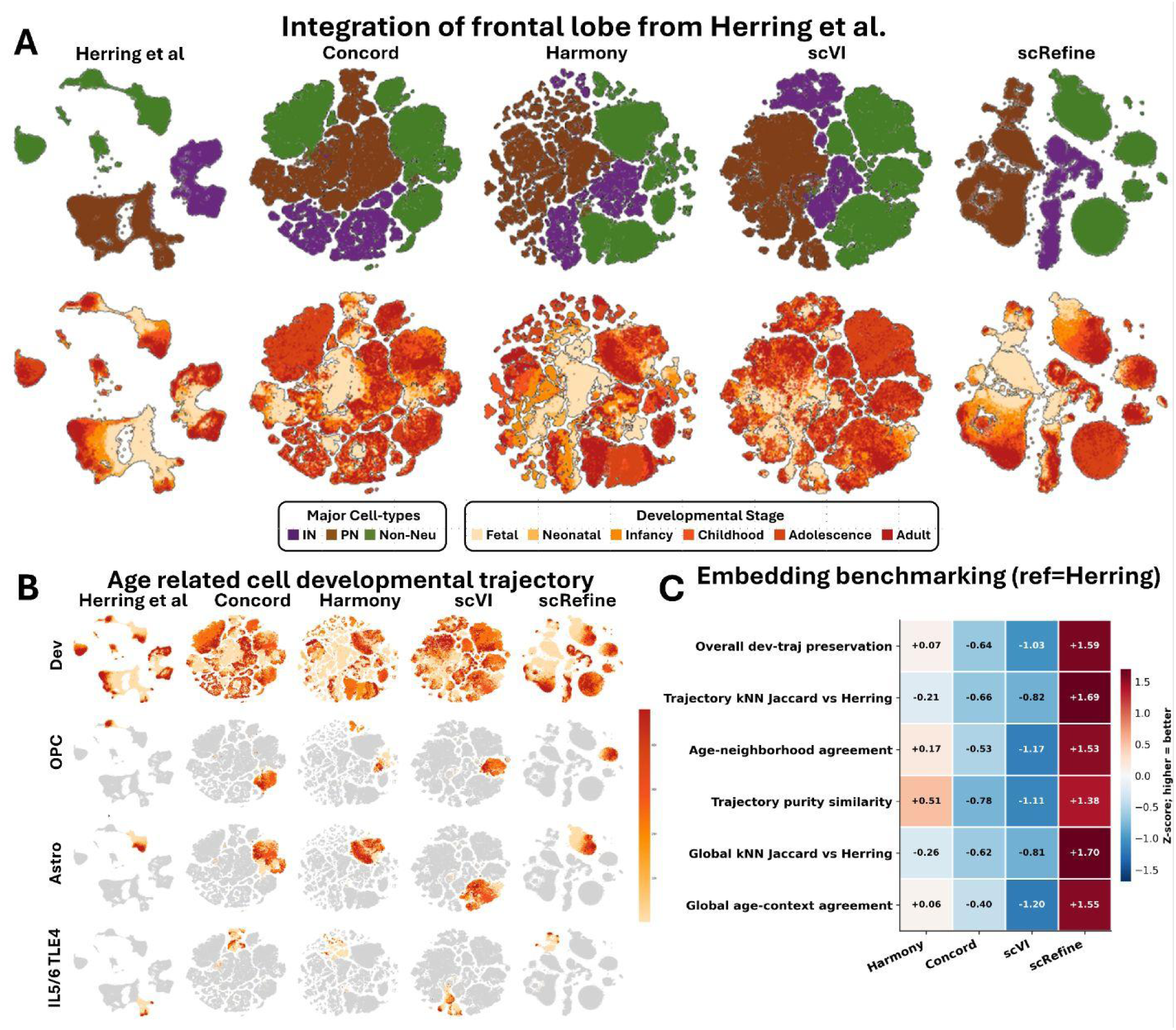
scRefine preserves developmental and lineage structure during integration of brain single-cell atlases. **(A**) Two-dimensional embeddings of the Herring *et al.* [65] reference atlas and four integrated representations generated using Concord, Harmony, scVI, and scRefine. Cells are colored by major cell classes, including inhibitory neurons (**IN**), projection neurons (**PN**), and non-neuronal cells (**Non-Neu**), and by developmental stage from fetal to adult. The reference embedding separates broad lineage classes into distinct regions. Concord and Harmony recover major cell-class structure but show variable compaction and fragmentation of the manifold. scVI preserves broad lineage organization but displays more extensive mixing across developmental stages. In comparison, scRefine maintains clear separation of major neuronal and non-neuronal compartments while preserving an interpretable global organization of the atlas. (**B**) Developmental and trajectory-specific age gradients projected onto each embedding. The top row shows the global developmental age signal across all cells, while lower rows show curated trajectories, including oligodendrocyte precursor cells (**OPC**), astrocytes (**Astro**), and layer 5/6 **TLE4**-positive excitatory neurons. Gray points indicate the full embedding background, and colored points indicate cells from the indicated trajectory, with color intensity representing developmental age in months. scRefine shows compact and coherent trajectory-localized age structure, whereas the other methods show varying degrees of trajectory dispersion, fragmentation, or diffuse age mixing. (**C**) Quantitative benchmarking of integrated embeddings relative to the Herring *et al.* reference layout. The heatmap shows standardized z-scores for developmental trajectory preservation metrics, including trajectory-neighborhood overlap, age-neighborhood agreement, trajectory purity similarity, global nearest-neighbor overlap, and global age-context agreement. Higher z-scores indicate better preservation of biological structure. Across all evaluated metrics, scRefine achieves the strongest performance, supporting improved retention of developmental progression, lineage organization, and local neighborhood relationships compared with Concord, Harmony, and scVI.

To quantify the observed differences, we evaluated each embedding using a comprehensive set of metrics that captured trajectory preservation, structural organisation across developmental stages, and neighbourhood consistency relative to the reference embedding (Figure 5C, Supplementary Figure S17). The metrics included graph monotonicity along developmental pathways, similarity in trajectory direction, preservation of trajectory shape, correlation of stage geometry, similarity in stage density, smoothness of within-lineage age transitions, trustworthiness of neighbourhoods relative to the reference layout, separation of cell types, and overlap among k-nearest neighbours. Across these metrics, scRefine demonstrated the highest overall performance with the strongest aggregate score among the methods tested. In particular, scRefine showed consistently high performance in preserving trajectory geometry, maintaining fidelity to stage structure, and ensuring neighbourhood agreement with the reference embedding (Supplementary Figure S17). In contrast, scVI delivered balanced performance across several global structural metrics, yet exhibited deficiencies in local neighbourhood agreement and lineage-age smoothness. Harmony achieved high scores for lineage age smoothness due to the internal consistency of local neighbourhoods in age; however, this was achieved despite fragmented trajectories across multiple islands, indicating that reliance on local metrics alone can obscure larger global trajectory distortions. Concord showed moderate performance on trajectory-direction metrics but weaker preservation of stage geometry and local neighbourhood structure. Collectively, these findings suggest that scRefine effectively preserves both global lineage organisation and detailed developmental trajectories during integration. By maintaining coherent age gradients and stable neighbourhood structures across lineages, scRefine facilitates accurate reconstruction of developmental progression while preserving rare and transient cellular states that are frequently lost during the integration of large-scale datasets.

### Confidence-aware refinement improves cell-state resolution across tissues and pathological contexts

Reliable interpretation of large integrated single-cell atlases requires not only accurate batch correction but also mechanisms for distinguishing high-confidence cell states from ambiguous or poorly defined cellular entities. We therefore evaluated the ability of scRefine to support confidence-aware topological refinement using the fresh scRNA-seq subset of the metastatic breast cancer atlas reported by Klughammer et al. [66]. The analyzed dataset contained 157,531 cells from 30 biopsies representing six biopsy sources. All cell-type and clinicopathological labels used for evaluation were the author-defined annotations supplied with the published dataset; scRefine, Concord and scVI did not generate or revise these labels. We first compared t-SNE representations generated from scRefine, Concord and scVI (Figure 6A). Cells were colored independently by method-specific Leiden cluster, biopsy source and author-defined cell type. Additional comparisons colored by disease stage and histology are shown in Supplementary Figure S18. scRefine formed discrete neighborhoods corresponding to major author-defined malignant, lymphoid, myeloid, endothelial and stromal populations. Concord and scVI showed substantially greater intermixing of the same author-defined cell types. Biopsy-source labels remained distributed within the major scRefine cell-type neighborhoods, indicating alignment of shared biological populations across samples, while localized biopsy-associated structure was retained. These patterns were consistent with the organization described in the source atlas. Klughammer et al. reported that non-malignant cells grouped primarily according to cell type across biopsies, whereas malignant cells retained stronger patient-specific structure. The source study also identified patient identity, profiling method and biopsy site as major contributors to variation in cellular composition and expression. Accordingly, complete removal of biopsy-associated structure was not treated as the objective of integration. Instead, the desired result was alignment of comparable author-defined cell types while retaining biological variation associated with anatomical source and malignant-cell identity.

**Figure 6.**
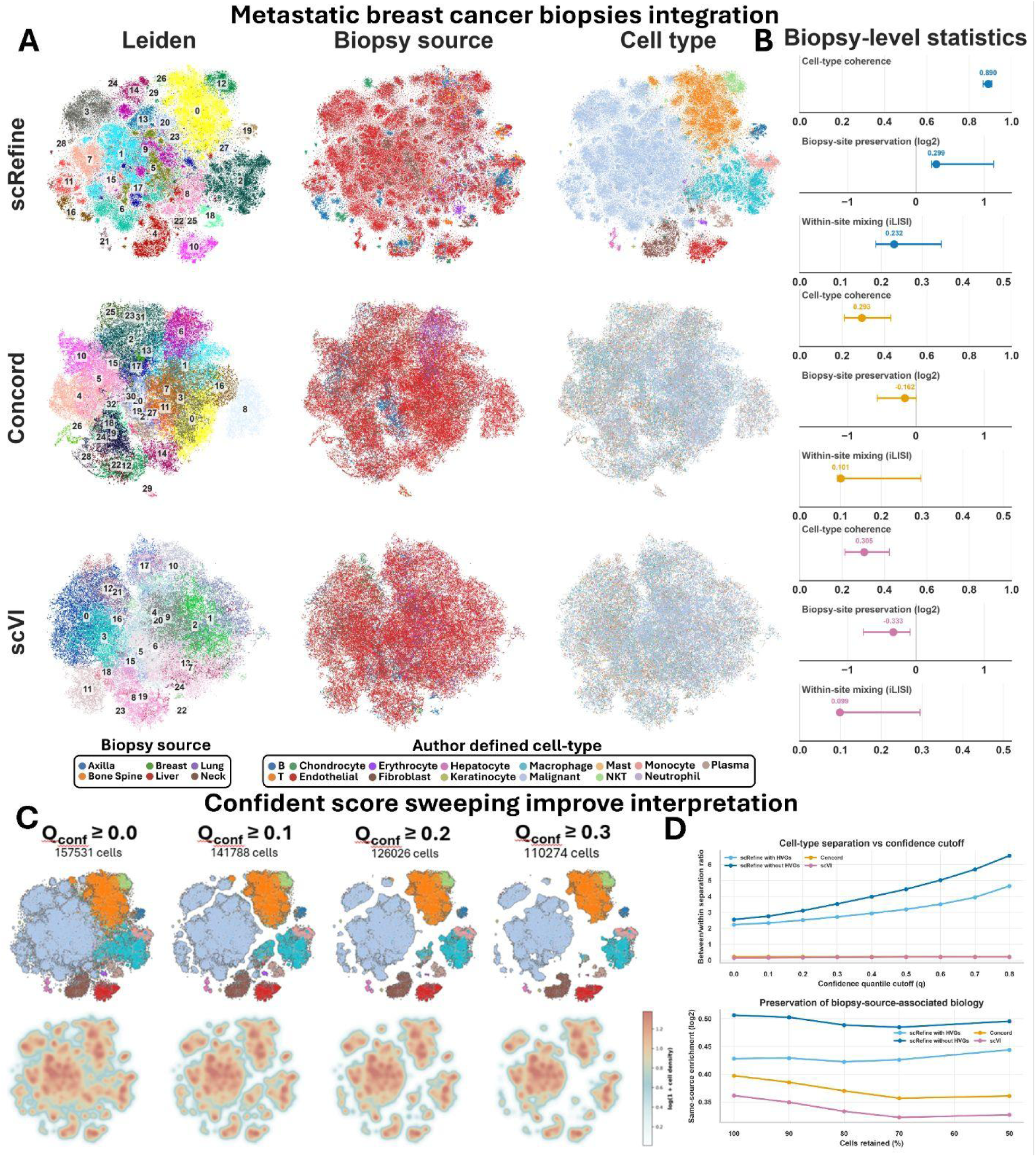
scRefine integration and confidence sweeping preserve cellular identity and biopsy-associated biological structure in fresh metastatic breast cancer scRNA-seq data. The analysis includes 157,531 cells from 30 fresh metastatic breast cancer biopsies. Cell identities and clinical annotations were obtained from the published metastatic breast cancer atlas by Klughammer *et al.* [ 71]. **(A) Comparison of integrated single-cell representations.** t-SNE projections are shown for scRefine, Concord and scVI. Each row represents one integration method, and columns show Leiden clusters, biopsy source and author-defined cell type. Biopsy sources comprise axilla, bone, breast, liver, lung and neck. Author-defined annotations comprise B cells, T cells, chondrocytes, endothelial cells, erythrocytes, fibroblasts, hepatocytes, keratinocytes, macrophages, malignant cells, mast cells, mature NK T cells, monocytes, neutrophils and plasma cells. scRefine produces visibly distinct cellular neighborhoods corresponding to the author-defined annotations, whereas cell types are more extensively intermingled in the Concord and scVI projections. Biopsy-source colors remain distributed within major scRefine cell-type neighborhoods, indicating integration across biopsies, while localized source-associated structure is retained. This representation is conceptually consistent with the original study, which showed that non-malignant cells group primarily according to cell identity across biopsies, whereas malignant-cell profiles retain greater patient and biopsy specificity. The published study also reported that patient identity, profiling method and biopsy site explain substantial variation in cellular composition and expression. Panel a therefore evaluates whether integration preserves this expected biological hierarchy rather than eliminating all biopsy-associated variation. **(B) Biopsy-level integration statistics.** Points indicate the median across biopsies and horizontal bars denote 95% bootstrap confidence intervals for the biopsy-level median. Cell-type coherence is the weighted fraction of graph-neighbor connectivity assigned to cells with the same author-defined cell type. Biopsy-site preservation is the log₂ enrichment of same-source neighbors calculated only among neighbors with the same cell type, thereby separating anatomical structure from differences in cell-type composition. Within-site mixing is the normalized inverse Simpson local diversity index calculated across biopsy identities among same-site, same-cell-type non-malignant neighbors. Malignant cells were excluded from the mixing calculation because the original article demonstrated that their expression profiles are strongly patient specific. scRefine produced a median cell-type coherence of 0.890, compared with 0.293 for Concord and 0.305 for scVI. Median biopsy-source preservation was positive for scRefine (log₂ enrichment, 0.299) but negative for Concord (−0.162) and scVI (−0.333). Within-site non-malignant biopsy mixing was also higher with scRefine (normalized iLISI, 0.232) than with Concord (0.101) or scVI (0.099). These results indicate that scRefine simultaneously improves alignment of shared cell identities and retains measurable biopsy-source-associated biology. **(C) scRefine confidence sweep.** The scRefine cell-type projection is shown after applying increasingly stringent confidence quantile cutoffs. The columns retain 157,531 cells at Q₍conf₎ ≥ 0.0, 141,778 cells at Q₍conf₎ ≥ 0.1, 126,026 cells at Q₍conf₎ ≥ 0.2, and 110,274 cells at Q₍conf₎ ≥ 0.3. The upper row shows retained cells colored by author-defined cell type. The lower row shows matched kernel-density estimates calculated in the original embedding coordinates. Increasing the confidence cutoff preferentially removes cells occupying diffuse regions and interfaces between annotated populations while retaining the dense cores of the principal malignant, immune, myeloid and stromal compartments. **(D) Quantification of the confidence sweep.** The upper plot shows cell-type separation as the ratio of between-cell-type to within-cell-type distances; higher values indicate stronger separation. At each cutoff, identical retained-cell subsets defined by the scRefine confidence score were evaluated for scRefine with highly variable genes, scRefine without highly variable genes, Concord and scVI. Separation increased progressively for both scRefine representations, with the strongest increase observed for scRefine without highly variable genes. Concord and scVI remained comparatively unchanged. The lower plot measures preservation of biopsy-source-associated biology across the same sweep. The y-axis represents within-cell-type same-source enrichment on a log₂ scale. scRefine without highly variable genes consistently retained the strongest biopsy-source signal, followed by scRefine with highly variable genes, Concord and scVI. Moderate confidence filtering therefore increases cell-type separation without eliminating the anatomical structure expected from the original metastatic breast cancer atlas.

The qualitative comparison was supported by three biopsy-level statistics (Figure 6B). Cell-type coherence was defined as the weighted fraction of graph-neighbor connectivity assigned to cells with the same author-defined cell type. Biopsy-source preservation was defined as the log2 enrichment of same-source neighbors after restricting the calculation to neighbors with the same author-defined cell type. Within-site biopsy mixing was quantified as normalized inverse Simpson local diversity across biopsy identities among same-source, same-cell-type non-malignant neighbors. Malignant cells were excluded from the mixing calculation because their patient-specific organization is expected biology in the published atlas. Points in Figure 6B represent biopsy-level medians and horizontal bars show 95% bootstrap confidence intervals for the median.

scRefine achieved a median cell-type coherence of 0.890, compared with 0.293 for Concord and 0.305 for scVI (Figure 6B). Median biopsy-source preservation was positive for scRefine (log2 enrichment, 0.299) but negative for Concord (-0.162) and scVI (-0.333). Within-site non-malignant biopsy mixing was also higher for scRefine (normalized iLISI, 0.232) than for Concord (0.101) or scVI (0.099). Thus, scRefine simultaneously increased agreement with the author-defined cell types, retained measurable anatomical structure after controlling for cell identity and improved cross-biopsy mixing among comparable non-malignant populations.

Broader benchmarking across biological-preservation and batch-integration metrics is presented in Supplementary Figure S21. This analysis included cell-type and macro-cell-type k-nearest-neighbor purity, cell-type silhouette score, between/within-cell-type separation, mutual-exclusion contamination, Leiden agreement with the author-defined cell types, normalized batch iLISI and batch entropy. Metrics were standardized row-wise to facilitate comparison across scales. The scRefine configurations showed the strongest overall preservation of author-defined biological structure, whereas Concord and scVI showed stronger performance on some batch-mixing metrics. The comparison therefore exposed the expected trade-off between maximizing biological separation and maximizing sample mixing, rather than reducing integration quality to a single metric.

We next asked whether scRefine confidence scores could identify cells located in ambiguous regions of the integrated representation. Cells were ranked by the scRefine confidence score and progressively filtered using global quantile cutoffs. The same retained-cell subset was then evaluated in every embedding at each cutoff, preventing differences in cell composition from confounding the method comparison. In Figure 6C, the displayed thresholds retain 157,531 cells at Qconf >= 0.0, 141,778 cells at Qconf >= 0.1, 126,026 cells at Qconf >= 0.2 and 110,274 cells at Qconf >= 0.3. These cutoffs correspond approximately to retaining 100%, 90%, 80% and 70% of cells, respectively.

Visualization of the scRefine representation showed that increasing the confidence cutoff preferentially removed cells from diffuse regions and interfaces between author-defined populations, while preserving the dense cores of the major malignant, immune, myeloid and stromal compartments (Figure 6C). Matched density maps confirmed that the principal high-density regions remained stable as lower-confidence cells were removed. The corresponding matched-cell projections and quantitative purity and contamination analyses across scRefine, Concord and scVI are provided in Supplementary Figure S20.

To quantify visualization-level separation, we calculated the ratio of between-cell-type distance to within-cell-type dispersion using the author-defined cell types (Figure 6D). Within-cell-type dispersion was the mean distance between cells and the centroid of their author-defined cell type, and between-cell-type distance was the size-weighted mean distance between pairs of cell-type centroids. Higher ratios therefore indicate that author-defined cell types are farther apart relative to their internal dispersion. Both scRefine configurations showed progressively increasing separation as the confidence threshold became more stringent, whereas Concord and scVI remained low and changed only marginally. The scRefine configuration without HVGs showed the strongest separation across the confidence sweep (Figure 6D). This result should be considered together with Supplementary Figure S19, where the HVG-based configuration attained the strongest composite biological-preservation score across a broader collection of metrics. The two analyses therefore indicate metric-dependent strengths rather than a universal advantage of one feature-selection strategy.

We additionally tested whether the gain in cell-type separation was obtained by erasing anatomical structure. Across all retained-cell fractions, scRefine without HVGs showed the strongest within-cell-type biopsy-source enrichment, followed by scRefine with HVGs, Concord and scVI (Figure 6D). Biopsy-source enrichment changed only modestly over the confidence sweep, whereas scRefine cell-type separation increased substantially. Confidence filtering therefore enriched the most coherent regions of the scRefine representation without eliminating the biopsy-source signal expected from the original atlas.

## Discussion

The integration of single-cell datasets from various tissues, studies, and biological contexts remains a significant challenge in single-cell transcriptomics. Large reference atlases are often assembled from diverse datasets that differ in sequencing platforms, experimental protocols, and biological compositions. These differences create notable batch effects that can mask biological structures, distort developmental trajectories, and complicate cell-type annotations. Here, we introduce scRefine, a unified framework that brings together several key steps in single-cell data analysis—including batch correction, data integration, cell-type annotation, and confidence assessment—within a single model. Rather than handling these tasks independently, scRefine learns them jointly, generating integrated representations that retain meaningful biological relationships while delivering accurate cell-type predictions with calibrated confidence estimates. Furthermore, the design principles underlying scRefine favour the retention of biologically meaningful transcriptional topology while limiting the overcorrection of genuine biological variation Our results demonstrate that scRefine achieves robust integration across heterogeneous datasets while maintaining clear biological organisation of cell populations. In multi-tissue integration benchmarks, scRefine produces embeddings that not only minimise dataset-driven effects but also reflect meaningful biological axes, such as tumour versus normal states and lineage-specific marker expressions.

Compared to existing methods including Harmony, scVI, Scanorama, and Concord; scRefine yields more balanced embeddings that avoid both under-correction, indicated by dataset-driven clustering, and over-correction, marked by artificial homogenization of biologically distinct populations. These findings highlight the importance of explicitly modelling hierarchical sources of variation, enabling scRefine to effectively disentangle global dataset effects from shared biological signals. Beyond improving overall integration, scRefine also enhances the resolution of cell-type annotations and lineage structures. In integrated multi-tissue atlases, scRefine embeddings show clear separation of major immune, stromal, epithelial, and parenchymal cell populations, while also maintaining consistent neighbourhood relationships within each lineage. The expression patterns of marker genes align closely with expected biological compartments, increasing the interpretability of the learned embeddings. Notably, scRefine enables high-resolution identification of hepatocyte subpopulations and reveals a periportal hepatocyte programme associated with metastatic disease. These hepatocytes display a characteristic acute-phase transcriptional signature including HP, FGA/FGB/FGG, and SERPINA1, along with suppression of typical pericentral markers such as GLUL and CYP2E1. This pattern matches the activation of injury-responsive hybrid hepatocyte progenitors previously observed in regenerative liver biology, suggesting that early metastatic niche formation may involve periportal hepatocyte responses before subsequent pericentral remodelling. Our findings further suggest a biphasic porto-central model of metastatic niche formation in the liver. In this model, periportal hepatocytes initiate a localised acute-phase and extracellular matrix response that may help trap and stabilise tumour cells near portal inflow regions. As metastases grow and lobular architecture remodels, growth may shift towards pericentral areas rich in regenerative metabolic programs. Although this hypothesis requires further validation through spatial and longitudinal datasets, our results demonstrate how advanced integration and annotation frameworks can reveal subtle tissue-state programs that are otherwise hidden by technical variation. Additionally, scRefine improves the recovery of functional subpopulations within immune compartments. Benchmarks across macrophage and melanocyte populations show that, although primary lineage separation is consistent across methods, the internal structure of macrophage states varies significantly depending on the embedding. scRefine achieves the highest neighbourhood purity for SPP1⁺ macrophages, an activated state associated with inflammatory and tumour microenvironment programs. Importantly, scRefine embeddings cluster SPP1-expressing cells into coherent submanifolds rather than dispersing them across disconnected clusters, thus enhancing interpretability for downstream analyses such as state-specific differential expression or trajectory inference.

scRefine demonstrates a clear advantage in its ability to preserve developmental pathways and rare cell states during integration. Developmental benchmarking with brain atlas data shows that scRefine effectively maintains consistent lineage structures while ensuring smooth developmental gradients across both fetal and postnatal stages. Quantitative metrics confirm that scRefine outperforms other embeddings in preserving trajectory geometry, stage organisation, and neighbourhood consistency relative to the reference atlas. These findings suggest that the hierarchical latent structure used by scRefine is vital for capturing multi-scale biological variation, enabling the simultaneous representation of global lineage connections and detailed developmental dynamics. Furthermore, scRefine allows confidence-aware refinement of integrated atlases. By assigning confidence scores to cell embeddings, it facilitates the gradual exclusion of ambiguous cells near cluster boundaries or exhibiting poorly defined transitional states. Removing low-confidence cells results in more compact and distinct cell-type manifolds, with noticeable improvements in separation metrics as confidence thresholds rise. This provides a practical approach to constructing high-confidence reference atlases while preserving the broader biological landscape of the entire dataset. However, several limitations remain. Firstly, certain biological insights—for example, the periportal enrichment of metastasis-associated hepatocytes—are currently based on relatively small cell subsets and need validation through larger cohorts and independent spatial transcriptomic datasets. Secondly, although scRefine enhances neighbourhood coherence and trajectory preservation, integrating highly heterogeneous datasets with limited overlap in cell states remains an open challenge. Future iterations of scRefine could benefit from incorporating explicit spatial context, multimodal data, or graph-based priors to further improve integration across diverse datasets.

Overall, these findings highlight the importance of combining generative modelling, contrastive learning, and biological priors in building comprehensive single-cell reference atlases. Hierarchical latent structures aid scRefine in distinguishing dataset-specific technical variation from shared biological signals, while contrastive objectives sharpen the depiction of subtle differences among closely related cell states. Conditioning the embedding on biologically meaningful markers further ensures that the learned representation remains interpretable for downstream biological analysis. In conclusion, scRefine provides a robust framework for integrating heterogeneous single-cell datasets while maintaining biological structure, developmental trajectories, and functional cell states. Its capability for biologically accurate integration and high-resolution annotation across tissues and datasets supports the development of large-scale reference atlases and paves the way for discovering novel cellular programs in health and disease.

## Methods

### Overview to the scRefine framework

To address batch effects, scRefine employs dataset-specific latent priors that influence the generative process. Each dataset is associated with a dedicated latent factor, allowing the model to capture biases related to the dataset, such as differences in library size, sequencing depth, and technical noise. These dataset-specific factors are multiplicatively combined with a global Gaussian prior. This method enables the model to adapt reconstruction strategies for each dataset while preserving a shared embedding space. Thus, the model first infers a primary latent variable *z*_*i*^(0) ∈ ℝ^*d*, regularised using a global Gaussian prior, that represents biological variation shared across datasets.

For every batch factor *m* ∈ {1, …, *M*}, the model defines a corresponding latent variable *z*_*i*^(*m*) ∈ ℝ^*d*. Its prior is a Gaussian distribution conditioned on the primary latent variable *z*_*i*^(0) and on the observed category *b*_*im* of batch factor *m* and represents the technical variation

Practically, dataset identity is represented through learned embeddings provided as input to both the encoder and decoder, ensuring that the model remains aware of the data’s origin during both inference and generation. Importantly, (only the primary latent variable where lower-level variables capture fine-grained cellular variation and higher-level variables encode broader dataset-specific or technical effects), is used as the final embedding for downstream analysis, while higher-level latent variables are designed to capture and mitigate dataset-specific variations. This distinction allows scRefine to achieve effective integration without losing biologically relevant differences across datasets. To condition the latent embedding according to previously established expert knowledge, marker genes can be used for either classification or contrastive learning. Details of the implementation can be found in the following sections.

Only the primary latent representation is used as the final embedding for downstream analyses. The additional latent variables are used within the generative pathway to model batch-specific and technical variation.

### Hierarchical variational autoencoder

Conventional variational autoencoders model gene expression using a single latent variable, often conflating biological variation with batch-specific effects. To address this limitation, scRefine introduces a hierarchical latent structure comprising multiple latent variables, each capturing distinct levels of variation. Unlike standard hierarchical variational autoencoders, which impose a rigid sequential dependency among latent variables, scRefine employs a partially decoupled hierarchy.

For each cell i, the model defines a primary latent variable *z*_*i*^(0) ∈ ℝ^*d*, together with M additional latent variables *z*_*i*^(1), …, *z*_*i*^(*M*). The primary latent variable *z*_*i*^(0) is intended to encode biological variation shared across datasets and is used as the final representation for downstream analyses

In a conventional hierarchical variational autoencoder, the latent variables are often arranged as a strictly sequential hierarchy: *z*_*i*^(0) → *z*_*i*^(1) → *z*_*i*^(2) → … → *z*_*i*^(*M*).

Here, every latent variable depends on the preceding level. Consequently, information associated with one factor may be passed through several intermediate variables, and dependencies between technical factors may be introduced even when those factors are conceptually independent. scRefine instead employs a partially decoupled hierarchical structure. Each additional latent variable depends directly on the primary latent variable rather than on the preceding batch-specific latent variable:

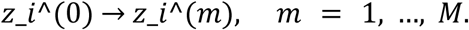

The resulting dependency structure can be represented as

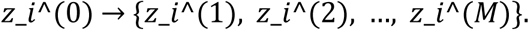

All secondary latent variables share the same biological baseline *z*_*i*^(0), but they do not directly depend on one another. This structure allows each batch-specific factor to model a distinct source of technical variation without requiring information to pass through latent variables associated with other factors

The conditional independence assumption is expressed as

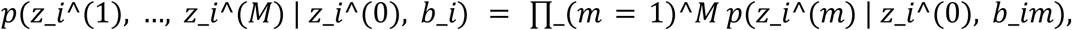

where *b*_*im* denotes the observed category of cell i for batch factor m. Each secondary latent variable therefore has its own conditional Gaussian prior. This design allows multiple latent variables to depend directly on the primary latent representation. This modification enables the independent modelling of dataset-specific factors, thereby allowing clear disentanglement of technical and biological signals. The encoder maps observed gene expression profiles into latent distributions characterised by mean and variance vectors, while the decoder reconstructs gene expression from sampled latent variables. The reconstruction process uses a zero-inflated negative binomial (ZINB) distribution, effectively capturing both overdispersion and dropout, which are common in single-cell RNA sequencing data.

### Marker-conditioned representation learning

To ensure that the learned embedding accurately reflects biologically relevant structures, scRefine incorporates prior knowledge by conditioning on marker genes. A carefully curated set of marker genes, associated with established cell identities or differentiation programs, guides representation learning. Let *M* = {*g*₁, …, *g*_*G*} be a predefined set of *G* marker genes associated with known cell identities, cellular states, or differentiation programs. Because individual expression values are continuous and may differ substantially in scale between genes, the expression of each marker gene is discretised into K ordered bins

For each marker gene g, a discretisation function *B*_*g* maps the observed expression value to an ordinal target *t*_*ig* = *B*_*g*(*x*_*ig*), where *t*_*ig* ∈ {0, 1, …, *K*}. The bin boundaries are defined separately for each marker gene. The primary latent representation *z*_*i*^(0) ∈ ℝ^*d* is then supplied to a marker-prediction network implemented as a multi-layer perceptron. Although the target values represent discrete bins, the network output is treated as a continuous estimate of the ordinal bin position. The marker-prediction loss is then calculated using the mean squared error. Therefore predicting a neighbouring bin is penalised less strongly than predicting a distant bin.

This auxiliary prediction task introduces a supervised signal that helps the embedding retain information pertinent to key biological markers while remaining robust to technical noise. Alternatively, scRefine provides a contrastive module, employing NT-Xent loss to condition the latent embedding according to the cosine similarity of the chosen marker gene set, explained in more detail below (Contrastive representation learning). By directly integrating marker-based supervision into the latent space, scRefine balances unsupervised representation learning with biologically informed constraints, thereby enhancing both interpretability and annotation accuracy.

### Contrastive representation learning

As a complementary methodology, scRefine incorporates a contrastive learning module for representation learning.

Using the input representation of cell i in a minibatch of size B *x*_*i* ∈ ℝ^*d*, *i* = 1, …, *B*, the model uses a projection to calculate a latent space *z*_*i* = *g*_**φ**(ℎ_*i*), before using cosine distance *sim*(*i*, *j*) = (*z*_*i*^*T z*_*j*) / (‖*z*_*i*‖₂ ‖*z*_*j*‖₂) to evaluate similarity.

To determine how related cells in the embedding are we make use of marker-gene expression on a continuous scale.

Let *e*_*ig* = *log*(1 + *x*_*ig*) be the log-normalised expression of marker gene *g* ∈ *G* in cell i. For each gene, a pair of cells (*i*, *j*) gets a soft positive weight from a Laplacian kernel over their expression difference:

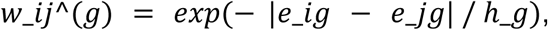

where the bandwidth ℎ_*g* is the standard deviation of gene g within the minibatch, so the kernel adapts to each gene’s scale. Cells with similar expression of gene g are therefore strong positive pairs. Cells with very different expressions get weights near zero and act mostly as negatives.

For any cell i the the log-probability of the pair (i, j) is then:

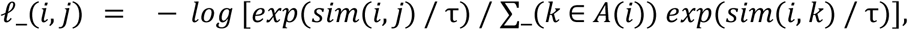

where τ > 0 denotes the temperature parameter. For all positive weights we then calculate for a gene g: :

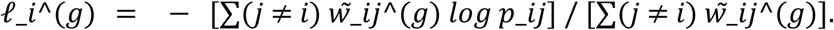

The complete marker gene based contrastive loss is then defined as:

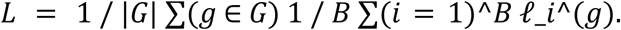

This objective fosters the development of a structured latent space, in which biologically related cells are grouped while maintaining clear distinctions between different cell types.Both HVAE-based and contrastive embeddings are interchangeable for downstream analyses, offering flexibility contingent on data availability and the quality of annotations.

### Loss function and optimization

scRefine is trained using a combined objective consisting of a zero-inflated negative binomial reconstruction loss, the hierarchical variational regularisation terms, and the auxiliary marker-gene prediction loss:

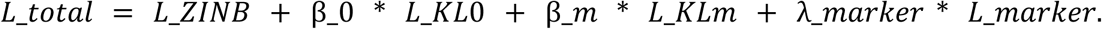

We employ the Zero-Inflated Negative Binomial ZINB framework to address overdispersion due to technical and biological zeros in the data. The ZINB framework achieves this by combining a Negative Binomial distribution, which accounts for overdispersion, with an additional zero-inflation component that explicitly models dropout events. Standard MSE-Loss failed to provide adequate results, due to the higher variability of the scRNA-seq data and an excess of zeros.

The Zero-Inflated Negative Binomial (ZINB) reconstruction loss models the observed gene-expression counts *x*_*ig* using a mixture of a negative binomial distribution and an additional probability mass at zero. For each cell i and gene g, the decoder predicts the mean expression µ_*ig*, dispersion θ_*ig*, and zero-inflation probability π_*ig*:

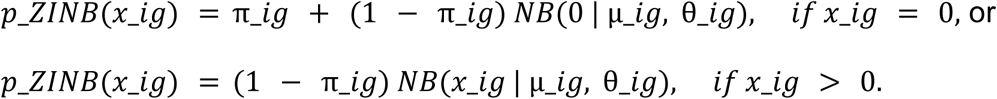

The reconstruction loss is the negative log-likelihood:

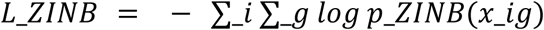

The hierarchical VAE additionally regularises each approximate latent posterior toward its corresponding prior. The primary latent variable is regularised toward the global Gaussian prior:

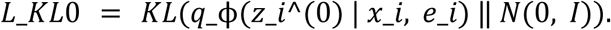

Each additional latent variable is regularised toward its conditional batch-specific prior:

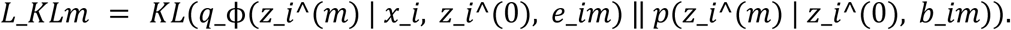

Finally, the auxiliary marker-prediction task estimates the ten-bin marker-expression targets t_ig from the primary latent representation. Its loss is

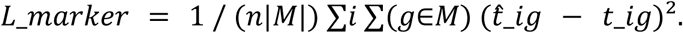

The weighting parameters β_0, β_*m*, and λ_*marker* control the relative contributions of latent regularisation and marker-based supervision

### Library size modeling and batch correction

In addition to latent factorisation, scRefine explicitly models the effects of library size using hierarchical latent variables. Dataset-specific latent factors are used to predict scaling components that adjust the reconstructed gene expression, thereby enhancing robustness to variations in sequencing depth and capture efficiency.

This methodology circumvents the need for explicit batch-correction procedures by integrating technical adjustments directly into the generative model. As a result, it facilitates more accurate preservation of biological variation.

### Post hoc neighbourhood-based confidence estimation and filtering

To enhance the reliability of annotations, scRefine incorporates an additional optional neighbourhood-based refinement step. Using the learned latent space *x*_*i* ∈ ℝ^*d*, *i* = 1, …, *n* and a previous label assignment based on clustering *y*_*i* ∈ *C* = {1, …, *K*}, a radius-based nearest-neighbour classifier estimates label assignment confidence based on neighbourhoods by *N*_*r*(*i*) = {*j* : ‖*z*_*i* − *z*_*j*‖₂ ≤ *r*} using a standardized dimensionality reduced embedding *z*_*i*. For each cluster *c* ∈ *C*, the local cluster probability is calculated as the proportion of cells within the neighbourhood of cell i that belong to cluster c: *p*_*i*(*c*) = [∑_(*j* ∈ *N*_*r*(*i*)) *I*(*y*_*j* = *c*)] / |*N*_*r*(*i*)|, with *I*(·) denoting the indicator function. The The resulting probability vector *p*_*i* = (*p*_*i*(1), …, *p*_*i*(*K*))describes the local distribution of predefined cluster labels around cell i. The confidence of the existing assignment is defined as the probability associated with the cell’s assigned cluster:

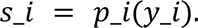

Cells whose confidence falls below the threshold of their assigned cluster are marked as ambiguous and may be excluded from subsequent analyses. This procedure identifies cells whose cluster assignments receive weak support from their local neighbourhood and thereby improves the robustness of downstream biological interpretations.

### Benchmarking metrics

To benchmark scRefine against existing data-integration and dimensionality-reduction methods, we combined established metrics for batch correction and biological conservation with two cell-type-aware tissue-integration metrics introduced here. This framework allows us to compare biological label conservation and batch-correction. Local integration was quantified in the high-dimensional representation generated by each integration method, whereas the corresponding two-dimensional t-SNE embedding was used only for visualization of the resulting per-cell scores.

#### 1. Cell-type-specific tissue reference

To distinguish technical tissue segregation from biologically expected tissue separation, the set of tissues represented by each major cell type was defined before evaluating local tissue mixing. Let *C*_*c* denote the set of cells assigned to cell type c, and *T*_*t* the set of cells originating from tissue t. The cells belonging to cell type c and tissue t are therefore given by the intersection: *C*_*c* ∩ *T*_*t*.

Tissue t was considered represented for cell type (c) when at least 30 cells belonged to Both *C*_*c* and *T*_*t*, and these cells constituted at least 0.5% of all cells of type c. The expected tissue set for cell type c was therefore defined as

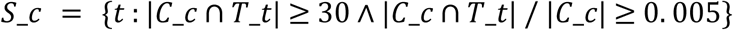

This reference was fixed across integration methods so that every method was evaluated against the same biological cell-type-by-tissue composition. For each tissue (*t* ∈ *S*_*c*), the reference tissue probability distribution *Q*_*c* was defined from the relative number of cells in the corresponding intersection:

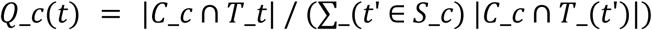

#### 2. Cell-type-directed tissue mixing (CT-TMix)

Cell-type-directed tissue mixing (CT-TMix) was designed to reward cross-tissue integration only when local biological cell-type identity was simultaneously preserved. For each focal cell, the local neighborhood was therefore defined among all cells in the integrated representation rather than restricting the search to cells of the same type. This prevents an integration method from receiving a high tissue-mixing score simply by indiscriminately mixing distinct cell types.

For a focal cell i of cell type c, the distance-weighted local cell-type purity *P*_*CT*(*i*) was defined as the total neighbor probability mass assigned to cells of the same cell type that originated from one of the expected tissues in *S*_*c*:

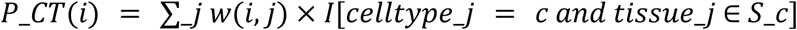

Here, *w*(*i*, *j*) denotes the normalized distance-derived weight of neighbor j, and I[.] is the indicator function. *P*_*CT*(*i*) therefore approaches 1 when the local neighborhood is dominated by cells of the same biological cell type and decreases when other cell types occupy the neighborhood.

Tissue mixing was then evaluated within the same cell-type fraction of the neighborhood. The weights of neighbors with cell type c and tissue in S_c were renormalized to sum to 1 and used to construct the observed local tissue distribution P_i across the expected tissues. This distribution was compared with the reference tissue distribution Q_c using the Jensen-Shannon divergence [67]:

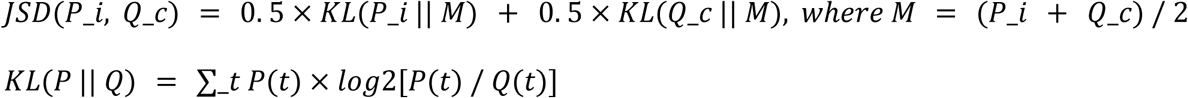

Using base-2 logarithms, JSD ranges from 0 for identical distributions to 1 for maximally divergent distributions. Tissue-composition similarity was therefore defined as 1 – *JSD*(*P_i, Q_c*).

The final CT-TMix score combined local cell-type preservation with appropriate cross-tissue mixing:

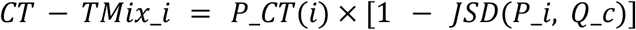

CT-TMix ranges from 0 to 1. High values therefore require both preservation of the focal cell type and a local tissue composition that resembles the expected tissue composition of that cell type. Indiscriminate mixing of different cell types is penalized through *P*_*CT*(*i*), while cell types that are biologically restricted to particular tissues are not penalized for the absence of irrelevant tissues. For cell types represented in only one eligible tissue, the tissue-composition similarity term was set to 1, such that CT-TMix reduces to local cell-type purity

#### 3. Cell-type-directed tissue coverage (CT-TCov)

Cell-type-directed tissue coverage (CT-TCov) was calculated as a complementary measure of whether the expected tissue sources of a cell type were represented within its local neighborhood. For each focal cell i of cell type c, the raw coverage C_raw(i) was defined as the fraction of tissues in S_c that were represented by at least one same-cell-type neighbor within the 90-neighbor local neighborhood:

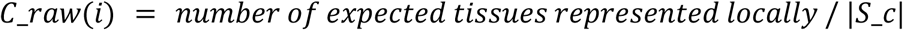

The raw coverage score was then adjusted by the local cell-type purity *P*_*CT*(*i*):

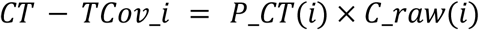

CT-TCov therefore approaches 1 only when the local neighborhood both preserves the focal cell type and contains cells from all tissues expected for that cell type. In contrast to CT-TMix, CT-TCov considers only whether an expected tissue is present locally, irrespective of its relative abundance. CT-TCov therefore captures tissue-source coverage, whereas CT-TMix additionally evaluates whether the local tissue composition is consistent with the expected tissue distribution.

#### 4. Additional quality measurements

Batch correction metrics adapted from existing packages [68–70]

- Graph connectivity: Uses knn graph to evaluate whether cells with the same label form a connected component (range 0–1; higher values indicate better connectivity).
- Integration local inverse Simpson’s index (iLISI): Follows the general LISI framework [27] to estimate the effective number of batches within the local neighbourhood (range 0–1; higher values indicate better batch mixing).
- kNN batch-effect test (KBET): Tests whether the batch composition within local cell neighborhoods matches the expected batch composition. Rejection rates are averaged across biological labels and subtracted from 1, with higher values indicating better batch mixing.
- Principal component regression comparison (PCR comparison): regresses principal components on batch labels to quantify variance contribution of batch effects before and after integration (rescaled to 0–1).

Evaluation of cluster quality:

- silhouette score: used to calculate if clusters are well separated and compact (rescaled to 0–1; higher values indicate better conservation).
- NMI (normalized mutual information): Measures shared information between leiden clusters and underlying true labels, ranging from 0 (no correspondence) to 1 (perfect correspondence).
- ARI (adjusted Rand index): Agreement between leiden clusters and underlying true labels, ranging from 0 (random) to 1 (perfect match).

## Data availability

All datasets used in this study are publicly available from the original publications and repositories cited in the manuscript. Accession numbers and dataset sources are listed in Supplementary Table S1.

## Code availability

The scRefine framework is available as open-source software at https://github.com/BackofenLab/scRefine under the MIT License. The repository contains the full source code, documentation and installation instructions for reproducing the analyses presented in this study. scRefine is built with PyTorch and supports scalable training on large-scale single-cell datasets. The model is designed to work with normalised gene expression matrices and can incorporate multiple datasets simultaneously via dataset embeddings. Hyperparameters governing latent dimensionality, loss weighting, and marker supervision can be adjusted according to the dataset’s complexity and the specific objectives of the analysis.

## Supplementary File

**Figure S1:**
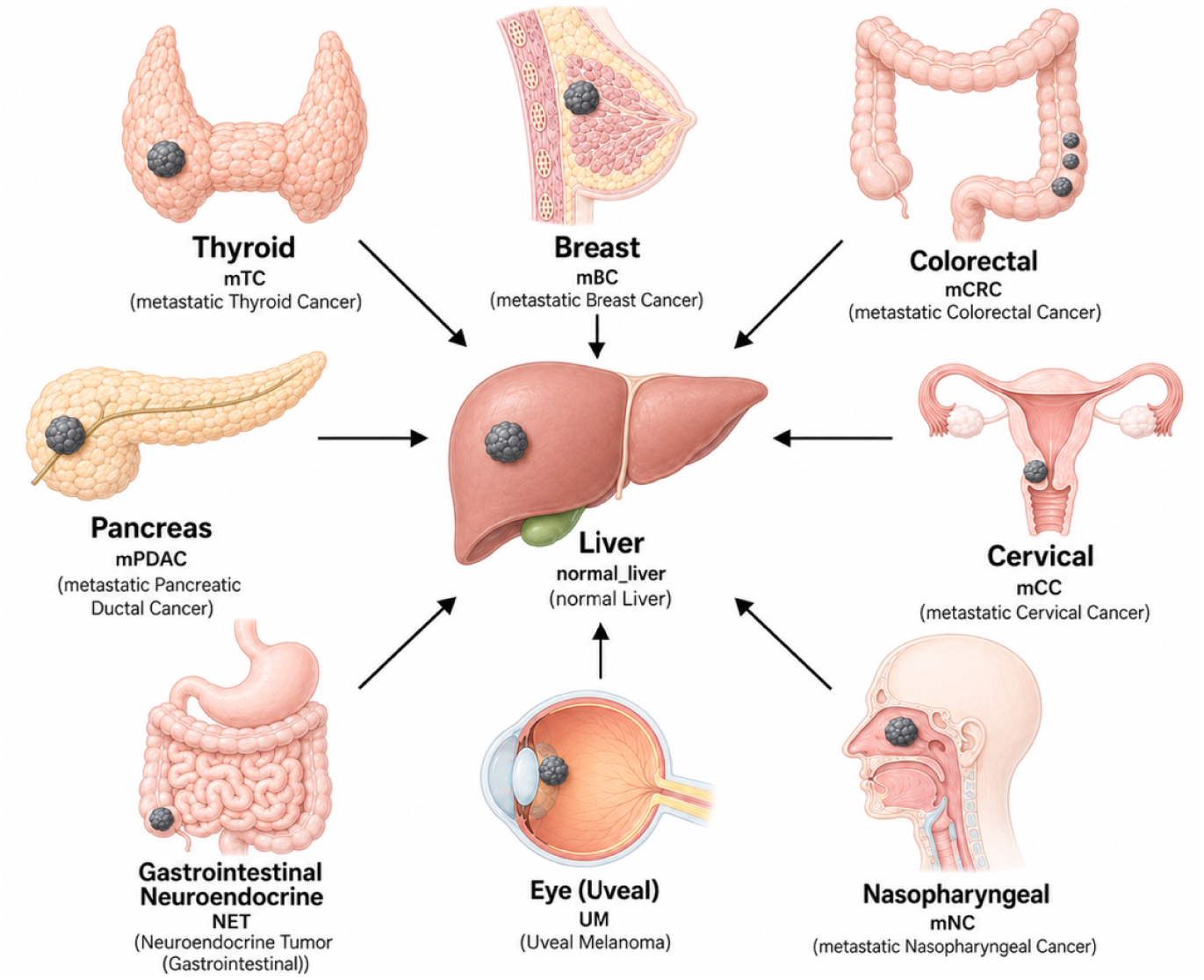
Tissue composition of the multi-study liver metastasis single-cell atlas. Schematic overview of the tissue sources incorporated into the atlas, encompassing primary and metastatic malignancies from multiple organs with a particular focus on tumors metastasizing to the liver. The integrated atlas includes metastatic thyroid cancer (mTC), metastatic breast cancer (mBC), metastatic colorectal cancer (mCRC), metastatic pancreatic ductal adenocarcinoma (mPDAC), metastatic cervical cancer (mCC), metastatic nasopharyngeal cancer (mNC), neuroendocrine tumors of gastrointestinal origin (NET), and uveal melanoma (UM), together with normal liver tissue from the liver reference atlas. Arrows indicate the integration of these diverse tissue sources into a unified liver-centered single-cell atlas for the characterization of cellular and transcriptional heterogeneity across primary and metastatic disease contexts.

**Figure S2.**
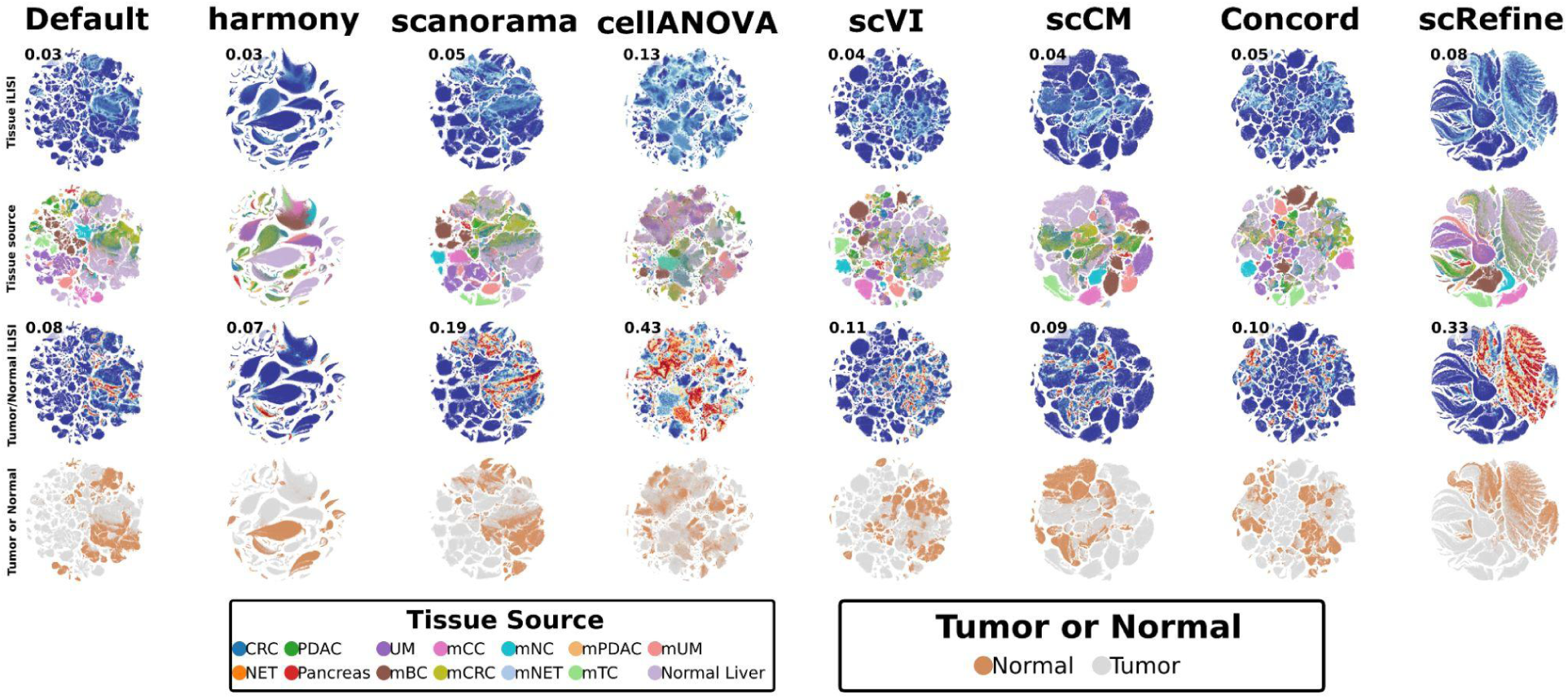
Comparative evaluation of tissue-source integration and tumor–normal cellular mixing across single-cell integration methods. Comparison of the uncorrected embedding (Default) with seven integration approaches—Harmony, Scanorama, cellANOVA, scVI, scCM, Concord, and scRefine—across the integrated liver metastasis single-cell transcriptomic atlas. The figure evaluates the extent of tissue-source mixing and the preservation of tumor–normal transcriptional differences using local inverse Simpson’s index (iLISI)-based measurements and complementary cellular annotations. The first row illustrates local mixing among different tissue sources, with the corresponding mean tissue-source iLISI-derived score displayed above each embedding. The second row shows the same cellular projections colored according to tissue or disease origin, including colorectal cancer (CRC), metastatic colorectal cancer (mCRC), neuroendocrine tumors (NET), metastatic neuroendocrine tumors (mNET), pancreas-derived samples, pancreatic ductal adenocarcinoma (PDAC), metastatic pancreatic ductal adenocarcinoma (mPDAC), uveal melanoma (UM), metastatic uveal melanoma (mUM), additional metastatic cancer categories (mNC, mBC, mCC, and mTC), and normal liver reference tissue. Among the evaluated methods, cellANOVA exhibits the highest tissue-source mixing score (0.13), followed by scRefine (0.08), whereas Scanorama and Concord achieve 0.05, scVI and scCM achieve 0.04, and both the Default and Harmony representations exhibit the lowest values (0.03). These results indicate method-dependent differences in the intermixing of cells originating from distinct tissue sources, with cellANOVA and scRefine showing comparatively greater tissue-source integration. The third row evaluates local mixing between tumor-associated and normal cells using tumor–normal iLISI-derived scores, while the fourth row displays the corresponding binary tumor–normal annotations, with normal cells shown in orange and tumor-associated cells in light gray. cellANOVA exhibits the highest tumor–normal mixing score (0.43), followed by scRefine (0.33) and Scanorama (0.19). Lower values are observed for scVI (0.11), Concord (0.10), scCM (0.09), Default (0.08), and Harmony (0.07). The elevated tumor–normal mixing observed with cellANOVA and scRefine indicates greater local intermixing of tumor-associated and normal cellular populations in their respective embedding spaces. Notably, although scRefine exhibits lower tumor–normal mixing than cellANOVA, its value remains substantially higher than that of the uncorrected representation, indicating that tumor–normal neighborhood organization is also affected by integration. Together, these results demonstrate distinct trade-offs between tissue-source integration and tumor–normal cellular separation across the evaluated methods. cellANOVA achieves the highest mixing scores for both tissue source and tumor–normal identity, whereas scRefine exhibits the second-highest scores for both comparisons. These findings highlight the importance of evaluating tissue-source mixing alongside the retention of biologically meaningful cellular heterogeneity, as increased mixing between tumor and normal populations may reflect either improved alignment of shared cellular identities or attenuation of genuine disease-associated transcriptional differences.

**Figure S3:**
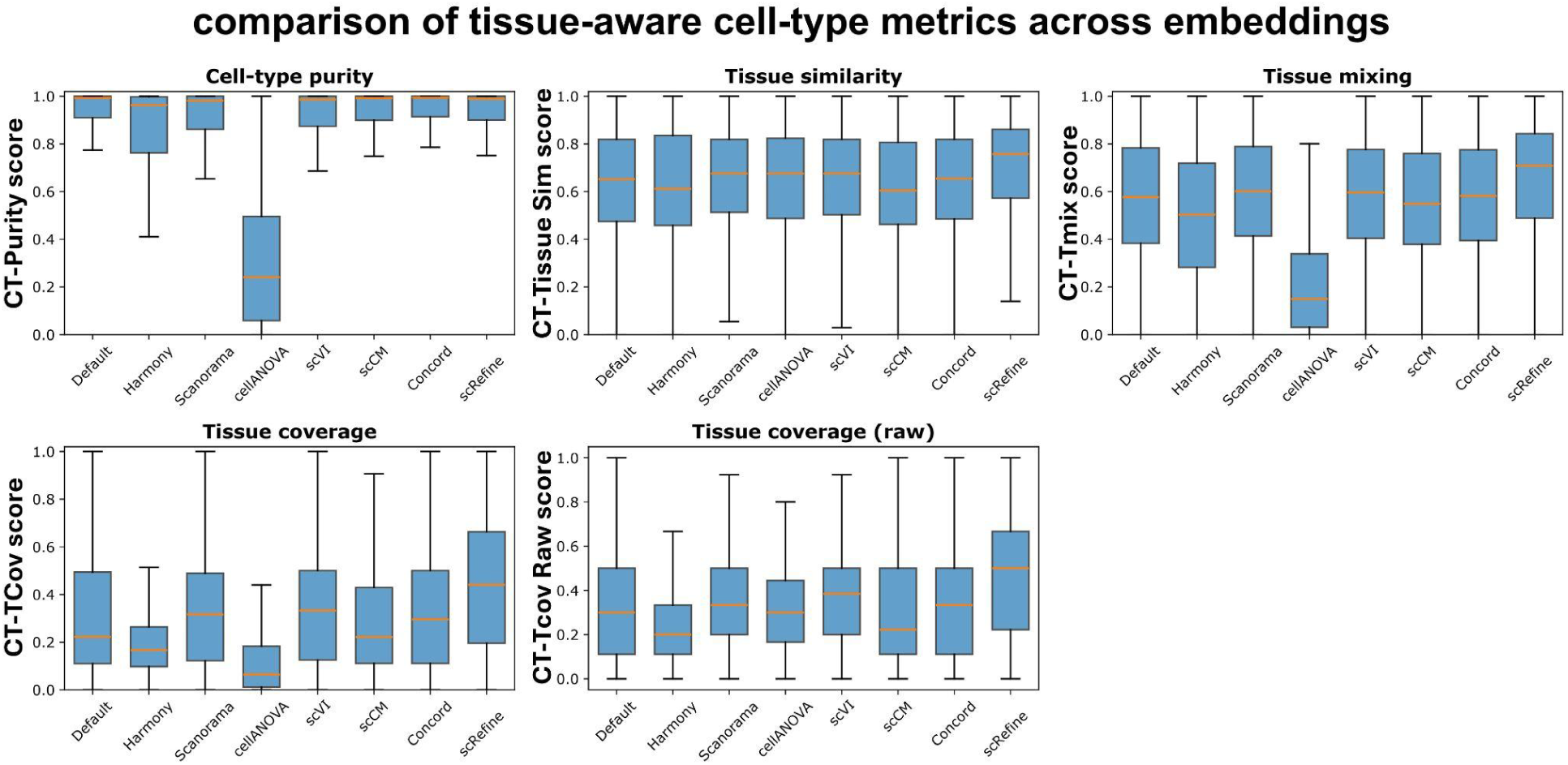
Comparative assessment of cell-type preservation and tissue-aware neighborhood mixing across single-cell integration methods. Boxplots illustrate the distributions of five complementary cell-type- and tissue-aware neighborhood metrics across eight integration methods: Default (PCA), Harmony, Scanorama, cellANOVA, scVI, scCM, Concord, and scRefine. The analysis was performed on the integrated multi-tissue cancer atlas using the corrected PanHuman Azimuth cell-type annotations as the biological reference. For each integration method, neighborhood relationships were independently calculated in the corresponding high-dimensional latent representation using 90 nearest neighbors and a neighborhood-weighting perplexity of 30. Calculations were restricted to cells with resolved cell-type annotations and valid tissue metadata. Approximately 630,400 cells contributed finite scores to each displayed metric for each method. Cell-type purity (CT-Purity; upper left) quantifies the proportion of distance-weighted neighborhood probability assigned to cells sharing the focal cell’s annotated identity and originating from tissues in which that cell type is expected to occur. Higher values indicate stronger local preservation of cell-type identity within the expected tissue context, whereas lower values indicate greater neighborhood mixing with unrelated cell types or cells from non-eligible tissue sources. Most methods exhibit high median purity, with Default, scVI, scCM, Concord, and scRefine retaining median values close to 1.0. In contrast, cellANOVA shows markedly reduced purity and a substantially broader distribution, indicating reduced preservation of cell-type-specific neighborhood structure under this metric. Tissue similarity (CT-TissueSim; upper middle) evaluates how closely the tissue-source distribution among same-cell-type neighbors matches the expected tissue distribution for that cell type across the atlas. Similarity is calculated as one minus the Jensen–Shannon divergence between the observed local and expected tissue-frequency distributions, producing values between 0 and 1. Higher values indicate stronger agreement with the expected tissue representation, rather than indiscriminate mixing across all tissues. scRefine exhibits the highest median tissue similarity (approximately 0.76), suggesting that its local neighborhoods more closely reproduce the tissue distributions expected for the corresponding cell types. For cell types associated with only one eligible tissue, similarity is defined as 1 when matching neighborhood probability is present. Tissue mixing (CT-TMix; upper right) combines cell-type preservation and tissue-distribution similarity into a single composite measure, calculated as CT-Purity multiplied by CT-TissueSim. This formulation rewards neighborhoods that preserve cell-type identity while maintaining tissue representation consistent with the expected biological distribution. Consequently, high tissue diversity alone does not yield a high score if the neighborhood mixes unrelated cell types. scRefine demonstrates the highest median tissue-mixing score (approximately 0.71), compared with approximately 0.50 -- 0.60 for most baseline methods. cellANOVA displays markedly lower scores, reflecting the strong contribution of its reduced cell-type purity to the combined metric. Tissue coverage (CT-TCov; lower left) measures the representation of expected tissue sources within the local neighborhood while accounting for cell-type preservation. For each cell type, eligible tissues were defined from the atlas-wide cell-type-by-tissue frequency distribution, requiring at least 30 cells and at least 0.5% of the annotated population of that cell type. Local coverage is determined by the fraction of eligible tissue categories represented by at least one same-cell-type neighbor and is multiplied by CT-Purity to obtain the adjusted coverage score. Higher values therefore indicate broader representation of expected tissues within neighborhoods that retain the focal cell’s identity. scRefine achieves the highest median adjusted tissue coverage (approximately 0.44), while cellANOVA shows the lowest median coverage. Raw tissue coverage (CT-TCovRaw; lower middle) represents the unadjusted fraction of eligible tissue categories detected among same-cell-type neighbors, without multiplying by cell-type purity. This metric distinguishes the breadth of local tissue representation from the degree of cell-type preservation. scRefine exhibits the highest median raw tissue coverage (approximately 0.50), followed by several baseline approaches with intermediate coverage. Differences between raw and adjusted coverage highlight the importance of evaluating tissue representation alongside cell-type-specific neighborhood structure, since broader tissue representation can occur even in neighborhoods with poor cell-type preservation. Boxplot interpretation and benchmarking considerations. Each box represents the interquartile range (25th–75th percentiles) of the per-cell metric distribution, with the central orange line indicating the median. Whiskers extend to the most extreme observations within 1.5 times the interquartile range; individual observations beyond the whiskers are omitted from visualization. All five metrics are represented on a common 0–1 scale, with higher scores indicating greater agreement with the specific preservation, similarity, or coverage criterion being evaluated. Scores were calculated independently from each method’s high-dimensional embedding, rather than from two-dimensional t-SNE or UMAP projections. Collectively, these results reveal substantial method-dependent differences in the balance between cell-type preservation and tissue-aware neighborhood organization. scRefine maintains high cell-type purity while showing comparatively greater tissue-distribution similarity, tissue mixing, and tissue coverage. In contrast, cellANOVA exhibits broad tissue-similarity distributions but substantially reduced cell-type purity, leading to low composite mixing and adjusted coverage scores. These findings support the importance of jointly evaluating local cell-type conservation and biologically expected tissue representation when comparing integration methods. Because expected tissue frequencies are estimated from the atlas itself and may reflect tissue sampling imbalance, the metrics should be interpreted as measures of neighborhood organization relative to the observed reference distribution, rather than as independent evidence of biological correctness. No statistical significance testing is depicted, and individual cells should not be treated as independent biological replicates.

**Figure S4:**
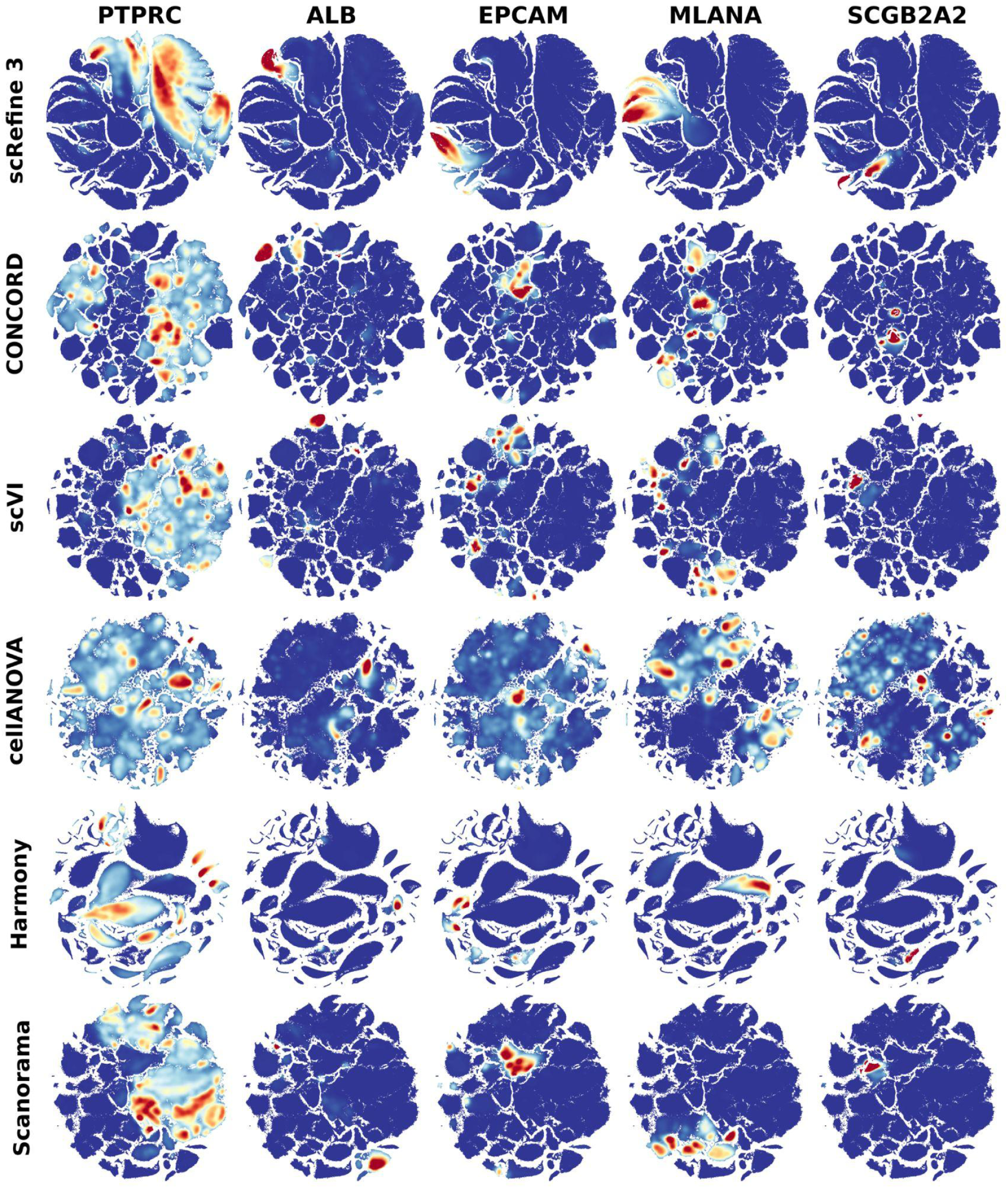
Preservation of lineage-specific gene-expression patterns across integration methods. Gene-expression density maps for representative canonical markers projected onto embeddings generated using scRefine, Concord, scVI, cellANOVA, Harmony, and Scanorama. Markers represent major cellular compartments within the integrated liver metastasis atlas, including PTPRC for immune cells, ALB for hepatocytes, EPCAM for epithelial cells, MLANA for melanocytic/melanoma cells, and SCGB2A2 for breast epithelial/tumor-associated cells. Warmer colors indicate regions of higher expression density. The comparison illustrates differences among integration approaches in their ability to preserve spatially coherent, lineage-specific transcriptional signals within the integrated latent space.

**Figure S5:**
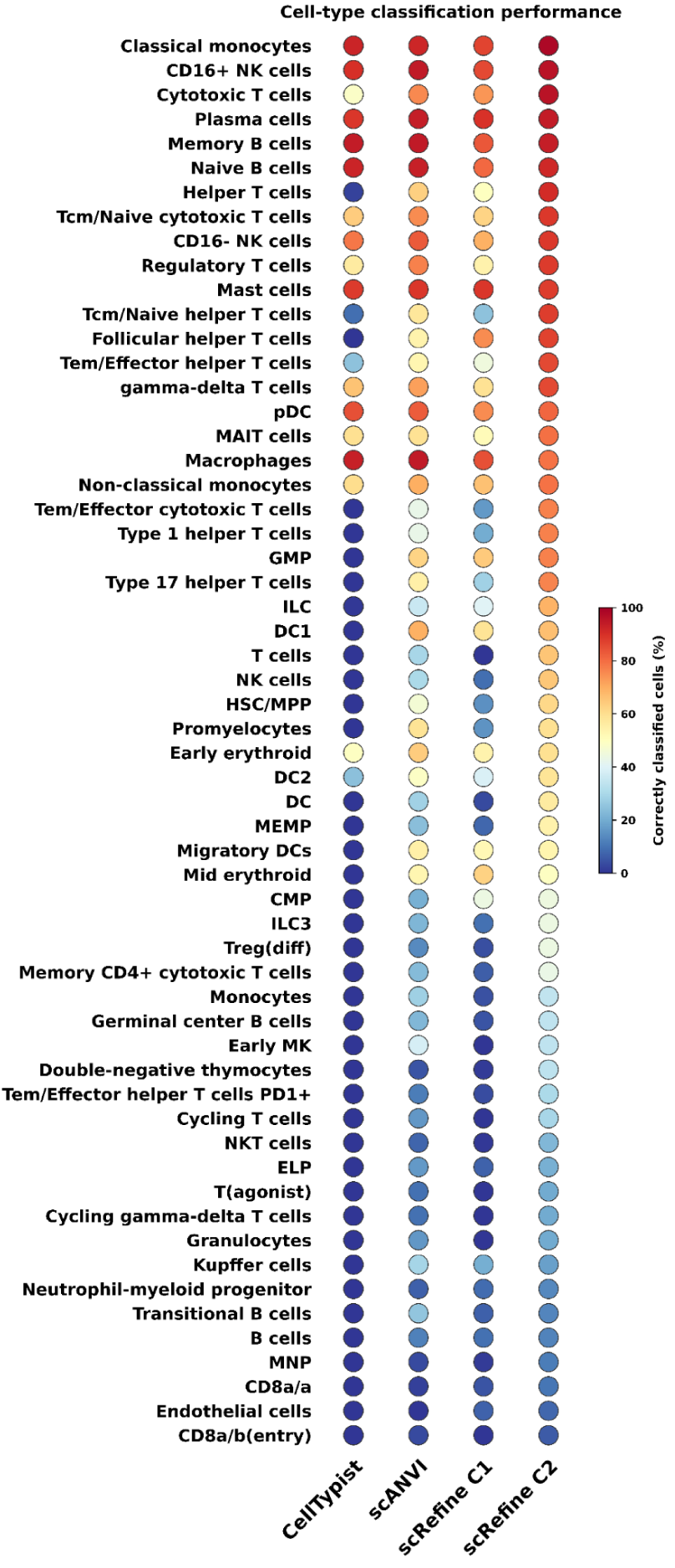
Cell-type–specific classification performance across annotation approaches. Dot plot comparing the proportion of correctly classified cells across CellTypist, scANVI, scRefine with NN based classification, and scRefine with contrastive k-means for individual cell types. Dot colour represents the percentage of correctly classified cells, ranging from low (blue) to high (red) classification performance. scRefine C2 generally achieves higher classification accuracy across a broad range of immune and progenitor cell populations, while performance varies between methods for more closely related or fine-grained cell states.

**Figure S6:**
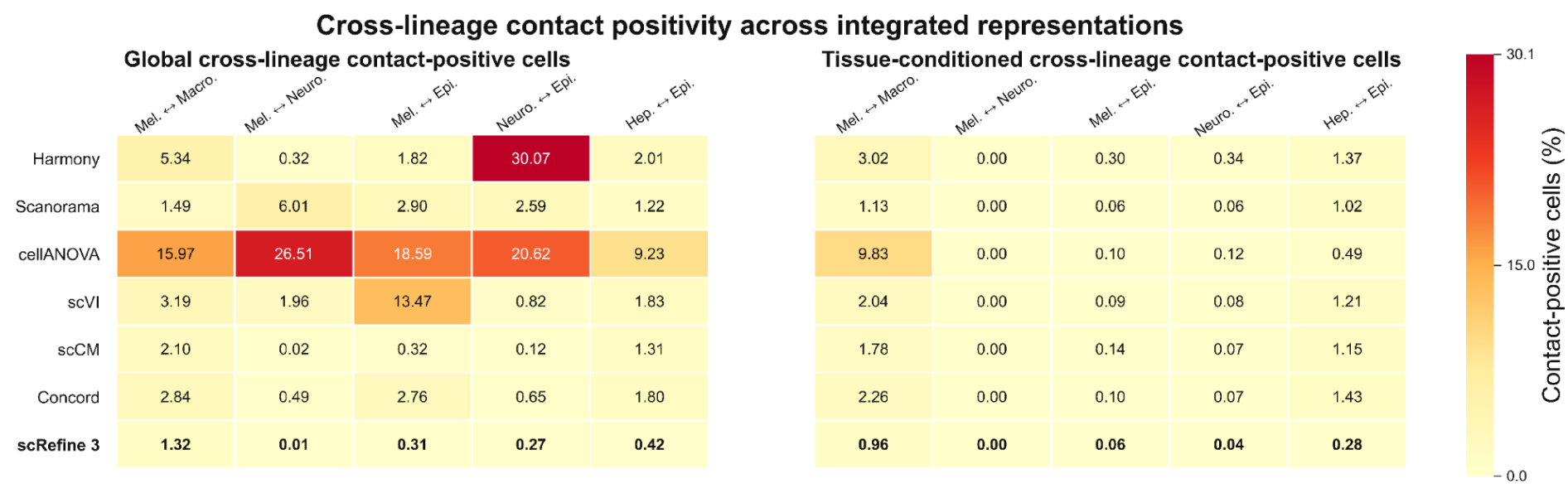
scRefine preserves cross-lineage separation across integrated representations. Heatmaps show the percentage of cells classified as contact-positive for each prespecified cross-lineage pair across integrated embeddings. A cell was considered contact-positive when at least 3 of its 30 nearest neighbours belonged to the paired opposite lineage. Left, global contact-positive fractions calculated across the full atlas. Right, tissue-conditioned fractions, requiring the opposite-lineage neighbour to share the focal cell’s tissue annotation. Values are calculated among cells eligible for each lineage pair; darker shading indicates a higher fraction of cross-lineage contacts. scRefine showed consistently low global and tissue-conditioned contact positivity across all evaluated lineage pairs, supporting preservation of distinct cellular neighbourhood structure.

**Figure S7:**
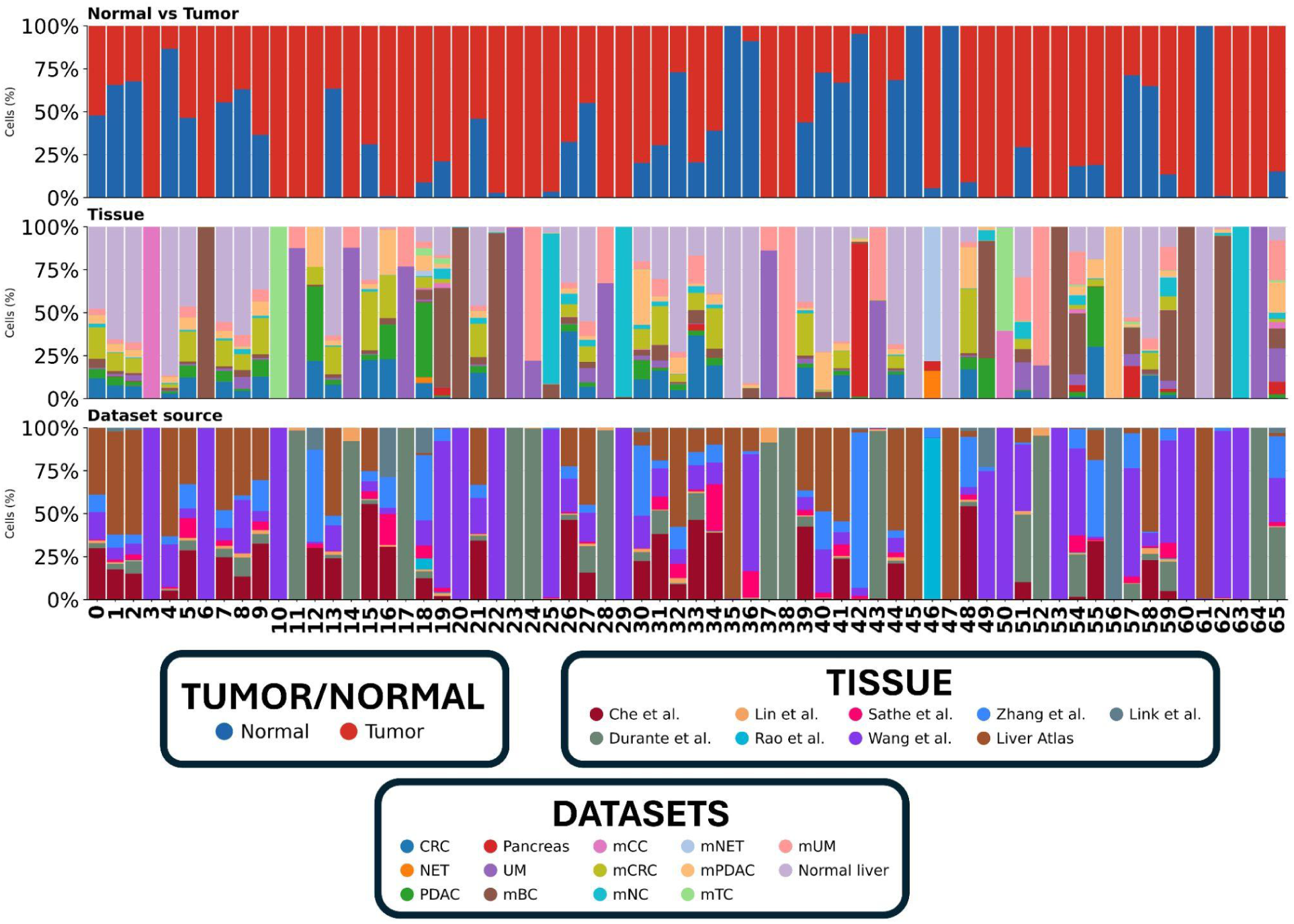
Composition of scRefine-derived cellular clusters across datasets, tissue sources, and disease states. Stacked bar plots illustrate the relative cellular composition of 66 Leiden clusters (0–65), identified using the scRefine embedding at a clustering resolution of 1.5. Each column represents an individual Leiden cluster, with the proportion of cells normalized to 100% to facilitate comparisons across clusters of different sizes. The **upper panel** shows the distribution of cells classified as normal (blue) or tumor-associated (red), highlighting clusters enriched for either category and clusters containing cells from both conditions. The **middle panel** illustrates the tissue or disease origin of cells within each cluster, encompassing colorectal cancer (CRC), neuroendocrine tumors (NET), pancreatic ductal adenocarcinoma (PDAC), pancreas-derived samples, uveal melanoma (UM), metastatic breast cancer (mBC), metastatic cervical cancer (mCC), metastatic colorectal cancer (mCRC), metastatic neuroendocrine tumors (mNET), metastatic pancreatic ductal adenocarcinoma (mPDAC), metastatic uveal melanoma (mUM), and additional metastatic cancer categories (mNC and mTC), alongside normal liver samples. The **lower panel** depicts the relative contributions of the individual source studies, including Che et al., Durante et al., Lin et al., Link et al., Rao et al., Sathe et al., Wang et al., Zhang et al., and the Liver Atlas reference [3–10]. Together, these distributions reveal heterogeneity in tumor–normal composition, tissue enrichment, and study representation across transcriptionally defined clusters. Clusters containing contributions from multiple tissue and dataset sources suggest the presence of transcriptionally shared cellular populations, whereas clusters dominated by individual sources may reflect tissue-specific biology, differences in cellular composition, or residual study-associated effects. The figure provides a complementary assessment of cellular organization and source composition following integration with scRefine.

**Figure S8:**
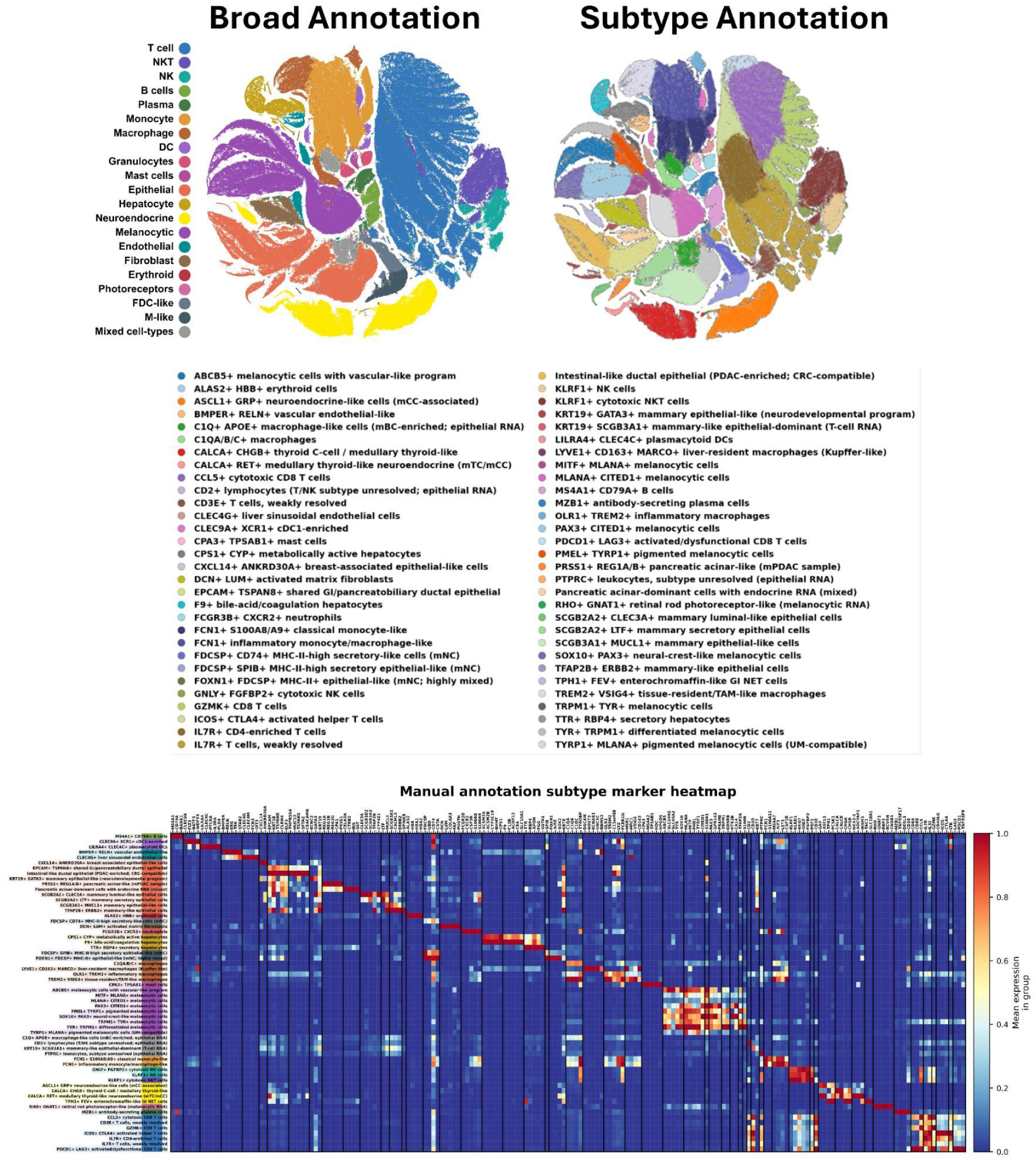
Differentially expressed gene signatures defining annotated cellular subtypes in the scRefine-integrated atlas. Heatmap showing the mean expression of representative differentially expressed genes (DEGs) across annotated cell subtypes identified within the scRefine latent space. Rows represent annotated cellular subtypes and transcriptional states, while columns correspond to subtype-associated marker genes. Expression values are scaled within each gene, with warmer colors indicating higher mean expression within a given subtype. Canonical lineage-specific and state-associated expression programs support the annotation of major immune, myeloid, epithelial, endothelial, hepatocyte, melanocytic, neuroendocrine, and stromal populations, as well as tumor-associated and stress-related cellular states. The lower panel shows the scRefine embedding colored by the corresponding subtype annotations, illustrating the spatial organization of transcriptionally distinct cell populations within the integrated atlas.

**Figure S9:**
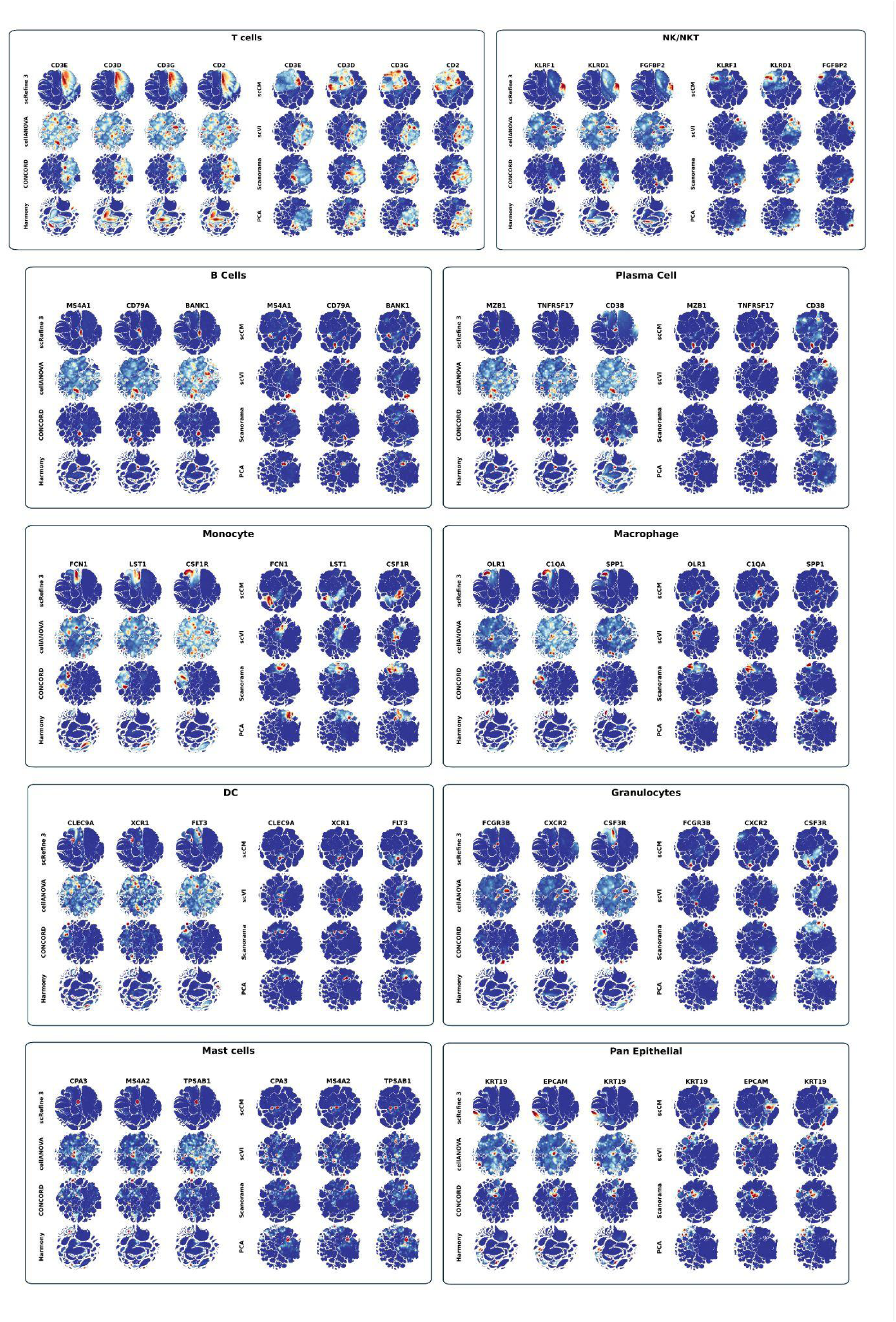
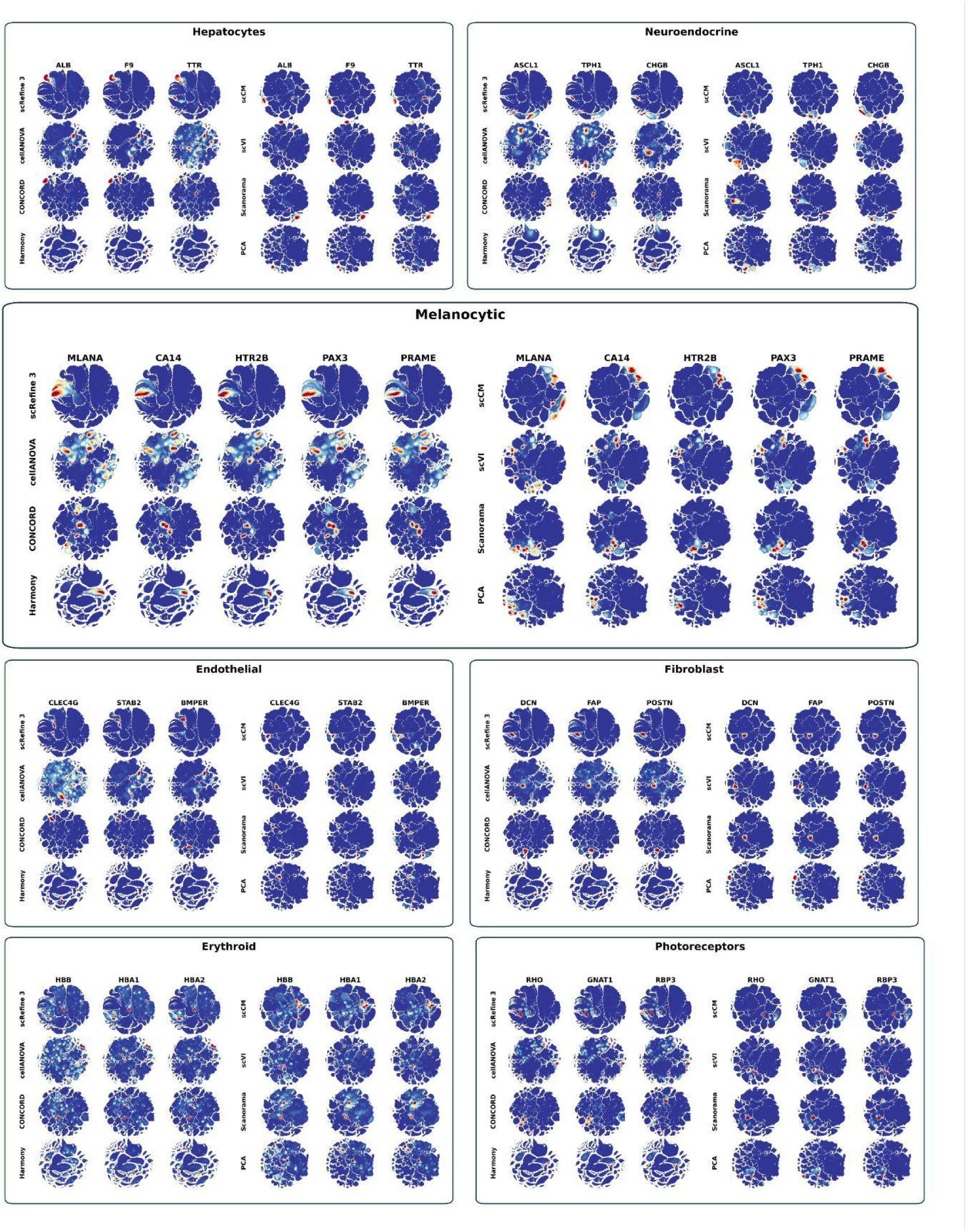
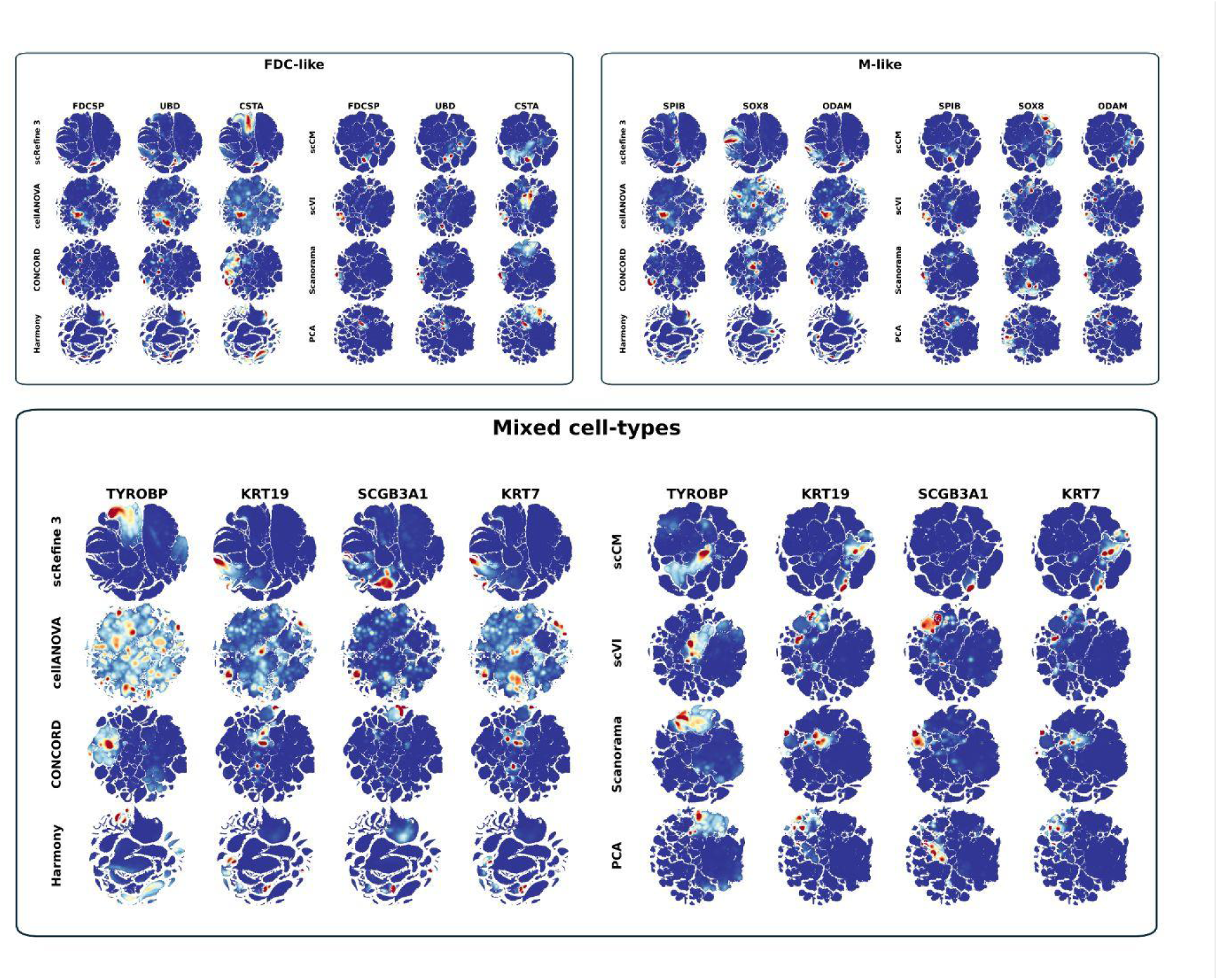
Comparison of canonical cell-type marker expression across integration methods. Feature plots show the expression of representative lineage and cell-state markers projected onto embeddings generated using scRefine, cellANOVA, Concord, Harmony, scCM, scVI, Scanorama, and PCA. Marker panels represent CD4/CD8 T cells (CD3E, CD3D, CD4, CD8A), tissue-resident macrophages (CD5L, MARCO, FOLR2), plasmacytoid dendritic cells (LILRA4, CLEC4C, PTCRA), SPP1+ TREM2 TAM-like macrophages (SPP1, MSR1, OLR1), NK cells (KLRF1, GNLY, FGFBP2), melanocyte/melanoma-like cells (MLANA, MITF, PMEL), mast cells (TPSAB1, MS4A2, CPA3), and hepatocytes (ALB, TTR, CPS1). Comparison of marker-expression patterns demonstrates method-dependent differences in the preservation of lineage-specific cellular neighbourhoods, with scRefine generally maintaining coherent localization of canonical marker programs while preserving transcriptional heterogeneity across major cell populations.

**Figure S10:**
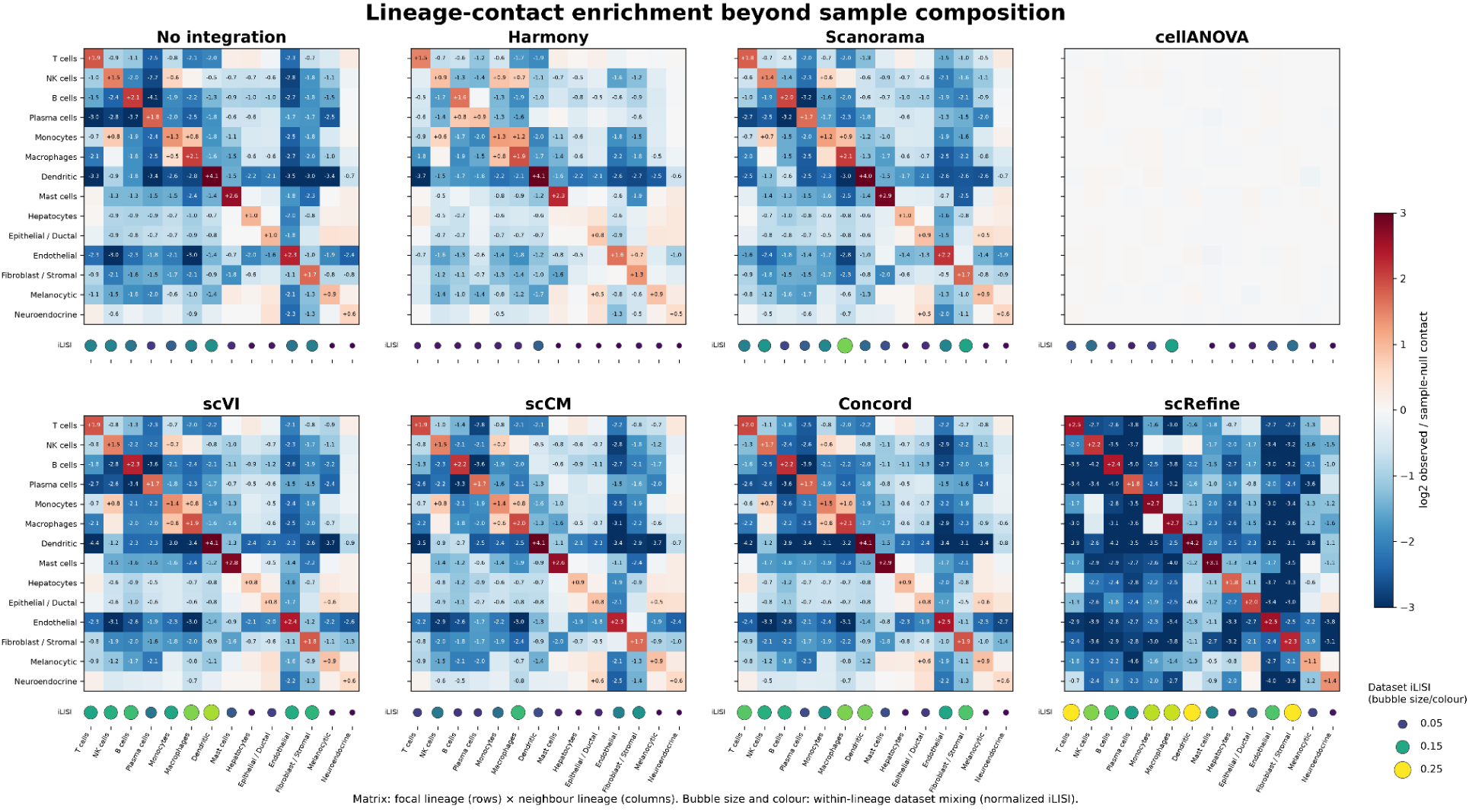
Normalized lineage-contact enrichment across integration methods. Heatmap values represent log₂-observed-to-expected contact frequencies among 30 nearest neighbours, relative to a sample-preserving compositional null. Red indicates enriched and blue indicates depleted contacts. Bubble size and colour show normalized dataset iLISI within each lineage, with larger bubbles indicating greater cross-dataset mixing. cellANOVA showed evidence of over-integration in this analysis, with reduced preservation of lineage-specific neighbourhood structure. This may partly reflect our integration design, in which the normal liver atlas was explicitly included as the normal reference, potentially encouraging stronger correction of disease-associated variation by cellANOVA.

**Figure S11:**
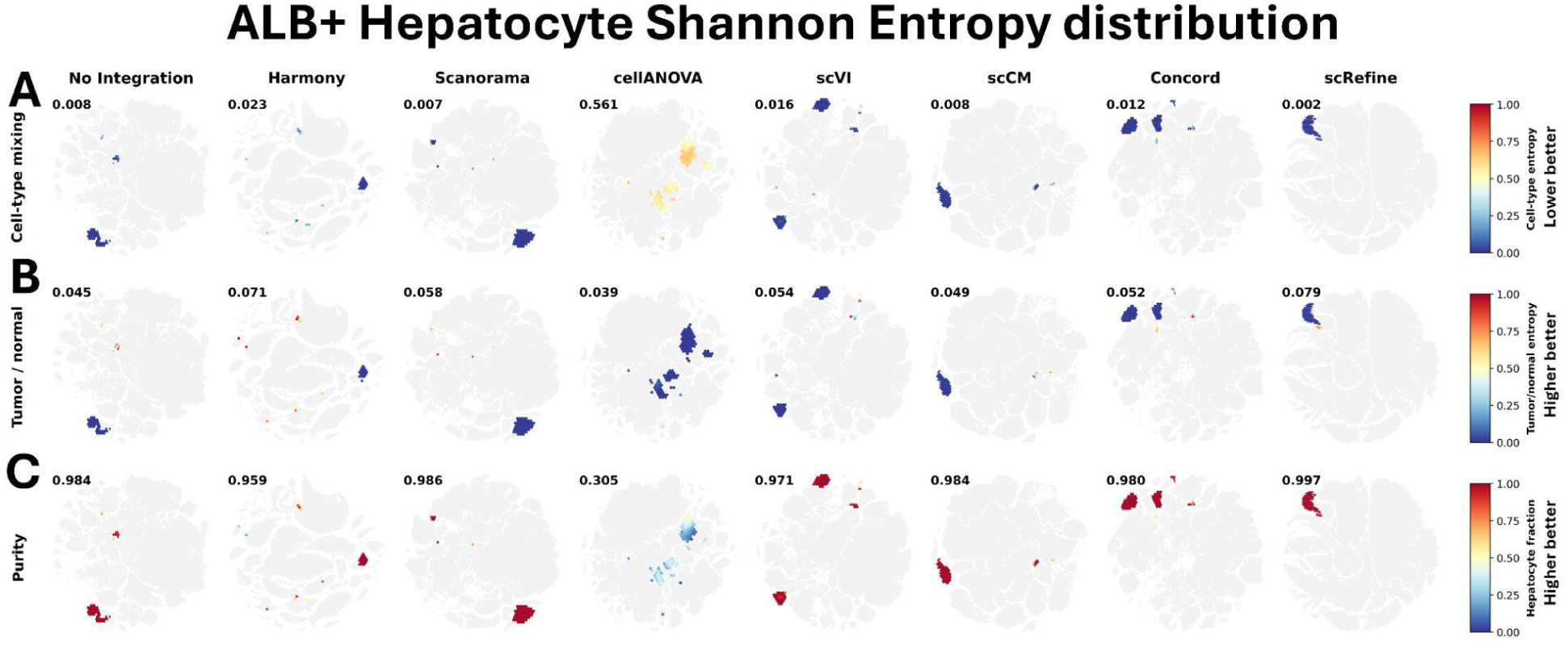
t-SNE visualization of ALB-high cells across eight integration embeddings. Gray indicates the complete background, while colored regions represent ALB-high cells. Values in the upper-left corner are the mean metric values for each embedding. **A. cell-type mixing**: low values indicate that nearby cells are mostly of one cell type, whereas high values indicate a diverse and relatively balanced mixture of cell types. **B. tumor/normal mixing**: low values indicate that normal-liver and liver-metastasis cells occupy separate neighborhoods, while high values indicate that they are locally intermixed. **C. hepatocyte-neighborhood purity**: high values indicate that ALB-high cells are surrounded mainly by hepatocytes, whereas low values indicate neighborhoods dominated by other cell types.

**Figure S12:**
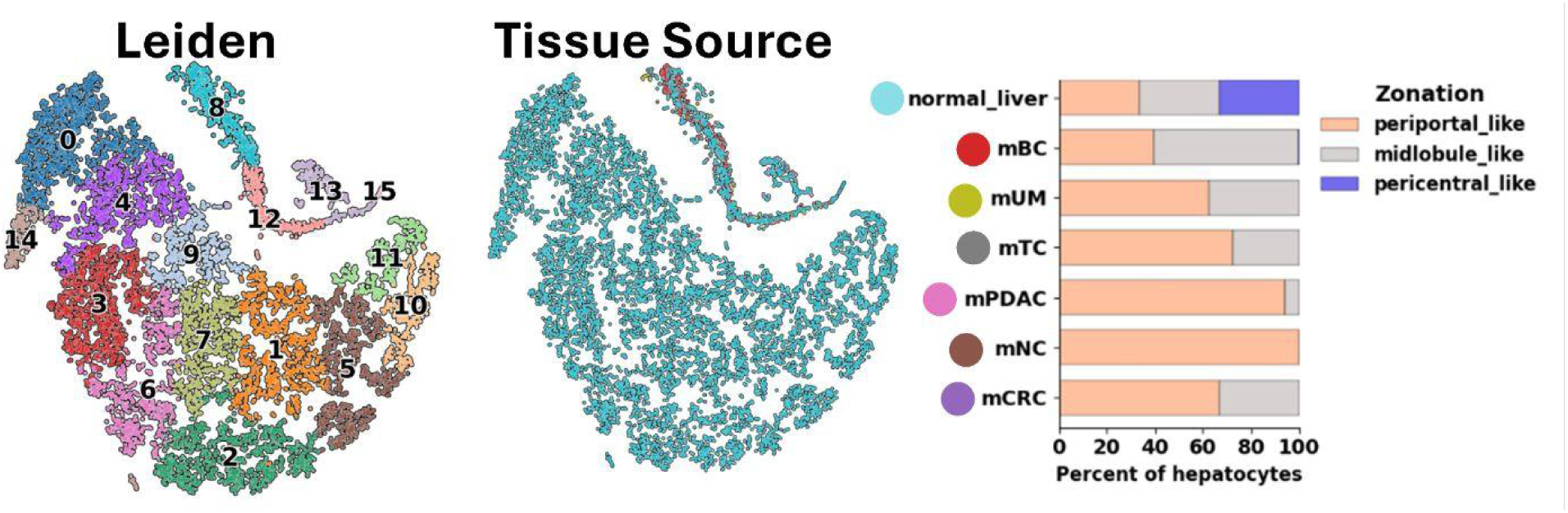
Hepatocyte re-clustering and zonation profiles across tissue sources. Hepatocyte-specific re-clustering identified multiple transcriptionally distinct Leiden subclusters within the integrated atlas (left). ALB expression across the hepatocyte embedding supports the hepatic identity of the re-clustered cells (middle). Stacked bar plots show the relative proportions of periportal-like, midlobular-like, and pericentral-like hepatocyte states across normal liver and metastatic tissue sources (right). Normal liver displays a broader distribution of zonation states, whereas hepatocytes associated with metastatic tissues are predominantly enriched for periportal-like transcriptional programs, indicating remodeling of the physiological hepatocyte zonation landscape in the metastatic liver microenvironment.

**Figure S13:**
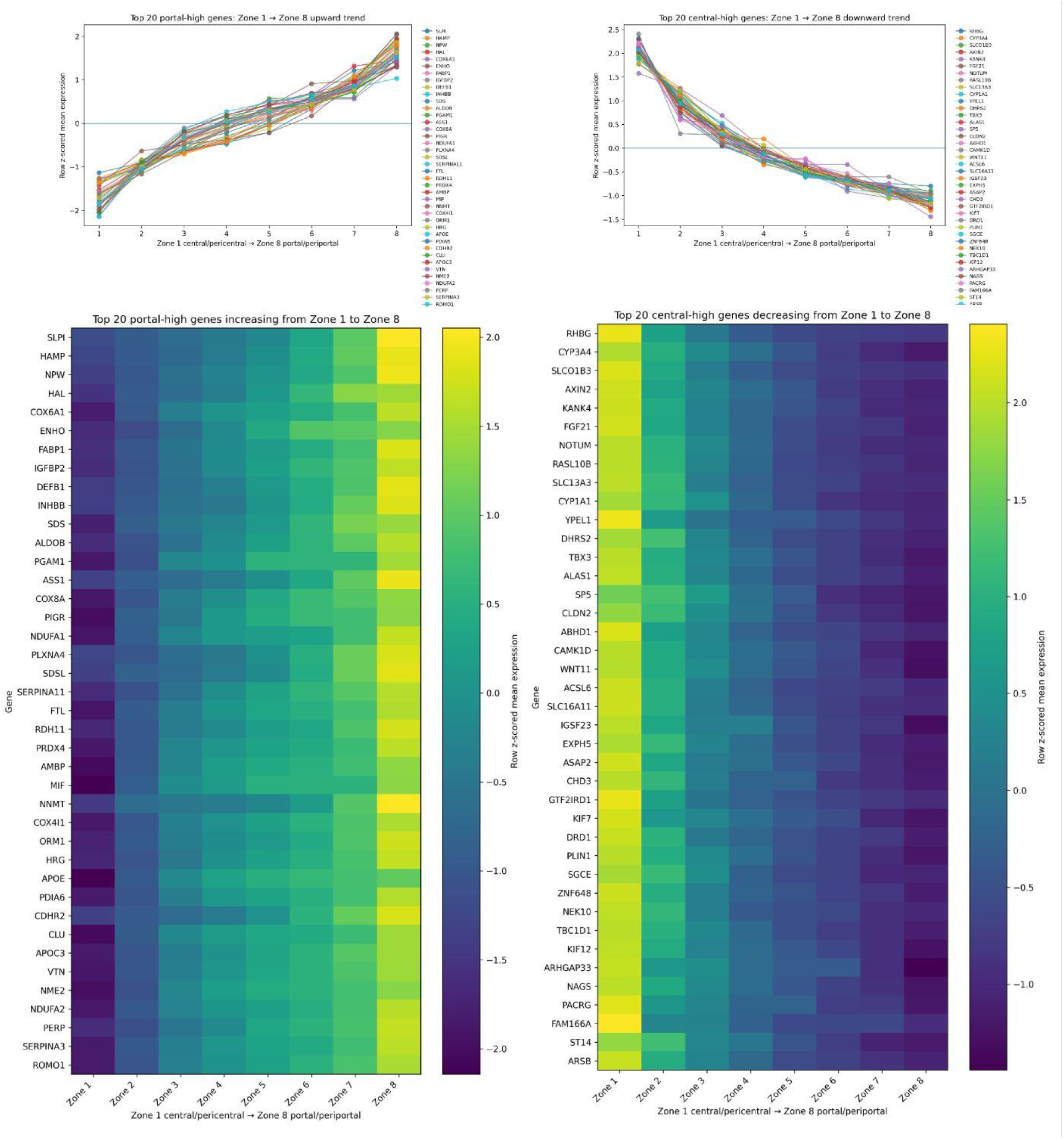
Reconstruction of hepatocyte zonation using reference gene signatures from the healthy human liver spatial atlas. Hepatocyte zonation was inferred using central/pericentral- and portal/periportal-associated gene-expression trends derived from the zonation analysis reported by Yakubovsky *et al.* [54] in the healthy human liver spatial atlas. The upper panels show the mean z-scored expression trajectories of the top central-high genes (left) and portal-high genes (right) across eight ordered zonation bins, progressing from Zone 1 (central/pericentral) to Zone 8 (portal/periportal). The corresponding heatmaps (lower panels) visualize gene-specific expression gradients across the same zonation axis. Central-associated genes progressively decrease from Zone 1 to Zone 8, whereas portal-associated genes show the reciprocal increase, supporting reconstruction of a continuous porto-central hepatocyte zonation gradient. The reference study established hepatocyte zonation along the porto-central axis using spatially resolved landmark gene expression in healthy human liver.

**Figure S14:**
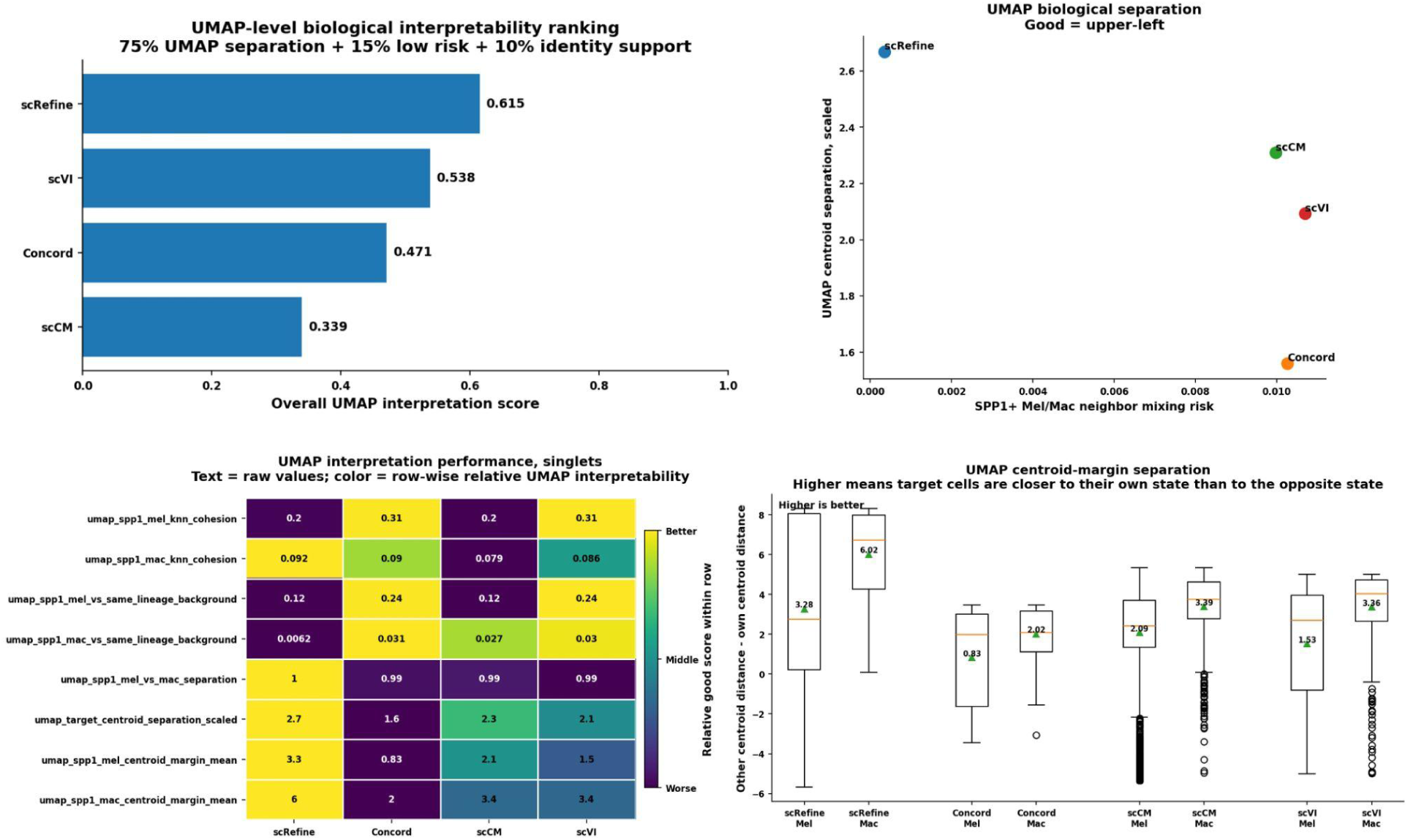
UMAP-level separation of SPP1-high macrophages and SPP1-high melanocytic cells across integration methods. Comparative evaluation of scRefine, CONCORD, scCM, and scVI UMAPs based on their ability to distinguish SPP1-high macrophage and SPP1-high melanocytic populations. The upper-left panel shows the aggregate UMAP interpretability score, calculated using 75% biological separation, 15% low-risk cross-lineage mixing, and 10% cell-identity support. scRefine achieved the highest aggregate score (0.809), followed by scVI (0.385), CONCORD (0.319), and scCM (0.185). The upper-right panel compares scaled centroid separation with SPP1-high melanocyte–macrophage neighbourhood-mixing risk; favourable embeddings show greater vertical separation and lower horizontal mixing. The lower-left heatmap displays the raw component metrics, with colour representing the relative performance of each method within a metric. Metrics include SPP1-high melanocyte and macrophage kNN cohesion, same-lineage background mixing, scaled target-state centroid separation, and melanocyte and macrophage control margins. The lower-right violin plots show cell-level neighbourhood cohesion for the two SPP1-high target states. scRefine showed particularly strong SPP1-high macrophage cohesion (approximately 0.34) compared with CONCORD, scCM, and scVI (approximately 0.065–0.077), while SPP1-high melanocyte cohesion was more similar across methods. Overall, scRefine provided the strongest UMAP-level separation between the cellular origins of SPP1 expression while maintaining favourable within-lineage neighbourhood organisation.

**Figure S15:**
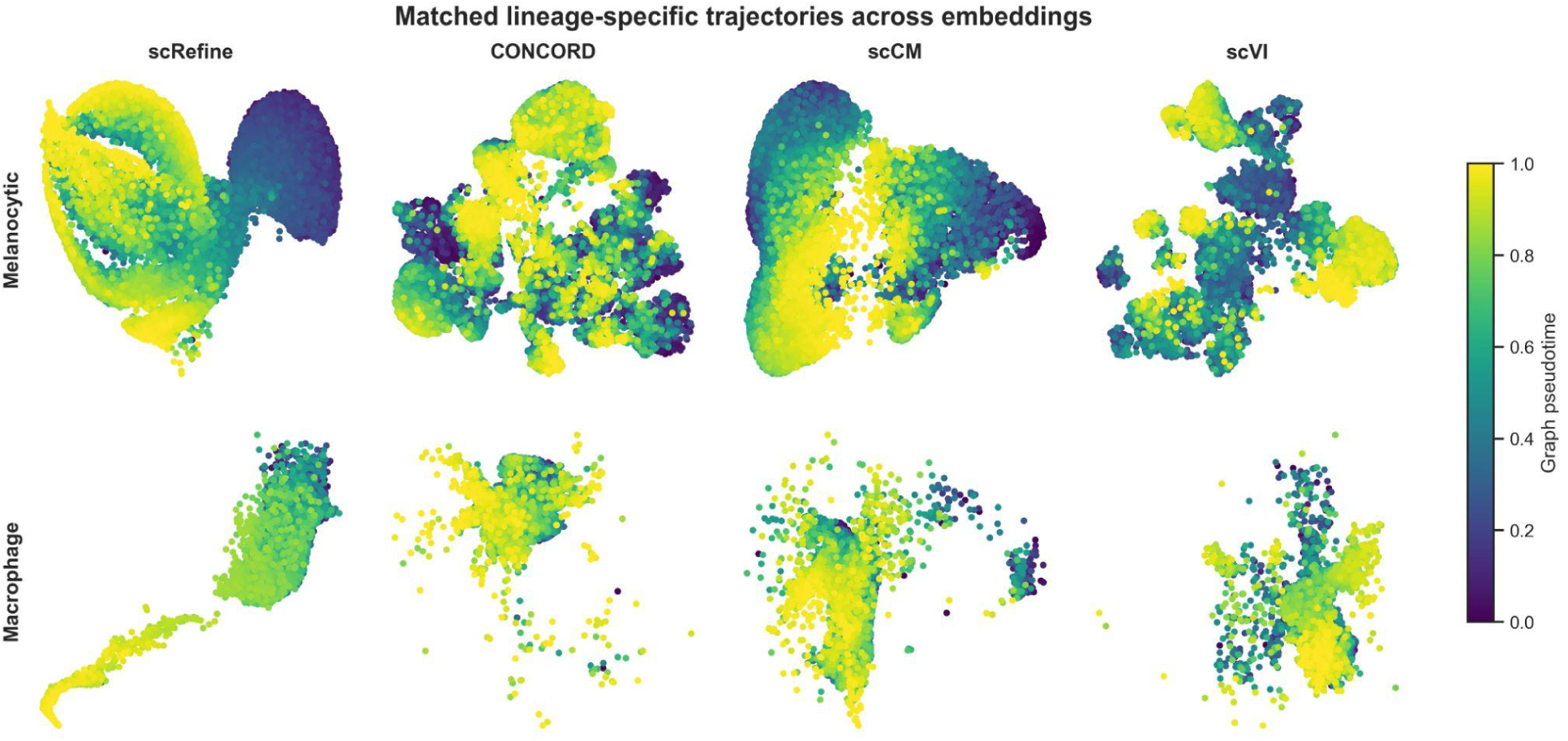
Matched lineage-specific primary-to-metastatic trajectories across integration embeddings. Method-specific UMAPs for scRefine, CONCORD, scCM, and scVI, coloured by graph pseudotime from 0 to 1. Melanocyte and macrophage trajectories were constructed independently using their corresponding high-dimensional latent embeddings. Primary, SPP1-low cells were used to orient the trajectory roots, with MLANA-high primary cells additionally used to define the melanocyte root population. The upper row shows the melanoma-cell trajectories, and the lower row shows the macrophage trajectories. The UMAPs provide two-dimensional visualisations of the latent-space pseudotime ordering and were not themselves used to calculate the trajectories. Quantitative trajectory comparisons are reported in Figure 4B.

**Figure S16:**
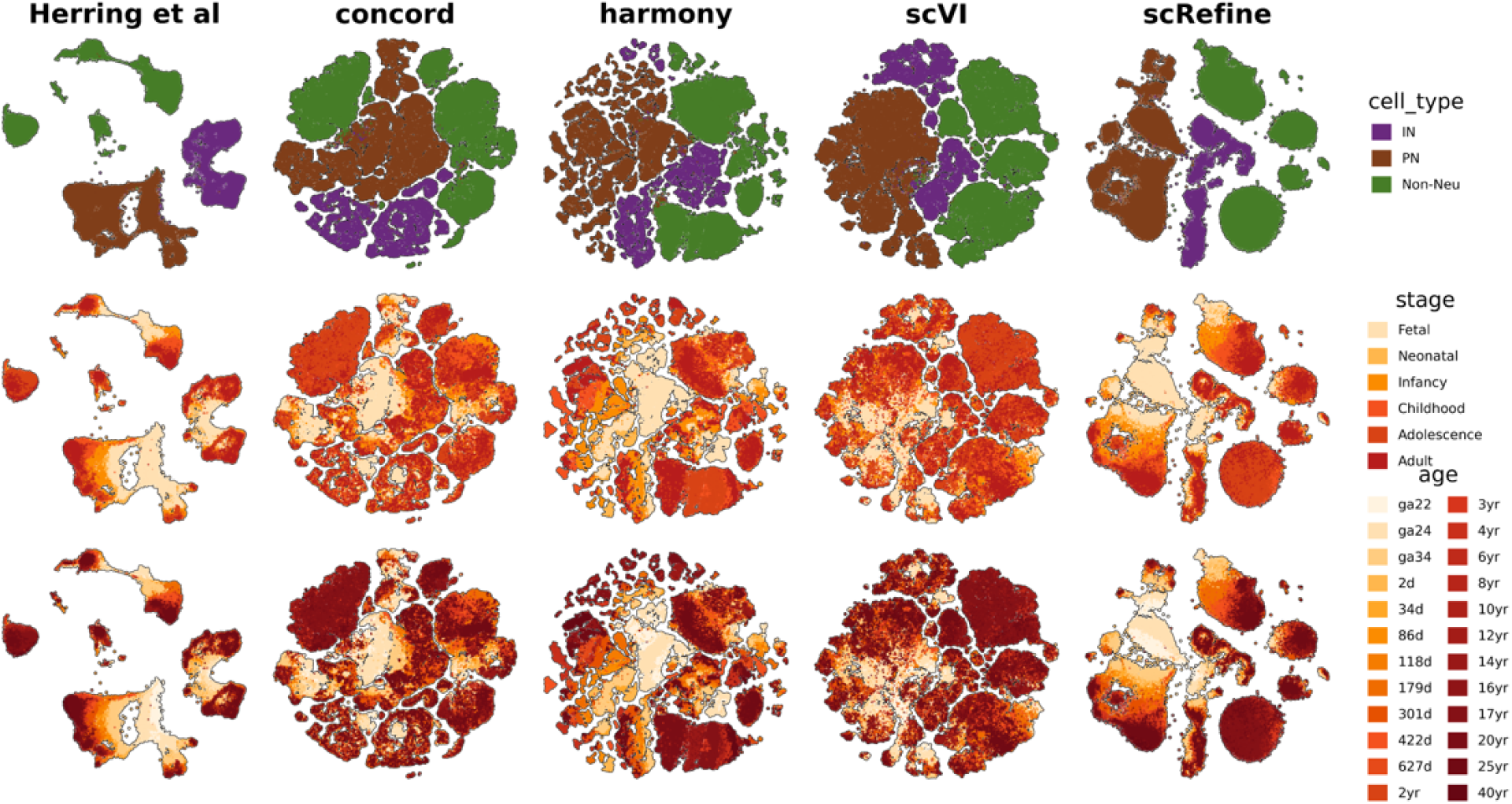
Preservation of developmental trajectories in the Herring et al. brain atlas across integration methods. Low-dimensional embeddings of the Herring et al. [1] developmental brain dataset are shown for the original representation and following integration with Concord, Harmony, scVI, and scRefine. Cells are colored by broad cell class (top), developmental stage (middle), and chronological age (bottom). scRefine most closely preserves the organization of the original dataset, maintaining separation of major cell classes together with continuous stage- and age-associated gradients. In comparison, alternative integration methods show greater fragmentation or redistribution of developmental states. These results indicate that scRefine better retains the underlying developmental trajectory and temporal structure of the reference atlas during integration.

**Figure S17:**
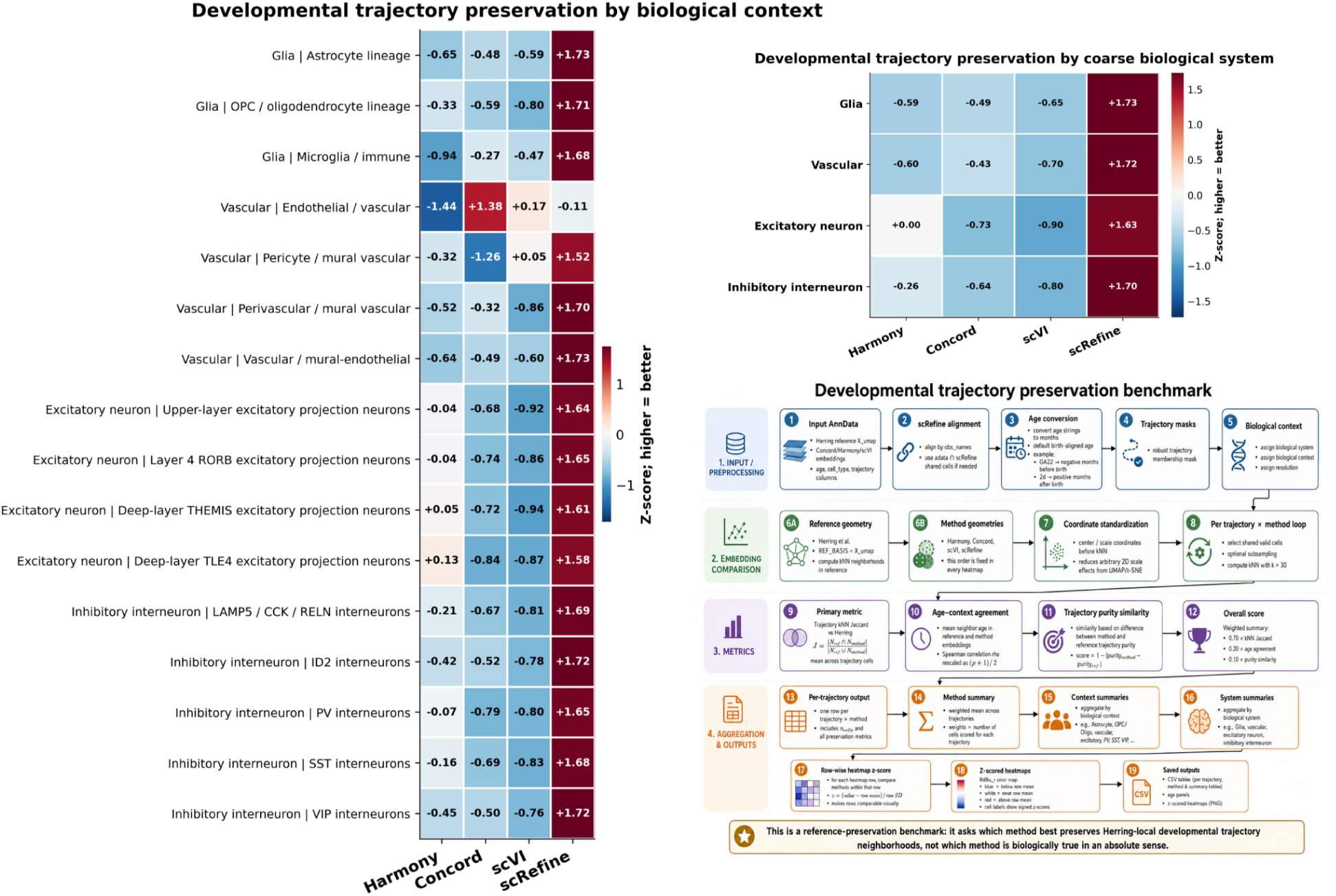
Benchmarking preservation of developmental trajectories across integration methods. Heatmaps summarize developmental trajectory preservation scores for Harmony, Concord, scVI, and scRefine across fine-grained biological contexts (left) and broader biological systems, including glial, vascular, excitatory neuronal, and inhibitory interneuron compartments (upper right). Scores are standardized as z-scores, with higher values indicating stronger preservation of the reference developmental trajectory. scRefine consistently achieves the highest trajectory-preservation scores across most biological contexts and coarse cell systems, indicating improved retention of age-associated and lineage-specific developmental structure. The lower-right schematic outlines the benchmarking framework used to compare each integrated embedding against the reference geometry, incorporating trajectory construction, age-context agreement, trajectory purity, and aggregation of performance across biological contexts.

**Figure S18:**
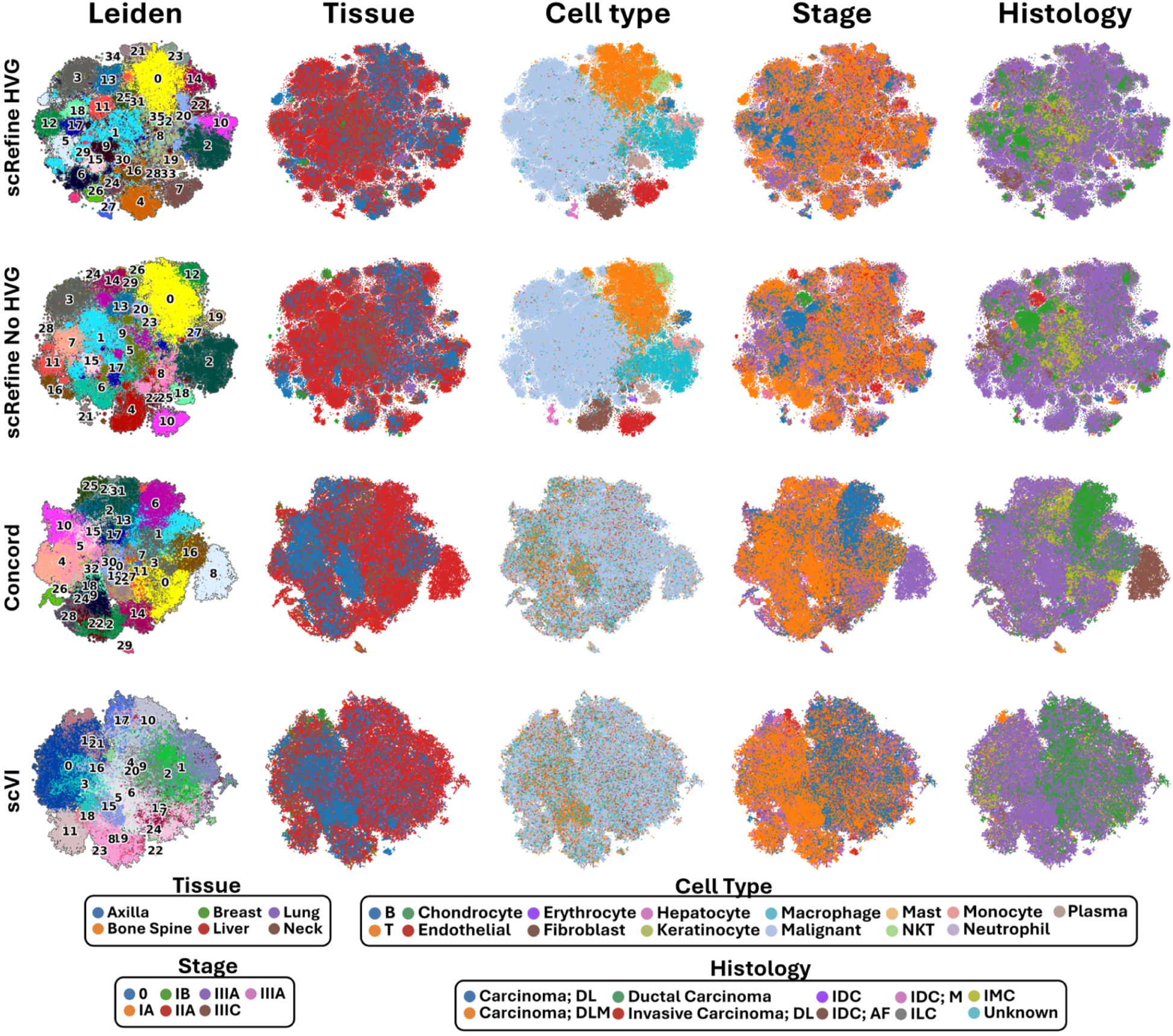
Preservation of author-provided cell-type and clinicopathological annotations in the metastatic breast cancer dataset across integration methods. Low-dimensional embeddings of the metastatic breast cancer single-cell dataset from Klughammer et al. [2] are shown following integration with scRefine using highly variable genes (HVGs), scRefine without HVG selection, Concord, and scVI. Cells are colored by Leiden cluster, tissue of origin, cell type, disease stage, and histological classification. Cell-type annotations were retained directly from the original study rather than reassigned in the present analysis. Comparison across embeddings assesses the extent to which each integration approach preserves the original biological and clinicopathological structure of the dataset, including cellular identities and tissue-associated heterogeneity within metastatic breast cancer samples, including liver metastases.

**Figure S19:**
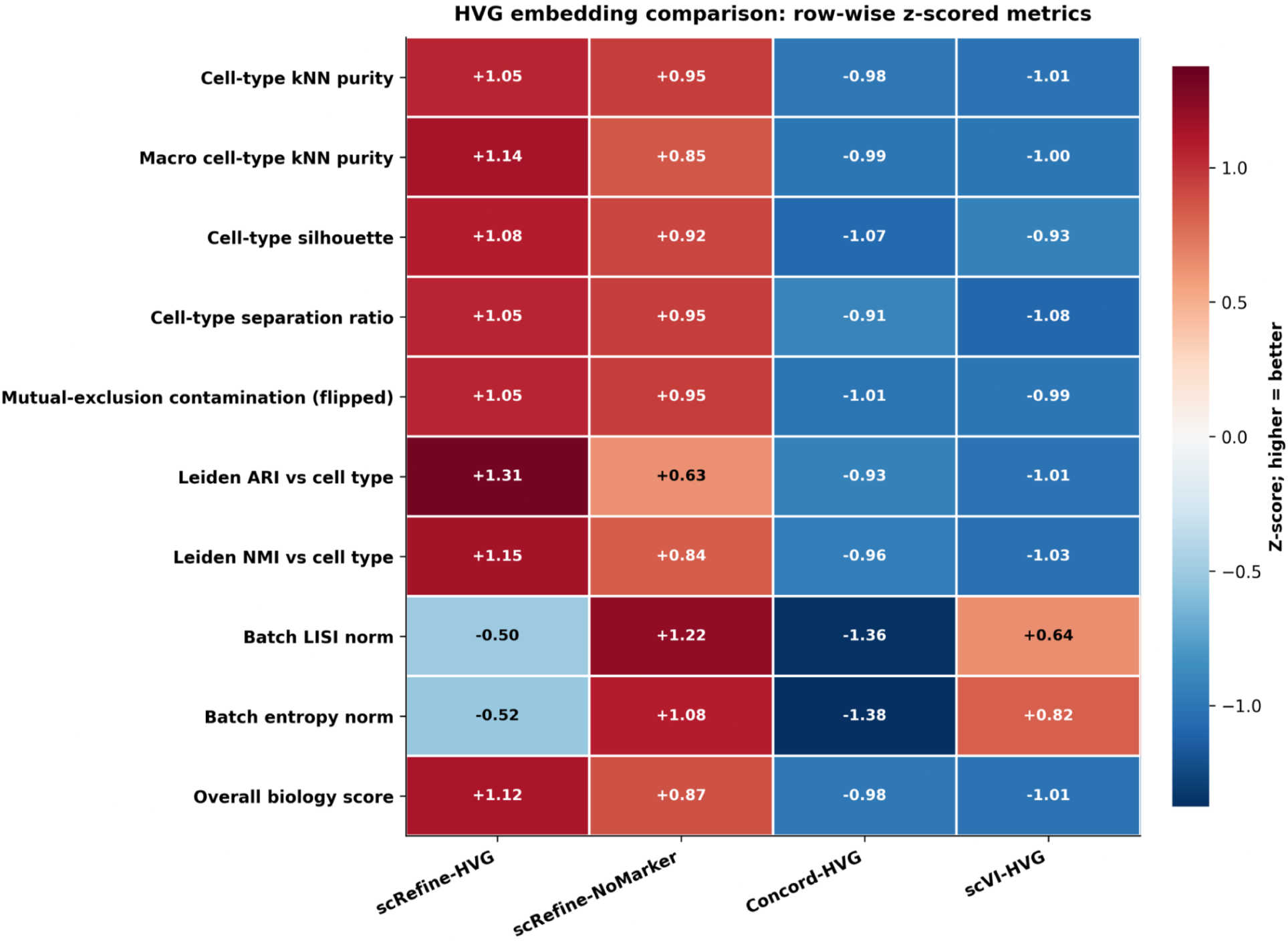
Quantitative benchmarking of biological preservation and batch integration in the metastatic breast cancer dataset. Heatmap comparing scRefine with highly variable gene (HVG) selection, scRefine without HVG selection, Concord, and scVI using standardized benchmarking metrics. Biological preservation was assessed using cell-type and macro cell-type k-nearest-neighbor (kNN) purity, cell-type silhouette score, cell-type separation ratio, mutual-exclusion contamination, and agreement between Leiden clusters and the original author-provided cell-type annotations using adjusted Rand index (ARI) and normalized mutual information (NMI). Batch integration was evaluated using normalized batch USI and batch entropy metrics. Values are shown as standardized z-scores, with higher scores indicating better relative performance. The overall biology score summarizes performance across biological preservation metrics. scRefine achieves the strongest overall preservation of the original cell-type structure, with the HVG-based implementation showing the highest overall biological score, while the comparison highlights the trade-off between biological structure preservation and batch mixing across integration methods.

**Figure S20:**
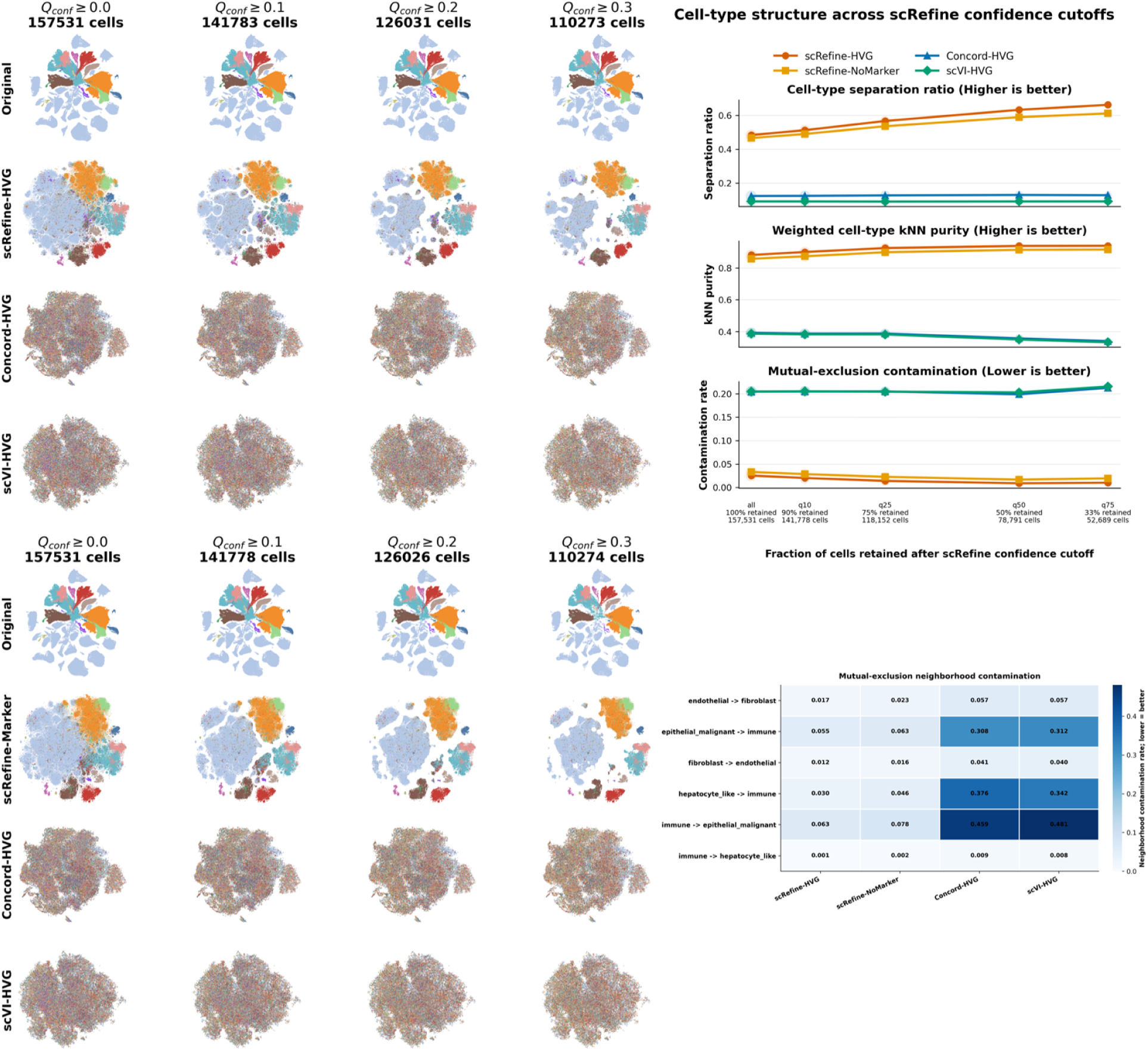
Effect of scRefine confidence-based cell filtering on preservation of cell-type structure across embeddings. The metastatic breast cancer dataset was progressively filtered using increasing scRefine confidence-score thresholds (Qconf≥0.0, 0.1, 0.2, and 0.3). At each threshold, the identical set of low-confidence cells identified by scRefine was removed from the original dataset and from all alternative embeddings, enabling direct comparison on exactly the same retained cells. Left panels show the author-provided cell-type annotations projected onto the original representation and embeddings generated using scRefine, Concord, and scVI after each confidence-based filtering step; the number of retained cells is indicated above each column. Right, quantitative benchmarking evaluates cell-type separation ratio, weighted cell-type kNN purity, and mutual-exclusion neighborhood contamination across increasingly stringent confidence cutoffs. The lower-right heatmap summarizes pairwise neighborhood contamination between biologically distinct cell compartments. Increasing scRefine confidence thresholds progressively enrich cells with clearer biological identity, with scRefine maintaining stronger cell-type separation and neighborhood purity and lower cross-lineage contamination than the alternative embeddings when evaluated on the same cell subsets.

